# Microbiomes change the rules of host evolution

**DOI:** 10.64898/2026.10.13.758714

**Authors:** Natalia L. Komarova, Margaret Y. Demmel, Jonathan B. Shurin, Dominik Wodarz

## Abstract

Microbiomes are pervasive determinants of host phenotype and fitness, yet their consequences for host population genetics remain poorly understood. Here we show that microbiome-associated population structure can fundamentally alter the evolutionary dynamics of host populations. We develop a general framework that maps hosts distributed among an arbitrary number of microbial states onto an effective population size, *N*_eff_, and effective selection coefficient, *s*_eff_, thereby reducing potentially high-dimensional host–microbiome dynamics to the quantities governing genetic drift and selection. Applying the framework to five empirically anchored host–microbe systems spanning ciliates, insects, and plants with bacterial, algal, and fungal associates reveals substantial departures from classical population-genetic predictions. Microbiome-associated structure can considerably alter both the effective strength of genetic drift and the strength of selection acting on host genetic variation, shifting populations into evolutionary regimes that differ markedly from those predicted from census size and intrinsic mutational effects alone. These changes can have profound consequences for evolutionary outcomes, including either suppression or enhancement of selection for both advantageous and disadvantageous mutations, sometimes reaching orders-of-magnitude differences. The effects are not determined simply by whether a microbial association is beneficial or detrimental, but by the ecological processes governing microbial acquisition, transmission, loss, and host demography. Our results provide a population-genetic extension of the holobiont perspective: when microbial associations generate distinct demographic states among hosts, they can alter the evolutionary forces acting on the host genome without requiring the host and microbiome to constitute a single hereditary unit.

## 1 Introduction

Evolution rarely occurs in ecological isolation. Animals, plants, and many unicellular eukaryotes live in association with microorganisms that colonize their surfaces, tissues, or intracellular compartments [1–4]. These microorganisms can influence nutrition, metabolism, development, immune defence, survival, and reproduction, and can therefore contribute substantially to host fitness [3, 5, 6]. For many organisms, the ecological environment in which host genomes evolve consequently includes a microbiome whose composition and effects vary among hosts, while individual hosts can transition between distinct microbial states.

A distinctive feature of this environment is that microbial and genetic inheritance need not coincide. The diversity of transmission mechanisms is illustrated by systems spanning almost complete environmental acquisition to high-fidelity vertical transmission and infection-like horizontal transmission. Juvenile squid recruit luminous bacterial symbionts from seawater, and broad-headed stinkbugs acquire beneficial *Burkholderia* from soil in each generation, whereas intracellular symbionts such as *Buchnera* are transmitted directly from aphid mothers to developing embryos [7–9]. Many microbiomes combine these routes, with incomplete parental transmission supplemented by environmental acquisition, as observed in strain-level mother-to-infant microbial transmission in humans [10]. Hosts and their descendants can therefore move among microbial states through acquisition, loss, and transmission, while the host genome follows a distinct path of inheritance.

This separation between genetic and microbial inheritance has potentially fundamental consequences for host evolution. Genetic drift and natural selection are the two principal forces determining the fate of genetic variation, and their effective strengths depend on the demographic structure through which lineages propagate [11–13]. In a microbiome-associated population, descendants carrying the same host genotype can occupy microbial states with different birth and death rates and can repeatedly move among those states. The microbiome can therefore alter the demographic and stochastic environment experienced by host genes even when the intrinsic effect of a host mutation itself is unchanged. Whether, and by how much, this changes the effective action of drift and selection is not captured by the fitness effect of the microbiome alone.

A substantial body of evolutionary theory has investigated host–microbiome associations from complementary perspectives. Early mathematical models examined the assembly and maintenance of beneficial microbial communities and host mechanisms that favor cooperative symbionts [14, 15]. Subsequent models have considered simultaneous selection on hosts and microbes, multilevel and holobiont selection, and the consequences of vertical and horizontal microbial transmission [16–19]. Other work has addressed the evolution of host control over microbial cooperation and the ecological and evolutionary processes governing microbial inheritance [20–23]. Complementary quantitative-genetic approaches ask how microbiome-mediated phenotypic variation and its transmission contribute to host responses to selection [24, 25]. Together with the broader holobiont and hologenome literature [26–29], these studies have established that microbial communities can participate in, and themselves be shaped by, evolutionary processes. Recent syntheses nevertheless emphasize that a general quantitative theory linking microbiome ecology to host evolutionary dynamics remains incomplete [4, 30]. The missing step is more fundamental than predicting the evolution of a particular host–microbe interaction. We lack a general population-genetic mapping from the ecological dynamics of microbial association to the basic evolutionary forces acting on the *host genome*. Given hosts distributed among microbial states with different demographic properties and connected by arbitrary processes of microbial acquisition, loss, and transmission, how are genetic drift and natural selection themselves changed? Answering this question requires translating potentially high-dimensional microbial-state dynamics into the effective population size and effective strength of selection experienced by host genetic variation. Such a mapping would make it possible to ask whether the presence of a microbiome changes the population-genetic regime in which the host evolves, rather than merely whether the microbiome changes host fitness.

Here we provide this mapping. We develop a general stochastic framework that derives the effective population size, *N*_eff_, and effective selection coefficient, *s*_eff_, directly from the demographic and transition processes connecting an arbitrary number of microbial states. These quantities characterize how microbiome-associated ecology modifies genetic drift and selection, with altered mutation fixation emerging as one consequence. Application to experimentally characterized host–microbe systems shows that these changes can be large under biologically relevant conditions. Our results therefore identify a previously unresolved population-genetic consequence of microbial association: microbiomes can change the effective rules by which host genomes evolve.

## 2 Results

Microbiomes of host organisms typically contain multiple microbial taxa, and hosts may consequently occupy many microbial states that differ in composition, function, and their effects on host demography [6, 31]. Transitions among these states can result from environmental microbial acquisition [8], horizontal and vertical transmission [32], and loss [33]. The resulting ecological models can therefore contain many host states and transition processes that are specific to the host system under consideration. A general population-genetic treatment should ideally accommodate ecological complexity without requiring exhaustive evolutionary simulations for each particular model. Here we develop such a framework. We consider hosts distributed among an arbitrary number of microbial states and describe their ecology in terms of demographic and transition events that change the numbers of hosts occupying these states. From this event structure, we derive the effective population size, *N*_eff_, and effective selection coefficient, *s*_eff_, which together determine the fixation probability of a rare host mutation under weak selection. We show that the population structure brought about by microbial associations can substantially alter these evolutionary measures compared to predictions for hosts evolving in isolation.

We start out by describing this approach mathematically in a general system containing *k* microbial states and their stochastic ecology. Our construction combines reproductive-value weighting of a structured population [34, 35] with a one-dimensional diffusion and fast-mode reduction [36, 37].

While this approach can in principle be applied to biological systems of arbitrary complexity, it is currently not possible to test the relevance of the mathematical findings against such biological systems due to the absence of parameter measurements. Estimates of relevant parameters can, however, be derived from experiments in a simpler setting, where only two host states are distinguished: individuals that are associated with a particular microbe and those that are not. Even in this minimal setting, the underlying ecology can take different forms. Microbes may be transmitted horizontally between hosts or acquired from an environmental reservoir, may be inherited with varying fidelity during host reproduction, and may alter host birth or death rates. Population regulation can likewise act through density-dependent reproduction, mortality, or both.

Hence, we will use our general framework to study an example system characterized by the association of a host with one specific microbe. We apply it to a number of different experimentally studied interactions where model parameterization can be carried out. We demonstrate that even for such relatively simple systems, microbe associations can substantially change the evolutionary dynamics of the host species in natural systems.

One of these biological systems allows partial model parameterization in a more complex case containing multiple microbial associations, although with higher parameter uncertainty (see SI Section 4). The application of our methodology to higher levels of complexity will require the experimental estimation of demographic parameters in host species characterized by multiple microbial associations and states. Although such measurements are not yet available, our framework identifies the required data and provides a direct route for translating future measurements of multistate host–microbe demography into quantitative evolutionary predictions.

### 2.1 Universal method of calculating *N*_eff_ and *s*_eff_

We represent hosts in *k* microbial states and their stochastic ecology by *R* events. We are interested in the fate of a host variant introduced into the population, given that its rates are slightly different from those of the wild-type. The selection coefficient *s*, with |*s*| ≪ 1, parameterizes a perturbation of the mutant event rates about neutrality; for a summary of notations please see Table 1 in SI Section 1.

**Table 1:** Effects of newborn allocation (columns) and microbiome type (rows) on effective population size (a) and effective selection coefficient (b). For a deleterious microbiome, the threshold separating ℛ_*s*_ *<* 1 and 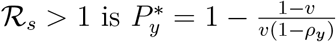.

| Microbiome type \ Newborn allocation | $P_y^{\text{de novo}} > P_y^{\text{census}}$ | $P_y^{\text{de novo}} < P_y^{\text{census}}$ |
| --- | --- | --- |
| Beneficial | $\mathcal{R}_N < 1$ | $\mathcal{R}_N > 1$ |
| Deleterious | $\mathcal{R}_N > 1$ | $\mathcal{R}_N < 1$ |

Table 1: Effects of newborn allocation (columns) and microbiome type (rows) on effective population size (a) and effective selection coefficient (b). For a deleterious microbiome, the threshold separating $\mathcal{R}_s < 1$ and $\mathcal{R}_s > 1$ is $P_y^* = 1 - \frac{1-v}{v(1-\rho_y)}$ .
| Microbiome type \ Newborn allocation | $P_y^{\text{de novo}} > P_y^*$ | $P_y^{\text{de novo}} < P_y^*$ |
| --- | --- | --- |
| Beneficial | Not attainable | $\mathcal{R}_s < 1$ |
| Deleterious | $\mathcal{R}_s > 1$ | $\mathcal{R}_s < 1$ |

The goal of our methodology is to determine whether and to what extent the compartmental structure induces changes in the evolutionary properties of a mutant host. In particular, we will determine if the probability of fixation of a mutant host is different compared to an unstructured reference, given by the Moran-type fixation function for the census size *N* and intrinsic selection coefficient *s*:

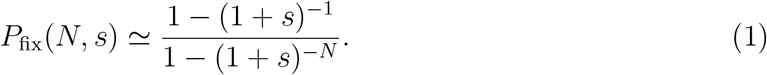

We will proceed by calculating the effective population size, *N*_eff_, and the effective selection coefficient, *s*_eff_. The corresponding approximation for the structured population is *P*_fix_(*N*_eff_, *s*_eff_).

To this end, we are interested in the regions of the parameter space where stable coexistence of the different compartments is observed; we refer to this stable solution as ***N*** = (*N*_1_, …, *N*_*k*_)^T^ and the total equilibrium population size as 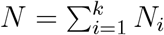. We focus on the near-equilibrium dynamics of rare mutants with selection coefficient (positive or negative) *s*.

Let ***M*** (*t*) denote the random vector of mutant abundances and let ***m*** = (*m*_1_, …, *m*_*k*_)^T^ denote one of its possible states. When the mutant is rare, the rate of event *r* can be linearized as

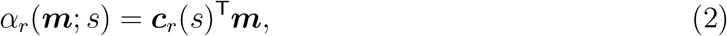

where *c*_*r,i*_(*s*) is the per-capita contribution of a mutant in compartment *i* to the rate of event *r*. The coefficients **c**_*r*_(*s*) are evaluated at the resident equilibrium ***N***; their dependence on ***N*** and on any equilibrium environmental variables is suppressed for notational simplicity. The expression **c**_*r*_(*s*)^T^**m** is the leading-order event rate when the mutant is rare. The biological way in which the mutation affects reproduction, mortality, or transitions is encoded in the derivatives of **c**_*r*_(*s*) at *s* = 0.

Define the mean matrix

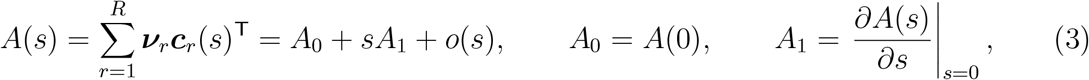

where the jump vector, ***ν***_*r*_, lists the change in the mutant count in each compartment that results from process *r*. Writing 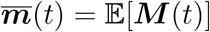, the mean mutant vector satisfies

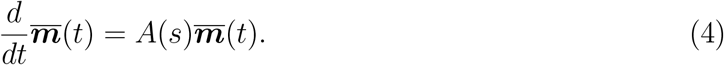

At neutrality, we assume that *A*_0_ has a simple zero eigenvalue, with positive right and left null vectors ***N*** and ***w***:

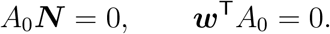

We further assume that all remaining eigenvalues have negative real parts, so that perturbations of the mutant composition relax toward the resident composition. The component *w*_*i*_ is the relative reproductive value of a mutant in compartment *i*.

Define vector ***P*** = (*P*_1_, …, *P*_*k*_)^T^, where *P*_*i*_ is the probability that the initial mutant is introduced into microbial state *i*. For de novo mutation, **P** is determined by the distribution of newborn production among the states. It is generally not the census distribution *N*_*i*_*/N*. Other ways of mutant introduction can be incorporated by choosing an appropriate **P**.

Then the effective parameters are given by

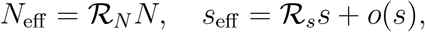

where

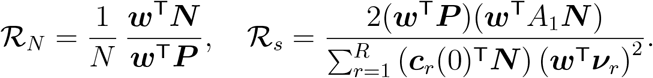

Details of the derivation are provided in SI Section 1.

The two modifiers combine in the effective approximation to the fixation probability. Relative to the classical prediction based on *N* and *s*, the fold change in fixation probability is, for |*s*| ≪ 1 but arbitrary *N* |*s*|, can be written as

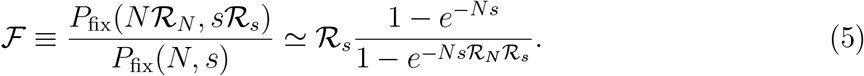

The quantity specifies in which way the presence of structure changes the probability of mutant fixation.

### 2.2 The two-state host–microbe system as a minimal example

We first illustrate the general framework with a minimal system containing two host states: microbiome-unassociated individuals (*x*) and microbiome-associated individuals (*y*), see figure 1(a). Both states reproduce and die. Unassociated hosts acquire the microbiome horizontally at rate *βxy*, whereas an associated parent transmits the microbiome to its offspring with probability *v*, which we call the *vertical transmission fidelity*. With probability 1 − *v*, its offspring is born without the microbiome. Thus, *v* = 0 corresponds to no vertical transmission and *v* = 1 to perfect vertical transmission. This ecological structure is related to susceptible– infected models incorporating both horizontal and imperfect vertical transmission [38–40] (see below for other model forms with environmental uptake rather than horizontal transmission).

**Figure 1:**
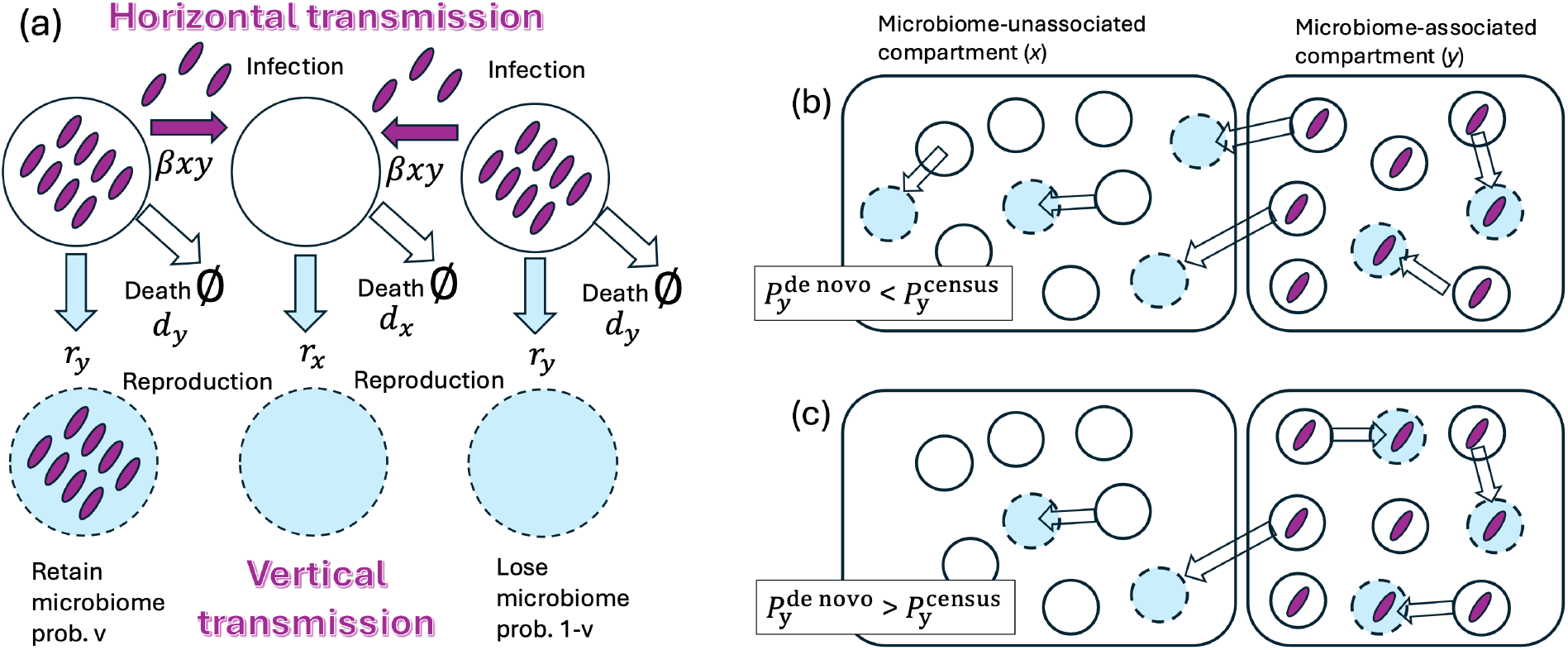
Demographic and transmission aspects of the two-state system. (a) Microbiome-unassociated hosts (*x*) can acquire the microbiome horizontally, whereas microbiome-associated hosts (*y*) can transmit it vertically during reproduction. The microbiome is shown by purple elongated shapes and newborn individuals by blue dashed circles. (b,c) Illustration of the distinction between the census fraction, 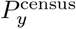, and the de novo fraction, 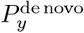, which gives the fraction of newborns, and hence new mutations, entering the *y* state. In (b), 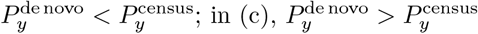.

Both host states reproduce under density regulation with carrying capacity *K*, with intrinsic reproduction rates *r*_*x*_ and *r*_*y*_, such that the per-capita reproduction rates are 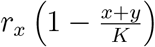 and 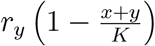 in the two states. Individuals in the two states die at rates *d*_*x*_ and *d*_*y*_, respectively. Denote their equilibrium abundances by *N*_*x*_ and *N*_*y*_. We assume that a host mutation changes the intrinsic reproduction rate by the same proportional amount in both microbial states,

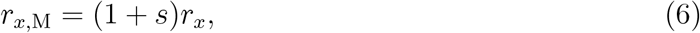

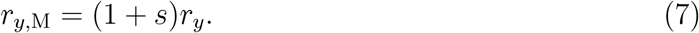

The parameter *s* is therefore the mutation’s intrinsic selective effect before accounting for movement of its descendants between microbial states.

We tested the analytical framework described in the previous section (see also SI Section 2) against direct Gillespie simulations [41]. Each realization began with the resident population at the equilibrium (*N*_*x*_, *N*_*y*_), and the first reproduction event generating a mutation introduced a single mutant into either the *x* or *y* state. The mutant lineage was then followed until extinction or fixation. Figure 2 compares these fixation probabilities with the classical prediction based only on census population size *N* = *N*_*x*_ + *N*_*y*_ and intrinsic selection coefficient *s*. The classical prediction can differ from the simulations by orders of magnitude, whereas replacing *N* and *s* by the effective quantities *N*_eff_ and *s*_eff_ derived in section 2.1 closely reproduces the stochastic results for the parameter sets examined.

**Figure 2:**
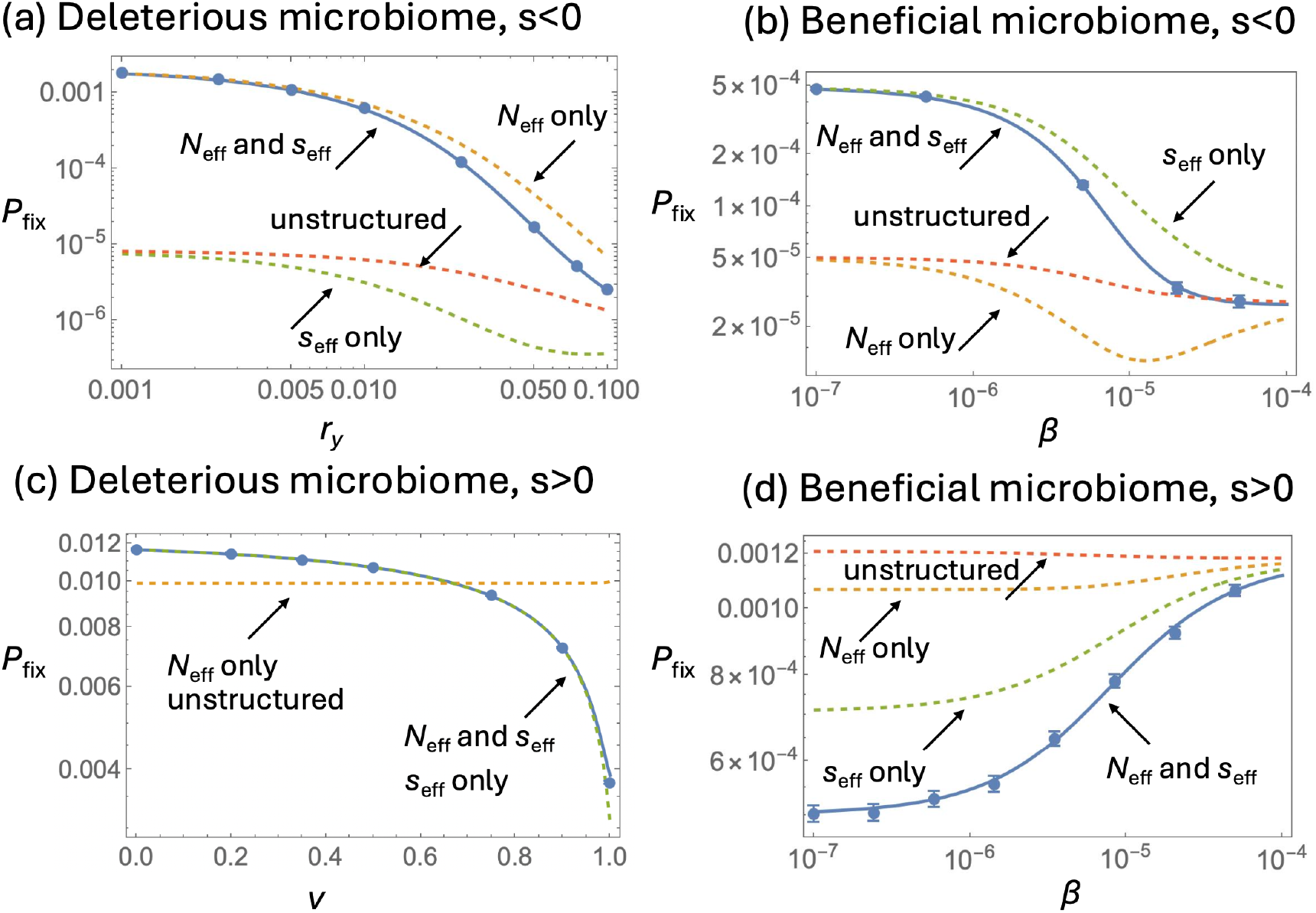
Numerically estimated fixation probabilities and theoretical predictions. Blue symbols represent stochastic simulations (at least 10^7^ runs per point; some 95% confidence intervals are too small to see). The dashed red line (unstructured) is *P*_fix_(*N, s*), equation (1). The dashed yellow and green lines are *P*_fix_(*N*_eff_, *s*) and *P*_fix_(*N, s*_eff_), respectively, and the solid blue line is *P*_fix_(*N*_eff_, *s*_eff_). (a) Deleterious microbiome, disadvantageous mutant (*s* = −0.005), with *r*_*x*_ = 0.3, *v* = 1, and *β* = 0.00008, varying *r*_*y*_ *< r*_*x*_. (b) Beneficial microbiome, disadvantageous mutant (*s* = −0.0025), with *r*_*x*_ = 0.075, *r*_*y*_ = 0.3, and *v* = 0.35, varying *β*. (c) Deleterious microbiome, advantageous mutant (*s* = 0.01), with *r*_*x*_ = 0.3, *r*_*y*_ = 0.075, and *β* = 0.00003, varying *v*. (d) Beneficial microbiome, advantageous mutant (*s* = 0.001), with *r*_*x*_ = 0.075, *r*_*y*_ = 0.3, and *v* = 0.35, varying *β*. The remaining parameters are *d*_*x*_ = 0.02, *d*_*y*_ = 0.01, and *K* = 1936.

### 2.3 Ecological complexity collapses onto effective drift and selection

Although this two-state model contains seven demographic and transmission parameters, their evolutionary consequences can be organized in terms of a small number of composite quantities. The first distinction is between the composition of the standing population and the distribution of newly produced individuals. The census fraction of associated hosts is

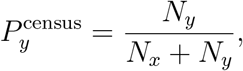

whereas

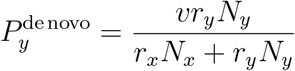

is the fraction of newborns entering the associated state. Because de novo mutations arise during reproduction, 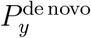 is also the probability that a new mutation initially occurs in state *y*. Thus, 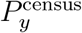 describes where individuals are, whereas 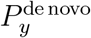 describes where new lineages enter the population.

The two states can also differ in their expected long-term genetic contribution. Taking the reproductive value of an *x* host to be one, the relative reproductive value of a *y* host is

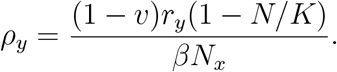

The demographic effect of microbiome association can be summarized by *B*_*y*_ = *r*_*y*_*d*_*x*_ − *r*_*x*_*d*_*y*_, with _*y*_ *>* 0 denoting a beneficial association and _*y*_ *<* 0 a deleterious one. At a positive coexistence equilibrium with *β >* 0,

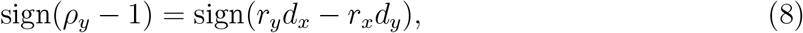

so beneficial and deleterious associated states have, respectively, higher and lower reproductive value than the unassociated state.

To express the effective parameters compactly, let

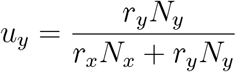

be the fraction of reproduction events occurring in state *y*. Because a fraction *v* of the offspring produced in state *y* retain the microbiome, 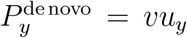. We normalize the reproductive values as *w*_*x*_ = 1 and *w*_*y*_ = *ρ*_*y*_. The mean reproductive value of a uniformly sampled census individual is then

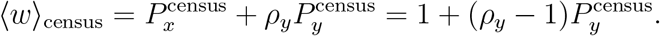

Similarly, the mean reproductive value of a newly arising mutant is

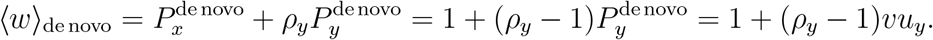

The effective population-size modifier can therefore be written as

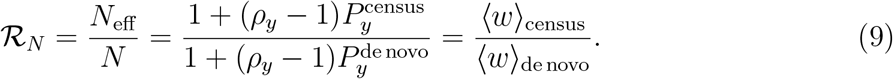

The selection modifier also depends on the reproductive values of parents and their offspring. Let *w*_p_ and *w*_n_ denote the reproductive values of the parent and newborn, respectively, in a randomly sampled reproduction event. A reproduction event in state *x* has *w*_p_ = *w*_n_ = 1. For a parent in state *y, w*_p_ = *ρ*_*y*_, while the offspring has reproductive value *ρ*_*y*_ with probability *v* and reproductive value 1 with probability 1 − *v*. Consequently,

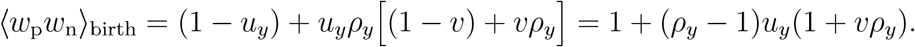

Under weak selection,

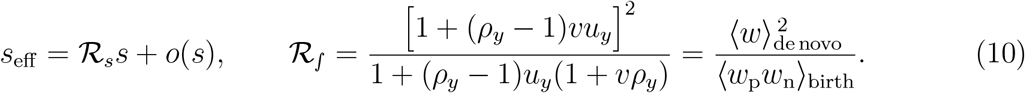

Thus the numerous ecological parameters act on host evolution through two effective modifiers: ℛ_*N*_, which changes the strength of genetic drift, and ℛ_*s*_, which changes the magnitude of selection.

### 2.4 Effective population size follows a universal rule

Equation (9) presents the modifier of the population size as a ratio between the mean reproductive value of the census population and the mean reproductive value of the newborn individuals. This is a general result that holds for any number of compartments and for any transmission rules. Equation (9) further gives a simple rule for the direction in which microbiome structure changes effective population size:

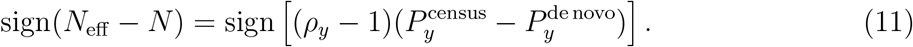

Thus, only two features determine whether drift is strengthened or weakened: which microbial state has greater reproductive value, and whether that state is over- or underrepresented among newly arising mutations relative to the standing population.

For a beneficial microbiome (*ρ*_*y*_ *>* 1), 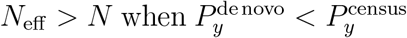 and *N*_eff_ *< N* when 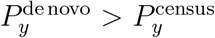; for a deleterious microbiome (*ρ*_*y*_ *<* 1), these relationships reverse (table 1(a)). For example, if the associated state has high reproductive value but is underrepresented among newborn mutants, a typical new mutant has lower reproductive value than a typical member of the resident population. Its neutral fixation probability is therefore below 1*/N*, corresponding to *N*_eff_ *> N*. The effective population size is consequently determined not by census size alone, but by the interaction between reproductive value and the states in which new mutations arise.

Figure 3(a) illustrates this rule across demographic parameter space. The sign of ℛ_*N*_ – 1 changes upon crossing either the boundary between beneficial and deleterious association or the boundary 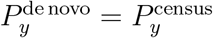.

**Figure 3:**
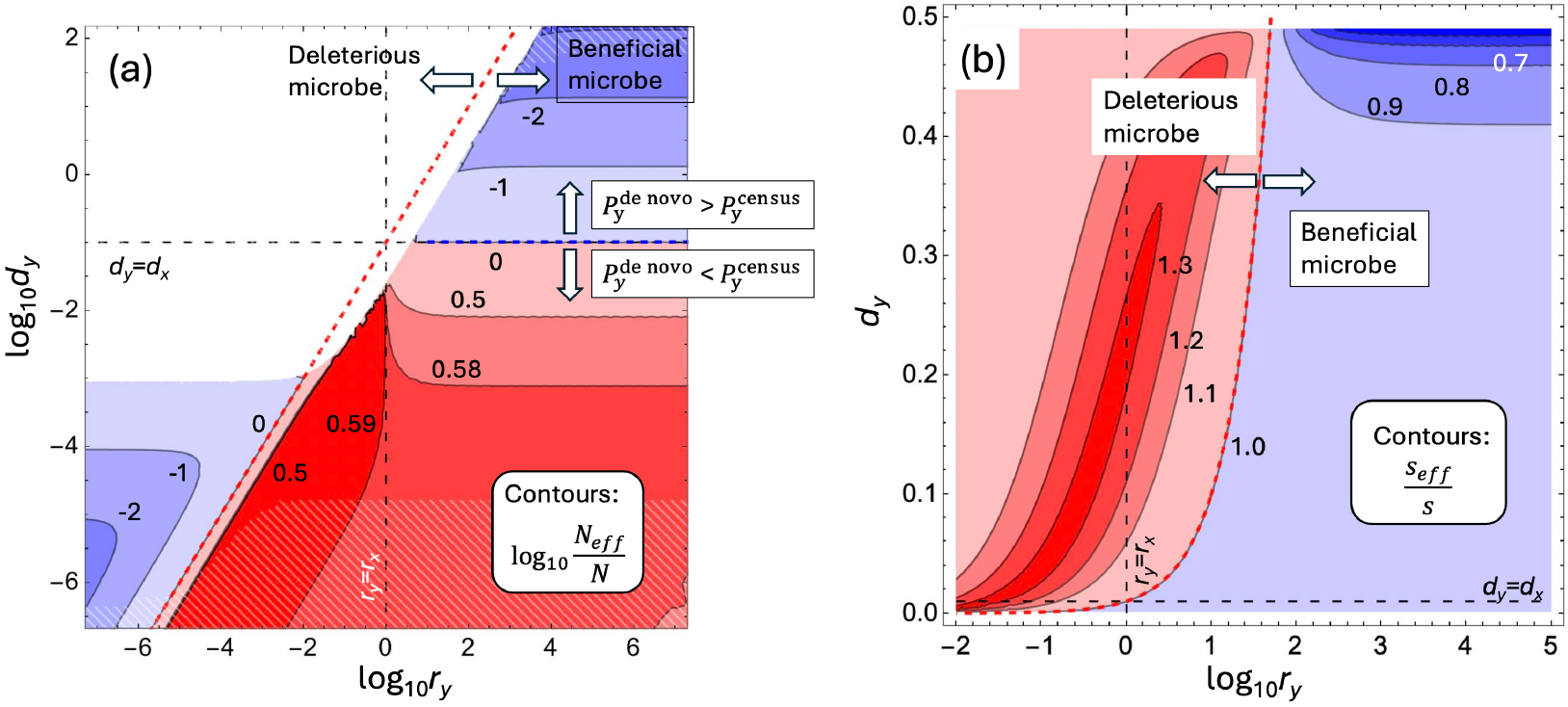
Effects of microbiome association on effective drift and selection. (a) log_10_(*N*_eff_*/N*) as a function of the reproduction and death rates of microbiome-associated hosts, log_10_ *r*_*y*_ and log_10_ *d*_*y*_. Blue regions have *N*_eff_ *< N* and red regions *N*_eff_ *> N*. White regions lack a stable coexistence equilibrium; white stripes indicate parameter regions in which either equilibrium population falls below 100. Parameters are *K* = 10^5^, *r*_*x*_ = 1, *d*_*x*_ = 0.1, *v* = 0.25, and *β* = 10^*−*8^. (b) The selection modifier ℛ_*s*_ = *s*_eff_*/s* as a function of log_10_ *r*_*y*_ and *d*_*y*_. Blue regions have ℛ_*s*_ *<* 1 and red regions ℛ_*s*_ *>* 1. Parameters are *K* = 10^5^, *r*_*x*_ = 1, *d*_*x*_ = 0.1, *v* = 0, and *β* = 5 × 10^*−*6^. In both panels, the red dashed line separates beneficial from deleterious microbiomes. In panel (a), the blue dashed line denotes 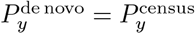.

### 2.5 Microbial transmission modifies the effective strength of selection

The effect on selection is asymmetric. A beneficial microbiome always gives ℛ_*s*_ *<* 1, suppressing the magnitude of selection on both advantageous and deleterious host mutations.

A deleterious microbiome can either increase or decrease the effective selection coefficient (table 1(b)). The boundary,

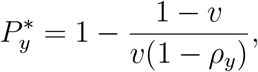

exists if 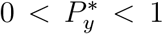, with amplification when 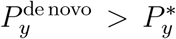 and suppression otherwise. Amplification therefore occurs when new mutations are sufficiently concentrated in the low-reproductive-value associated state; under imperfect vertical transmission (*v <* 1), mutants born in the deleterious compartment can escape to the other compartment of increased reproductive value, which amplifies the difference imposed by *s*. Even then the effect is bounded: for a deleterious microbiome, ℛ_*s*_ ≤ 4*/*3. Figure 3(b) illustrates the resulting selection regimes.

Horizontal and vertical transmission influence evolution by changing both the distribution of newborn mutants and their subsequent movement between microbial states. Their effects are therefore not generally monotonic: increasing *β* or *v* can move a population through different regions of the reduced parameter space. The limiting cases provide useful reference points. With purely horizontal transmission (*v* = 0), all mutants are born unassociated, so 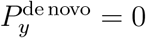, and

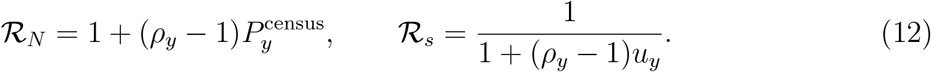

A beneficial microbiome then gives ℛ_*N*_ *>* 1 and ℛ_*s*_ *<* 1, whereas a deleterious microbiome gives ℛ_*N*_ *<* 1 and ℛ_*s*_ *>* 1. With purely vertical transmission (*β* = 0), stable coexistence in this model is possible only for a beneficial microbiome; selection is suppressed, while ℛ_*N*_ *<* 1 if *d*_*y*_ *> d*_*x*_ and ℛ_*N*_ *>* 1 if *d*_*y*_ *< d*_*x*_. Intermediate mixtures of horizontal and vertical transmission are best understood as trajectories through the reduced quantities rather than as interpolations between these endpoints, see SI Section 2.

As figure 2 shows, accounting for either *N*_eff_ or *s*_eff_ alone is generally insufficient; both are required to reproduce the stochastic fixation probabilities. The relative importance of the two modifiers depends on the scaled intrinsic selection strength *N*|*s*|. Near neutrality, *N*|*s*| ≪ 1, ℱ ≃ 1*/* ℛ_*N*_, so fixation is governed primarily by effective population size. For strongly advantageous mutations, *Ns* ≫ 1, ℱ ≃ ℛ_*s*_, and the effective strength of selection becomes dominant. At intermediate selection strengths both modifiers contribute, while for deleterious mutations the product ℛ_*N*_ ℛ_*s*_ becomes increasingly important. Thus, the evolutionary consequence of microbiome association cannot in general be inferred from its effect on drift or selection separately.

### 2.6 Application to empirical host–microbe systems

We next asked whether microbiome-associated population structure can substantially alter host evolution in empirically characterized biological systems. We considered five examples spanning unicellular and multicellular hosts and bacterial, algal, and fungal associates (SI Section 3). These systems differ not only in whether microbial association is beneficial or deleterious, but also in how the microbe is acquired and transmitted and in how host population size is regulated.

The examples below provide an opportunity to ask to what extent the effects of microbe associations on host evolutionary dynamics described mathematically above are observed in biologically realistic parameter regimes. They further allow us to explore how variation in demographic parameters in realistic ranges can change host evolutionary measures. We find that microbe associations result in substantial changes in host evolutionary dynamics across different biological systems, and that details of the population dynamics underlying these systems can alter this effect. This is described as follows.

#### *Paramecium caudatum*–*Holospora undulata*: a fully parameterized example

The clearest direct application is provided by the bacterial parasite *Holospora undulata* in the ciliate *Paramecium caudatum. Holospora* is transmitted vertically during host division and horizontally through infectious forms released from infected cells. Importantly, Nørgaard *et al*. fitted a continuous-time epidemiological model, closely related to the two-state model considered here, directly to longitudinal densities of infected and uninfected hosts [42]. The resulting posterior distributions therefore provide the demographic and transmission parameters needed for our calculation without requiring additional assumptions about unmeasured rates.

The experiment is particularly informative because parasite populations had been evolved either at the expanding front of a spatial range expansion (Front/UP) or in the range core (Core/DOWN). The resulting parasite types differed strongly in their life-history properties: Core/DOWN parasites were more virulent and had higher horizontal transmission than Front/UP parasites. Propagating the complete posterior parameter distributions through our framework shows that these differences also change the population-genetic environment experienced by host mutations. For Front/UP parasites, the posterior median modifiers are

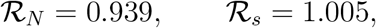

so the evolutionary dynamics remain close to those expected from census population size and intrinsic selection alone. In contrast, Core/DOWN parasites give

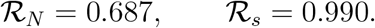

Thus, the principal effect of the Core/DOWN parameter combination is not a substantial change in the effective selection coefficient, but a reduction of the effective population size to about 69% of census size. Microbial association therefore increases the strength of genetic drift while leaving the deterministic strength of selection nearly unchanged.

This change in drift is sufficient to alter fixation appreciably. For example, for a host mutation with *Ns* = 1, the median fixation probability is only about 3% above the classical expectation in the Front/UP treatment, but approximately 26% higher in the Core/DOWN treatment. The effect is considerably larger for deleterious mutations, whose classical fixation probabilities can be small and become orders of magnitude larger when effective population size is reduced.

The *Paramecium* example shows that changed evolutionary dynamics described mathematically above can indeed occur in real biological systems under estimated parameter values. Further, this example shows that changes in the microbial parasite itself (changes in kinetics such as in the Core/DOWN and Front/UP scenarios) can alter the subsequent evolutionary dynamics of the host, even when the intrinsic effect of the host mutation is unchanged.

#### *Drosophila melanogaster* : transmission determines how microbial effects on host demography translate into evolution

The *Drosophila* system provides an example in which the demographic effects of several individual bacterial species have been measured, but their natural uptake and loss rates are less well characterized. In contrast to the *Paramecium* example, microbes are acquired by environmental uptake rather than horizontal transmission, requiring an adjustment of the mathematical model, see SI Section 3. Gould *et al*. compared germ-free flies with flies associated individually with five bacterial species and measured lifetime reproduction and survival [6]. Under our demographic criterion, two of these associations are deleterious and three are beneficial. We used these data to parameterize the corresponding host birth and death rates and then varied the rates of environmental acquisition and vertical transmission over broad ranges.

The results show that the fitness effect of a bacterium on its host does not, by itself, determine its effect on host evolution. The same demographic association can have almost no evolutionary effect under one transmission regime and a substantial effect under another. Beneficial bacteria generally suppress the effective strength of selection, but the magnitude of this effect depends strongly on the balance between environmental acquisition and vertical transmission. For deleterious bacteria, both the magnitude and, in some parameter regions, the direction of the change in fixation can depend on transmission. Across the explored parameter space, these effects can be large: for some beneficial associations, deleterious host mutations become up to approximately 10^3^ times more likely to fix than predicted for an unstructured population, whereas the fixation probability of advantageous mutations can be reduced by as much as 80%. These ranges should be interpreted as a sensitivity analysis rather than point predictions, because the transmission parameters were not measured in the source experiment. They identify transmission itself as a key biological quantity required to translate measured microbiome effects on host fitness into evolutionary predictions.

#### *Tetrahymena utriculariae*–*Micractinium tetrahymenae*: identical population growth can imply different evolutionary dynamics

The *Tetrahymena*–*Micractinium* system reveals a different source of uncertainty with an important general implication. The algal symbiont can be transmitted during host division and can also be reacquired from the environment. Under the oxic, bacteria-fed conditions used for our parameterization, symbiotic ciliates grow substantially more slowly than aposymbiotic ciliates [43, 44]. Population-density measurements, however, determine net population growth much more directly than they determine birth and death separately. Evidence from related *Tetrahymena* indicates that density regulation can act through reduced division, increased mortality, or both [45].

We therefore constructed the model so that the same deterministic net growth rate can be generated with density dependence acting entirely through birth, entirely through death, or through a mixture of the two (see SI Section 3). These alternatives are indistinguishable from the corresponding single-population density trajectories, but they are not evolutionarily equivalent. When density regulation acts primarily through mortality, the microbiome has only a modest effect on ℛ_*N*_ and ℛ_*s*_. In contrast, when density regulation acts through reproduction, the same measured net population dynamics can produce ℛ_*N*_ values as low as approximately 0.4 and ℛ_*s*_ values ranging from about 0.35 to 1.2, depending on vertical transmission and baseline mortality. The corresponding fixation effects can exceed three orders of magnitude for deleterious mutations, while advantageous mutations can be either substantially suppressed or, in other parameter regions, moderately enhanced.

This example illustrates a broader point. Details of the demographic processes underlying the particular biological system can have a major influence on the evolutionary dynamics, even if no difference is observed in the overall growth dynamics. How density regulation is implemented mechanistically (through reproduction, mortality, or both) can be as important as the presence of the microbiome itself.

#### *Aphis fabae*–*Hamiltonella defensa*: ecological context changes both microbial fitness effects and host evolution

The defensive endosymbiont *Hamiltonella defensa* illustrates how a third trophic interaction can reshape microbiome-mediated host evolution. In the absence of parasitoids, carrying *H. defensa* imposes a demographic cost on aphids, whereas the symbiont can strongly protect its host against parasitoid-induced mortality [46–49]. We therefore varied parasitoid attack rate together with horizontal transmission of the symbiont, while using experimentally informed values for host demography, maternal transmission, and protection.

Increasing parasitoid pressure changes the association from deleterious to beneficial because mortality rises much more rapidly in unassociated aphids than in symbiont-bearing aphids. For a range of attack rates where the endosymbiont is mildly beneficial, high-fidelity vertical transmission becomes sufficient to maintain the symbiont without substantial horizontal transmission. Near this boundary, the effect on effective population size remains relatively small, whereas the effective selection coefficient can become very strongly reduced. Consequently, microbiome-mediated suppression of selection can be pronounced: in the parameter regimes examined, deleterious host mutations can become roughly two orders of magnitude more likely to fix, while fixation of advantageous mutations can be reduced by approximately 60%. It is interesting that the strongest evolutionary effect (a reduction of ℛ_*s*_) is observed for the parameter range where the microbe is mildly beneficial, and decreases when it becomes more strongly beneficial.

Furthermore, this example illustrates a case where the microbial association can be “beneficial” or “deleterious” depending on the environment (the rate of attacks), and this ecological change can alter the strength with which selection acts on unrelated host mutations. The evolutionary effect of the microbiome is consequently determined not only by host–microbe interactions, but also by ecological processes acting on the host–microbe system.

#### *Agrostis hyemalis*–*Epichloë amarillans*: evolutionary quantities can remain identifiable despite incomplete parameterization

Finally, the association between winter bentgrass *Agrostis hyemalis* and the fungal endophyte *Epichloë amarillans* provides a predominantly vertically transmitted plant–microbe example. Field experiments measured reproductive components, vertical-transmission fidelity, and equilibrium endophyte prevalence under ambient and elevated precipitation [50]. Several continuous-time rates required by the ecological model, including mortality and horizontal acquisition, are not separately identified. Nevertheless, the general expressions reveal that, under the empirically motivated assumption of equal mortality in associated and unassociated plants, these unknown rates cancel from both ℛ_*N*_ and ℛ_*s*_ once the observed equilibrium prevalence, relative reproduction rate, and vertical-transmission fidelity are specified. The resulting estimates are

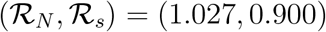

under ambient precipitation and

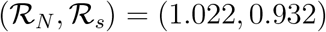

under elevated precipitation. Thus, the effect on genetic drift is small, whereas selection on host mutations is moderately weakened. The difference between precipitation treatments further illustrates how environmental conditions can modify host evolution indirectly by changing the demographic and transmission properties of the microbial association. More generally, this example shows that incomplete knowledge of every ecological rate does not necessarily prevent evolutionary inference: equilibrium constraints can sometimes identify the combinations of parameters that determine the evolutionary modifiers ℛ_*N*_ and ℛ_*s*_ even when individual rates remain unknown.

### Summary

These empirical examples reveal that there is no single “microbiome effect” on host evolution. A microbe’s direct effect on host birth and survival is only one component. The distribution of microbes among newborns, the balance between vertical and horizontal or environmental acquisition, the demographic processes through which population regulation acts, and interactions with the wider ecological community can all alter the reproductive values of host states and hence the effective action of drift and selection. In some systems the dominant effect is a change in *N*_eff_, as in the Core/DOWN *Paramecium* treatment; in others it is primarily a change in *s*_eff_, as in *Agrostis* and parts of the aphid parameter space; and in still others both quantities change substantially, see figure 4. The same framework can accommodate all of these cases.

**Figure 4:**
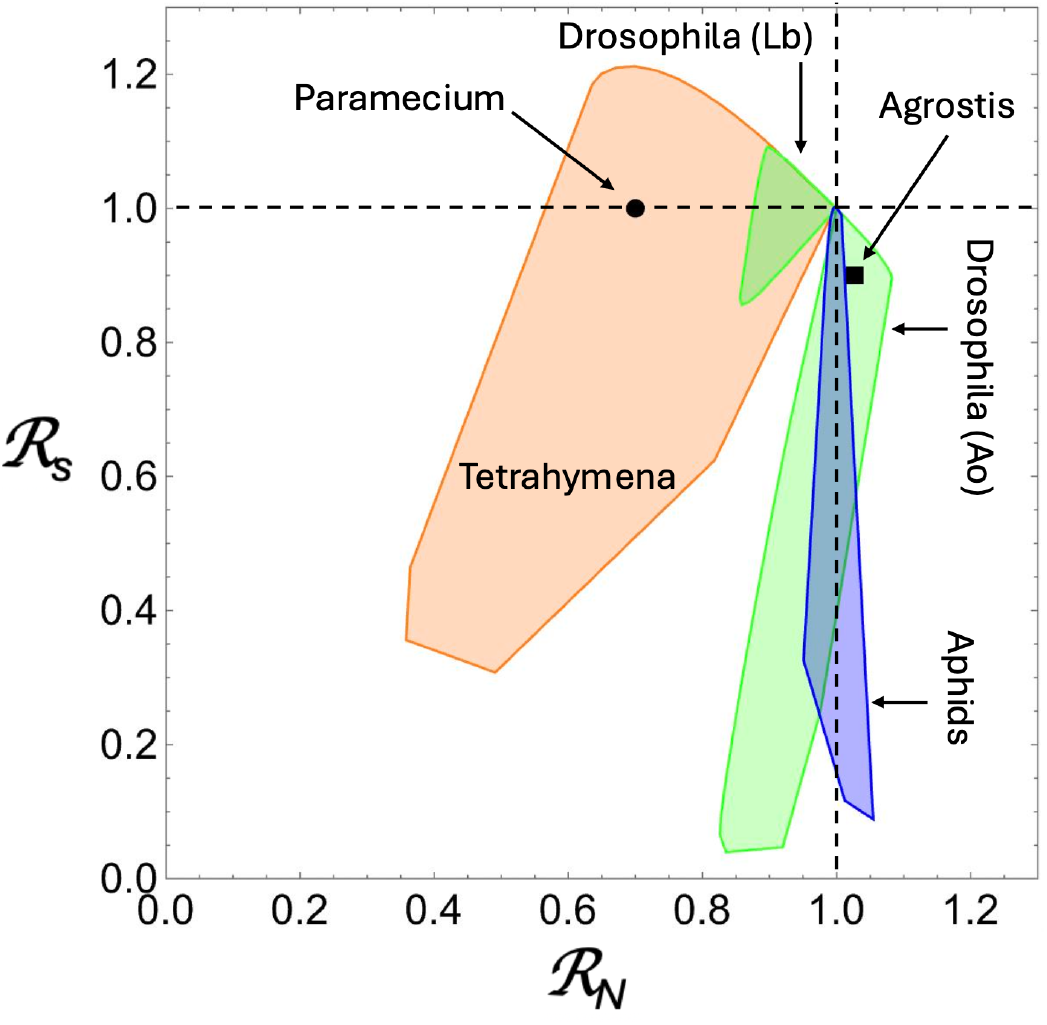
The five biological examples: effects of microbiome association on effective drift and selection. The values of the modifiers ℛ_*N*_ and ℛ_*s*_ are shown for each of the example systems, as we fix the measured parameters to their central values and vary parameters that could not be measured. This is the summary of calculation results presented in SI Section 3. Note that for Paramecium and Agrostis, we were able to provide a single central value, thus their regions are points. Only results for one deleterious (Lb) and one beneficial (Ao) microbe are shown for the Drosophila example.

The general framework is not restricted to binary host–microbe associations and can be applied to systems with multiple microbial states, as outlined in Section 2.1. To illustrate this for a specific biological example, we considered a four-state extension of the *Drosophila melanogaster* system in which hosts carry neither of two bacterial species, either species alone, or both simultaneously (SI Section 4). Demographic rates for the germ-free and single-species states were anchored in empirical measurements, whereas the less well characterized doubly colonized state was explored across a biologically plausible range. Despite the resulting increase in ecological complexity and parameter uncertainty, the same general calculation yields *N*_eff_, *s*_eff_, and fixation probability, and microbiome-associated structure continues to produce substantial changes in host evolutionary dynamics. The principal limitation in applying this framework to still more complex microbial communities is currently the availability of state-specific demographic and transition data rather than the population-genetic framework itself.

## 3 Discussion

Most animals, plants, and unicellular eukaryotes live in association with microorganisms that influence their survival and reproduction. Our results show that this pervasive feature of biology can alter host evolution at a more fundamental level than simply changing host fitness. When individuals move among microbial states with different demographic properties, the strength of genetic drift and selection experienced by host mutations can differ substantially from that predicted from census population size and intrinsic mutational effects alone. As a consequence, microbiome-associated population structure can strongly alter fixation probabilities: deleterious mutations can become orders of magnitude more (or less) likely to fix, and advantageous mutations can also be either suppressed or enhanced. Thus, two host populations with the same census size and the same intrinsic effect of a mutation can evolve differently because their microbial ecology differs.

The main theoretical contribution of this study is a systematic way to calculate these effects. For hosts distributed among an arbitrary number of microbial states, connected by demographic and transition processes, the stochastic ecological dynamics can be mapped onto an effective population size, *N*_eff_, and an effective selection coefficient, *s*_eff_, which together determine mutation fixation. The use of effective population sizes and selection coefficients is well established in population genetics; the advance here is a general procedure for deriving the relevant effective quantities from the underlying host–microbiome dynamics. The calculation incorporates differences in reproductive value among microbial states, the states in which new mutations arise, demographic stochasticity, and transitions generated by microbial acquisition, transmission, and loss. Consequently, the evolutionary effect of a microbiome cannot in general be deduced from its effect on host fitness or prevalence alone. It depends on the broader ecological dynamics through which microbial states are generated and maintained. The applications to experimentally studied host–microbe associations show that these effects occur in biologically realistic parameter regimes. We analyzed systems spanning ciliate, insect, and plant hosts and bacterial, algal, and fungal associates, including one system for which a previously fitted ecological model provides the complete set of demographic and transmission parameters required for a direct quantitative mapping. Across these examples, microbial association altered effective drift, effective selection, or both. The analyses also illustrate that transmission, environmental acquisition, population regulation, and interactions with other species can change the evolutionary consequences of otherwise similar microbial effects on host demography. These results show that significant microbiome-mediated changes in host population genetics are not confined to extreme or unrealistic/purely theoretical parameter combinations.

The empirical parameterizations nevertheless have important limitations. The published datasets used here were collected for diverse ecological purposes rather than specifically to estimate the parameters required by our theory. Some systems therefore provide direct estimates of most relevant rates, whereas others require assumptions or sensitivity analyses for quantities such as microbial acquisition, loss, vertical transmission, or the separate rates of reproduction and mortality. The numerical predictions for individual systems should consequently be interpreted as empirically anchored estimates rather than definitive measurements of *N*_eff_ and *s*_eff_. At the same time, the analysis identifies which quantities matter for evolutionary prediction and, in some cases, shows that not every microscopic rate must be estimated independently because equilibrium relationships constrain the combinations of parameters that enter the effective quantities. This provides a clear target for future experiments: measurements of host demography should be combined with measurements of how hosts move among microbial states through acquisition, loss, and transmission.

This need becomes especially important for natural microbiomes, which are far more complex than most of the systems that can currently be parameterized. Hosts can carry many microbial taxa, strains, and functional groups, producing multiple compositional and physiological states whose demographic effects may differ. The predominantly two-state (and maximally four-state) empirical models analyzed here should therefore not be interpreted as a claim that natural microbiomes are adequately described by the presence or absence of a single microorganism (or a very small number of microorganisms). Rather, they are systems in which the available data allow the theory to be tested quantitatively. The mathematical framework itself is formulated for an arbitrary number of microbial states.

Presently, the principal obstacle to applying the framework to much richer microbiomes is therefore empirical rather than mathematical. A literal state for every possible combination of microbial taxa would rarely be practical or biologically useful. Instead, microbiomes could be coarse-grained into states that differ in features relevant to host evolution, for example, community types, microbial loads, functional configurations, or combinations of taxa associated with distinct host demographic effects. What is required is information about the demographic properties of these states and the rates at which hosts move among them. This suggests an experimental direction that follows directly from the theory: longitudinal measurements of microbiome state should increasingly be linked to host reproduction, survival, and transitions among microbial configurations. Such data would allow the evolutionary consequences of realistic, high-dimensional microbiomes to be calculated.

Several assumptions define the present scope of the theory. The analytical calculation describes rare mutants near a stable resident ecological equilibrium and derives *s*_eff_ under weak selection. The approximation of ultimate fixation further assumes that the effective drift– selection relationship inferred when the mutant is rare remains informative as its frequency increases. Direct simulations support this approximation for the models examined here, but strong frequency dependence, major ecological transients, or changing microbial environments during a selective sweep could require extensions of the approach. We also focused largely on mutations with the same proportional effect across microbial states in order to isolate the consequences of microbiome-associated population structure itself. State-specific mutational effects and genotype–microbiome interactions can be incorporated naturally, as can more elaborate ecological state structures. Spatial dynamics, nonequilibrium microbiomes, and simultaneous host–microbe evolution represent further directions in which the evolutionary environment experienced by a host lineage may itself change through time.

Our results also place host population genetics in the context of the holobiont and hologenome concepts, which emphasize that host phenotype and fitness emerge from interactions between the host and its associated microorganisms [26, 29]. A central debate has been whether host–microbiome assemblages can themselves constitute evolutionary units, particularly when microbial partners are acquired horizontally or transmitted with limited fidelity [27, 28]. The framework developed here does not require resolving this question. We follow the fate of mutations in the host genome while allowing microbial states to be acquired, lost, imperfectly transmitted, and reorganized independently of host genetic inheritance. Nevertheless, these microbial dynamics can substantially alter the effective action of drift and selection on host genes. Thus, a microbiome need not behave as a vertically inherited extension of the host genome, or the host–microbiome assemblage as a single hereditary unit, to have fundamental consequences for host evolution. This provides a direct population-genetic link between the holobiont perspective and the evolution of host genomes, and suggests that understanding evolution in organisms with complex microbiomes will require quantitative knowledge not only of microbial composition, but of the demographic states and transitions generated by those communities.

## 4 Methods

To study host evolution in a population with a dynamically acquired microbiome, we mapped the structured process to an unstructured reference process with effective parameters *N*_eff_ = *N* ℛ_*N*_ and *s*_eff_ = *s* ℛ_*s*_ + *o*(*s*). To first order in weak selection, the fixation probability is then approximated by *P*_fix_(*s*_eff_, *N*_eff_), see equation (1), and neutral limit *P*_fix_ = 1*/N*_eff_. In this framework, the many ecological parameters affect fixation through only two population-genetic modifiers, ℛ_*N*_ and ℛ_*s*_, once the intrinsic *N* and *s* are specified.

To calculate the effective selection coefficient and population size, we project the stochastic mutant lineage onto reproductive value, which measures the expected long-term genetic contribution of a mutant in each microbial state. At neutrality, the reproductive-value-weighted mutant abundance has no systematic increase/decrease. Comparing its mean value when a de novo mutant first appears with its value at fixation gives the neutral fixation probability and hence *N*_eff_. To obtain *s*_eff_, we calculate how selection changes the growth of the reproductive-value-weighted lineage and compare this selective drift with the demographic variance generated by the neutral process. Matching these two quantities to those of an unstructured reference population determines *s*_eff_. The full calculation is presented in the SI.

