## Supplementary Information for "Microbiomes change the rules of host evolution"

**4 More complex microbiome states 43**

### 1 Determining effective selection in a compartmental stochastic population

A stochastic population divided among compartments is generally not equivalent to an unstructured population. Consequently, an effective population size and an effective selection coefficient are not intrinsic quantities, but they can be defined by specifying an unstructured reference process and stating which properties of the structured process are to be matched. Here we present an approach where the reference process is a one-type birth–death process.

In particular, at neutrality, we will construct a scalar martingale from the compartmental mutant counts, and normalize it such that one newly introduced mutant represents one effective mutant. Then we will use the scalar's value at fixation to define the effective population size, $N_{\text{eff}}$.

Table 1: Key population-genetic quantities in the two-state host–microbiome model. State $y$ denotes microbiome-associated hosts, state $x$ unassociated hosts, and $N = N_x + N_y$ is the census population size.

| Quantity | **Definition and interpretation** |
|---|---|
| $P_y^{\text{census}}$ | Fraction of the standing host population in microbial state $y$: $N_y/N$. |
| $P_y^{\text{de novo}}$ | Fraction of newborn hosts (and, assuming equal mutation probability per birth, newly arising mutations) entering state $y$. |
| $\rho_y$ | Reproductive value of a host in state $y$ relative to state $x$ (normalized to 1); values above/below 1 indicate greater/lower expected long-term genetic contribution. |
| $N_{\text{eff}}$ | Effective population size governing genetic drift; larger values correspond to weaker drift. |
| $s_{\text{eff}}$ | Effective selection coefficient governing the strength of selection on host mutations. |
| $\mathcal{R}_N$ | Effective-size modifier: $\mathcal{R}_N = N_{\text{eff}}/N$. Values $\mathcal{R}_N > 1$ indicate weaker drift and $\mathcal{R}_N < 1$ stronger drift relative to the census-size reference. |
| $\mathcal{R}_s$ | Selection modifier: $s_{\text{eff}} = \mathcal{R}_s s + o(s)$. Values $\mathcal{R}_s > 1$ amplify and $\mathcal{R}_s < 1$ suppress the magnitude of selection. |

To determine the effective selection coefficient, $s_{\text{eff}}$, we will use a reference one-type birth death process, and note that the neutral variance determines the effective birth–death timescale, and the leading eigenvalue under weak selection constrains the selective growth rate. Matching these two quantities gives the effective selection coefficient. Below we describe this procedure in some detail.

Suppose mutants can occupy $k$ compartments. The stochastic state of the mutant lineage is therefore a vector, not a single population count. The goal is to replace this structured process by an unstructured process with

- one scalar effective mutant count;
- an effective population size $N_{\text{eff}}$;
- an effective selection coefficient $s_{\text{eff}}$.

There is no unique way to perform this replacement. In the present derivation we use a moment-matching procedure. Neutral fluctuations will determine the effective clock, while selection induces the bias measured relative to that clock. This construction combines reproductive-value weighting of a structured population [1, 2] with a one-dimensional diffusion or fast-mode reduction [3, 4]. The first-order effect of selection is obtained from the standard sensitivity of the dominant eigenvalue [5]. We determine the parameters of the reference birth–death process by matching the long-time growth rate and infinitesimal demographic variance of the reproductive-value projection. Some key notations are listed in table 1.

##### 1.1 Structured continuous-time Markov process

**State, events, and rates.** Let

$$\boldsymbol{M}(t) = \begin{pmatrix} M_1(t) \\ \vdots \\ M_k(t) \end{pmatrix} \in \mathbb{N}_0^k$$

be the random column vector of mutant counts in each compartment at time $t$. Its $i$-th component, $M_i(t)$, is the number of mutants in compartment $i$. We will refer to a particular mutant state as

$\boldsymbol{m}$. We index the possible stochastic events by $r = 1, \dots, R$. Event $r$ changes the mutant state by the fixed column vector

$$\boldsymbol{\nu}_r \in \mathbb{Z}^k.$$

Thus, if event $r$ occurs while the state is $\boldsymbol{m}$, the new state is

$$\boldsymbol{m} + \boldsymbol{\nu}_r.$$

When mutants are rare, the resident population can be treated as fixed at its equilibrium, and the mutant event rates can be linearized in terms of the mutant counts. We write the rate of event $r$ as

$$\alpha_r(\boldsymbol{m}; s) = \boldsymbol{c}_r(s)^T \boldsymbol{m}, \tag{1}$$

where $\boldsymbol{c}_r(s)$ is a column vector of nonnegative per-capita event rates, and $s$ is the microscopic selection parameter.

**Generator and mean matrix.** For any scalar function $f(\boldsymbol{m})$, the generator of the jump process is

$$\mathcal{L}_s f(\boldsymbol{m}) = \sum_{r=1}^{R} \alpha_r(\boldsymbol{m}; s) \left[ f(\boldsymbol{m} + \boldsymbol{\nu}_r) - f(\boldsymbol{m}) \right]. \tag{2}$$

The generator gives the conditional instantaneous mean change of $f$:

$$\mathbb{E}[f(\boldsymbol{M}(t + dt)) - f(\boldsymbol{M}(t)) \mid \boldsymbol{M}(t) = \boldsymbol{m}] = \mathcal{L}_s f(\boldsymbol{m})\, dt + o(dt),$$

and the Dynkin's formula gives

$$\frac{d}{dt} \mathbb{E}[f(\boldsymbol{M}(t))] = \mathbb{E}[\mathcal{L}_s f(\boldsymbol{M}(t))]. \tag{3}$$

Define the $k \times k$ matrix

$$A(s) = \sum_{r=1}^{R} \boldsymbol{\nu}_r \boldsymbol{c}_r(s)^T. \tag{4}$$

To show that this is the mean matrix, apply the generator to the vector-valued identity function $f(\boldsymbol{m}) = \boldsymbol{m}$:

$$\mathcal{L}_s \boldsymbol{m} = \sum_{r=1}^{R} \left( \boldsymbol{c}_r(s)^T \boldsymbol{m} \right) \boldsymbol{\nu}_r = A(s) \boldsymbol{m}.$$

Taking expectations and denoting the mean mutant vector,

$$\overline{\boldsymbol{m}}(t) = \mathbb{E}[\boldsymbol{M}(t)],$$

gives the mean equation in the rare-mutant linear regime:

$$\frac{d}{dt} \overline{\boldsymbol{m}}(t) = A(s) \overline{\boldsymbol{m}}(t). \tag{5}$$

#### 1.2 Neutral assumptions and the reproductive-value martingale

**Neutral eigenvectors.** Let us denote by $A_0$ the neutral matrix,

$$A_0 = A(0),$$

and make the following assumptions:

1. $A_0$ has a simple eigenvalue equal to zero;
2. every other eigenvalue of $A_0$ has negative real part;
3. there is a positive right null vector $\boldsymbol{N}$ satisfying
$$A_0 \boldsymbol{N} = 0; \tag{6}$$
4. there is a positive left null vector $\boldsymbol{w}$ satisfying
$$\boldsymbol{w}^T A_0 = 0. \tag{7}$$

The column vector

$$\boldsymbol{N} = \begin{pmatrix} N_1 \\ \vdots \\ N_k \end{pmatrix}$$

contains the neutral equilibrium compartment sizes. The column vector

$$\boldsymbol{w} = \begin{pmatrix} w_1 \\ \vdots \\ w_k \end{pmatrix}$$

contains the reproductive-value weights. We also define the positive scalar,

$$W = \boldsymbol{w}^T \boldsymbol{N}. \tag{8}$$

**Construction of the martingale.** Using the reproductive-value weights, we construct the following scalar random variable that combines mutant populations of different compartments:

$$Z(t) = \boldsymbol{w}^T \boldsymbol{M}(t). \tag{9}$$

For a fixed state $\boldsymbol{m}$, the change in $Z$ caused by event $r$ is given by

$$\delta_r = \boldsymbol{w}^T \boldsymbol{\nu}_r. \tag{10}$$

The total expected change of $Z(t)$ given $\boldsymbol{m}$ is given (using equations (2) and (1)),

$$\begin{aligned} \mathcal{L}_0 Z(\boldsymbol{m}) &= \sum_{r=1}^{R} \left( \boldsymbol{c}_r(0)^T \boldsymbol{m} \right) \delta_r \\ &= \boldsymbol{w}^T \left[ \sum_{r=1}^{R} \boldsymbol{\nu}_r \boldsymbol{c}_r(0)^T \right] \boldsymbol{m} \\ &= \boldsymbol{w}^T A_0 \boldsymbol{m} = 0. \end{aligned} \tag{11}$$

Therefore $Z(t)$ has zero conditional drift at neutrality and is therefore a martingale. In particular,

$$\mathbb{E}[Z(t)] = \mathbb{E}[Z(0)], \qquad t \geq 0. \tag{12}$$

##### 1.3 Normalizing the martingale and defining $N_{\text{eff}}$

**Initial mutant placement.** Suppose one mutant is introduced. Let $P_i$ be the probability that it is introduced in compartment $i$, with $\sum_{i=1}^{k} P_i = 1$. Define the initial-placement probability vector

$$\boldsymbol{P} = \begin{pmatrix} P_1 \\ \vdots \\ P_k \end{pmatrix}.$$

Because there is initially one mutant,

$$\overline{\boldsymbol{m}}(0) = \mathbb{E}[\boldsymbol{M}(0)] = \boldsymbol{P}. \tag{13}$$

Denote by $z_0$ the mean initial value of the martingale:

$$z_0 = \mathbb{E}[Z(0)] = \boldsymbol{w}^T\boldsymbol{P} = \sum_{i=1}^{k} P_i w_i. \tag{14}$$

**Effective mutant count.** Define the scalar effective mutant count

$$\mathcal{Z}(t) = \frac{Z(t)}{z_0}. \tag{15}$$

This normalization is chosen so that $\mathbb{E}[\mathcal{Z}(0)] = 1$. We now assume that the neutral martingale extends to the full finite-population process and that this process is eventually absorbed either at mutant extinction ($\boldsymbol{M} = \boldsymbol{0}$) or at mutant fixation. At fixation, the mutant population is taken to have the resident equilibrium composition, $\boldsymbol{M} = \boldsymbol{N}$. The value of $\mathcal{Z}$ at extinction is $\mathcal{Z}_{\text{ext}} = 0$ and at fixation it is $\mathcal{Z}_{\text{fix}} = \frac{\boldsymbol{w}^T\boldsymbol{N}}{z_0} = \frac{W}{z_0}$. Therefore, at absorption (time $\tau$), we have

$$\mathbb{E}[\mathcal{Z}(\tau)] = P_{\text{fix}}^{(0)}\mathcal{Z}_{\text{fix}} + (1 - P_{\text{fix}}^{(0)})\mathcal{Z}_{\text{ext}} = P_{\text{fix}}^{(0)}\frac{W}{z_0}.$$

Because $\mathcal{Z}(t)$ is a martingale,

$$\mathbb{E}[\mathcal{Z}(t)] = \mathbb{E}[\mathcal{Z}(0)] = 1$$

for every $t$. In the full finite-population process, $\mathcal{Z}(t)$ is bounded; hence, if the absorption time $\tau$ is almost surely finite, the optional-stopping theorem gives

$$\mathbb{E}[\mathcal{Z}(\tau)] = \mathbb{E}[\mathcal{Z}(0)] = 1,$$

and therefore $P_{\text{fix}}^{(0)} = \left(\frac{W}{z_0}\right)^{-1}$. It follows that the quantity

$$N_{\text{eff}} = \frac{W}{z_0} = \frac{\boldsymbol{w}^T\boldsymbol{N}}{\boldsymbol{w}^T\boldsymbol{P}} \tag{16}$$

plays the role of the effective population size and we have

$$P_{\text{fix}}^{(0)} = \frac{1}{N_{\text{eff}}}.$$

In other words, equation (16) gives the size of an unstructured neutral population whose fixation probability from one mutant agrees with that of the structured process. Note that $N_{\text{eff}}$ in equation (16) depends not only on the equilibrium composition and reproductive values, but also on the mechanism determining the compartment in which a new mutant first appears.

It is convenient to define an evolutionary modifier of the population size, which we denote by $\mathcal{R}_N$:

$$\mathcal{R}_N = \frac{N_{\text{eff}}}{N} = \frac{1}{N}\frac{\boldsymbol{w}^T\boldsymbol{N}}{\boldsymbol{w}^T\boldsymbol{P}} \tag{17}$$

**Comparison with census population size.** Let

$$N = \sum_{i=1}^{k} N_i$$

be the total equilibrium population size, and define the census distribution among compartments by

$$P_i^{\text{census}} = \frac{N_i}{N}. \tag{18}$$

The initial-placement probabilities $P_i$ define the corresponding distribution of newly introduced mutants. When mutants arise de novo, that is, when they are produced by resident individuals through a birth event, we write

$$P_i^{\text{de novo}} = P_i. \tag{19}$$

The mean reproductive values of a randomly chosen individual (a census individual) and a newly born individual (or a de novo mutant) are, respectively,

$$\langle w \rangle_{\text{census}} = \boldsymbol{w}^{\mathsf{T}} \boldsymbol{P}^{\text{census}} = \frac{W}{N}, \qquad \langle w \rangle_{\text{de novo}} = \boldsymbol{w}^{\mathsf{T}} \boldsymbol{P}^{\text{de novo}} = z_0.$$

It follows that the population size modifier, equation (17), can be expressed as

$$\mathcal{R}_N = \frac{N_{\text{eff}}}{N} = \frac{\langle w \rangle_{\text{census}}}{\langle w \rangle_{\text{de novo}}}. \tag{20}$$

Thus, $\mathcal{R}_N > 1$ when a randomly sampled census individual has a higher mean reproductive value than a newly arising mutant, and $\mathcal{R}_N < 1$ when the reverse is true.

#### 1.4 Neutral variance of the reproductive-value coordinate

We have shown that at neutrality, the quantity $Z(t)$ is a martingale, and hence its expectation is constant. Individual events nevertheless change $Z(t)$, causing its variance to increase. We now calculate the rate at which this variance is produced.

Recall that event $r$ changes the mutant state by $\boldsymbol{\nu}_r$ and therefore changes $Z$ by

$$\delta_r = \boldsymbol{w}^{\mathsf{T}} \boldsymbol{\nu}_r.$$

For a fixed mutant state $\boldsymbol{m}$, write $Z(\boldsymbol{m}) = \boldsymbol{w}^{\mathsf{T}} \boldsymbol{m}$. Applying the neutral generator to $Z^2$ gives

$$\begin{aligned} \mathcal{L}_0 Z^2(\boldsymbol{m}) &= \sum_{r=1}^{R} \alpha_r(\boldsymbol{m}; 0) \left[ \left( Z(\boldsymbol{m}) + \delta_r \right)^2 - Z(\boldsymbol{m})^2 \right] \\ &= 2 Z(\boldsymbol{m})\, \mathcal{L}_0 Z(\boldsymbol{m}) + \sum_{r=1}^{R} \alpha_r(\boldsymbol{m}; 0) \delta_r^2. \end{aligned} \tag{21}$$

The first term vanishes because $\mathcal{L}_0 Z(\boldsymbol{m}) = 0$. Using $\alpha_r(\boldsymbol{m}; 0) = \boldsymbol{c}_r(0)^{\mathsf{T}} \boldsymbol{m}$ and taking the mean, by equation (3) we have

$$\frac{d}{dt} \mathbb{E}[Z(t)^2] = \sum_{r=1}^{R} \boldsymbol{c}_r(0)^{\mathsf{T}} \overline{\boldsymbol{m}}(t)\, \delta_r^2, \tag{22}$$

where

$$\overline{\boldsymbol{m}}(t) = \mathbb{E}[\boldsymbol{M}(t)].$$

Because $\mathbb{E}[Z(t)]$ is constant at neutrality,

$$\frac{d}{dt}\operatorname{Var}[Z(t)] = \frac{d}{dt}\left\{\mathbb{E}[Z(t)^2] - \mathbb{E}[Z(t)]^2\right\} = \frac{d}{dt}\mathbb{E}[Z(t)^2]. \tag{23}$$

Thus,

$$\frac{d}{dt}\operatorname{Var}[Z(t)] = \sum_{r=1}^{R} \boldsymbol{c}_r(0)^{\mathsf{T}}\overline{\boldsymbol{m}}(t)\,\delta_r^2. \tag{24}$$

**Long-time neutral mean composition.** At neutrality, initial-value problem (5,13) becomes

$$\frac{d}{dt}\overline{\boldsymbol{m}}(t) = A_0\overline{\boldsymbol{m}}(t), \qquad \overline{\boldsymbol{m}}(0) = \boldsymbol{P}.$$

Because zero is a simple eigenvalue of $A_0$ and all its other eigenvalues have negative real parts,

$$\overline{\boldsymbol{m}}(t) \longrightarrow c\boldsymbol{N} \qquad \text{as } t \to \infty$$

for some scalar $c$. Moreover,

$$\frac{d}{dt}\left[\boldsymbol{w}^{\mathsf{T}}\overline{\boldsymbol{m}}(t)\right] = \boldsymbol{w}^{\mathsf{T}}A_0\overline{\boldsymbol{m}}(t) = 0.$$

Hence

$$\boldsymbol{w}^{\mathsf{T}}\overline{\boldsymbol{m}}(t) = \boldsymbol{w}^{\mathsf{T}}\boldsymbol{P} = z_0$$

for every $t$. Since $\overline{\boldsymbol{m}}(t) \to c\boldsymbol{N}$, it follows that

$$z_0 = \boldsymbol{w}^{\mathsf{T}}(c\boldsymbol{N}) = cW.$$

Therefore,

$$\overline{\boldsymbol{m}}(t) \longrightarrow \frac{z_0}{W}\boldsymbol{N} \qquad \text{as } t \to \infty. \tag{25}$$

This result concerns the mean mutant vector; it does not imply that individual realizations converge to $(z_0/W)\boldsymbol{N}$.

Define the nonnegative scalar

$$G = \sum_{r=1}^{R} \left[\boldsymbol{c}_r(0)^{\mathsf{T}}\boldsymbol{N}\right]\left(\boldsymbol{w}^{\mathsf{T}}\boldsymbol{\nu}_r\right)^2. \tag{26}$$

Using equation (25) in equation (24), we obtain

$$\begin{aligned}\lim_{t\to\infty}\frac{d}{dt}\operatorname{Var}[Z(t)] &= \sum_{r=1}^{R} \boldsymbol{c}_r(0)^{\mathsf{T}}\left(\frac{z_0}{W}\boldsymbol{N}\right)\left(\boldsymbol{w}^{\mathsf{T}}\boldsymbol{\nu}_r\right)^2 \\ &= \frac{z_0}{W}G. \end{aligned}\tag{27}$$

Because the normalized effective mutant count is $\mathcal{Z}(t) = Z(t)/z_0$, its asymptotic neutral variance-production rate is

$$\lim_{t\to\infty}\frac{d}{dt}\operatorname{Var}[\mathcal{Z}(t)] = \frac{G}{z_0 W}. \tag{28}$$

Because every term in the definition of $G$ is nonnegative, $G > 0$ provided that at least one event occurring at a positive rate at equilibrium changes the reproductive-value coordinate, that is, provided that

$$\alpha_r(\boldsymbol{N};0) > 0 \quad \text{and} \quad \boldsymbol{w}^{\mathsf{T}}\boldsymbol{\nu}_r \neq 0$$

for at least one $r$.

##### 1.5 Growth of the reproductive-value coordinate under weak selection

We now consider the same structured mutant process in the presence of weak selection. Assume that $A(s)$ is differentiable at $s = 0$ and write

$$A(s) = A_0 + sA_1 + o(s), \qquad A_1 = \left.\frac{\partial A(s)}{\partial s}\right|_{s=0}. \tag{29}$$

By the spectral gap at neutrality and the continuity of the eigenvalues, for sufficiently small $|s|$, $A(s)$ has a unique eigenvalue $\lambda(s)$ near zero, while all its other eigenvalues retain strictly negative real parts bounded away from zero. Thus, the eigenvalue emerging from the neutral eigenvalue governs the long-time behaviour under weak selection. Let $\lambda(s)$ denote the leading eigenvalue of $A(s)$ and let $\boldsymbol{u}(s)$ be its corresponding right eigenvector:

$$A(s)\boldsymbol{u}(s) = \lambda(s)\boldsymbol{u}(s). \tag{30}$$

At neutrality,

$$\lambda(0) = 0, \qquad \boldsymbol{u}(0) = \boldsymbol{N}.$$

Differentiating equation (30) with respect to $s$ and evaluating at $s = 0$ gives

$$A_1\boldsymbol{N} + A_0\boldsymbol{u}'(0) = \lambda'(0)\boldsymbol{N}. \tag{31}$$

Multiplying from the left by $\boldsymbol{w}^\top$ eliminates the second term because $\boldsymbol{w}^\top A_0 = 0$. Hence

$$\boldsymbol{w}^\top A_1\boldsymbol{N} = \lambda'(0)\boldsymbol{w}^\top\boldsymbol{N}.$$

Define

$$L_{\mathrm{sel}} := \boldsymbol{w}^\top A_1\boldsymbol{N}. \tag{32}$$

Using $W = \boldsymbol{w}^\top\boldsymbol{N}$, we obtain

$$\lambda'(0) = \frac{L_{\mathrm{sel}}}{W}, \tag{33}$$

and therefore

$$\lambda(s) = \frac{L_{\mathrm{sel}}}{W}s + o(s). \tag{34}$$

The mean mutant vector satisfies the initial value problem (5,13), whose solution is

$$\overline{\boldsymbol{m}}(t) = e^{A(s)t}\boldsymbol{P}, \tag{35}$$

where $e^{A(s)t}$ denotes the matrix exponential. After the transient compartmental modes have decayed, its leading component grows or declines as $e^{\lambda(s)t}$. Consequently, provided that the initial mutant distribution has a nonzero component along the leading mode,

$$\mathbb{E}[\mathcal{Z}(t)] \sim C(s)e^{\lambda(s)t}$$

for some prefactor $C(s)$. Dividing $Z(t)$ by the constant $z_0$ does not change this exponential rate. Thus, to first order in $s$, the asymptotic selective growth rate of the normalized reproductive-value coordinate is

$$\frac{L_{\mathrm{sel}}}{W}s. \tag{36}$$

This quantity has units of inverse time. It is a population-growth rate, which we will use to find the effective selection coefficient.

##### 1.6 Unstructured reference process and moment matching

We have now obtained two properties of the normalized reproductive-value coordinate $\mathcal{Z}(t)$: its asymptotic neutral variance-production rate and its asymptotic growth rate under weak selection. We introduce an unstructured one-type reference process whose parameters are chosen to reproduce these two quantities.

Let $Y(t)$ be a linear birth–death process with $Y(0) = 1$. When $Y(t) = y$, let

$$y \longrightarrow y + 1 \qquad \text{at rate} \qquad b_{\text{eff}}(1 + s_{\text{eff}})y,$$

and

$$y \longrightarrow y - 1 \qquad \text{at rate} \qquad b_{\text{eff}}y.$$

The parameter $b_{\text{eff}} > 0$ sets the demographic timescale, whereas the dimensionless parameter $s_{\text{eff}}$ describes the selective bias relative to that timescale.

At neutrality, $s_{\text{eff}} = 0$, births and deaths occur at equal rates. Since $\mathbb{E}[Y(t)] = 1$ when $Y(0) = 1$, its variance grows at the constant rate

$$\frac{d}{dt}\,\text{Var}[Y(t)] = 2b_{\text{eff}}. \tag{37}$$

Under selection, its mean satisfies

$$\frac{d}{dt}\mathbb{E}[Y(t)] = b_{\text{eff}}s_{\text{eff}}\,\mathbb{E}[Y(t)], \tag{38}$$

so its selective exponential growth rate is $b_{\text{eff}}s_{\text{eff}}$.

We first match the neutral variance-production rates of $Y(t)$ and $\mathcal{Z}(t)$. Equations (28) and (37) give

$$2b_{\text{eff}} = \frac{G}{z_0 W}, \tag{39}$$

and hence

$$b_{\text{eff}} = \frac{G}{2z_0 W}. \tag{40}$$

We then match their selective exponential growth rates. Equations (36) and (38) give

$$b_{\text{eff}}s_{\text{eff}} = \frac{L_{\text{sel}}}{W}s + o(s). \tag{41}$$

Substituting equation (40) and solving for $s_{\text{eff}}$ yields

$$s_{\text{eff}} = \frac{2z_0 L_{\text{sel}}}{G}s + o(s). \tag{42}$$

Using $z_0 = \boldsymbol{w}^\mathsf{T}\boldsymbol{P}$, $L_{\text{sel}} = \boldsymbol{w}^\mathsf{T}A_1\boldsymbol{N}$ and equation (26), this can be written explicitly as

$$s_{\text{eff}} = \frac{2\left(\boldsymbol{w}^\mathsf{T}\boldsymbol{P}\right)\left(\boldsymbol{w}^\mathsf{T}A_1\boldsymbol{N}\right)}{\displaystyle\sum_{r=1}^{R}\left[\boldsymbol{c}_r(0)^\mathsf{T}\boldsymbol{N}\right]\left(\boldsymbol{w}^\mathsf{T}\boldsymbol{\nu}_r\right)^2}\,s + o(s). \tag{43}$$

The evolutionary modifier of selection, $Rs$, is then given by

$$\mathcal{R}_s = \lim_{|s|\to 0}\frac{s_{\text{eff}}}{s} = \frac{2\left(\boldsymbol{w}^\mathsf{T}\boldsymbol{P}\right)\left(\boldsymbol{w}^\mathsf{T}A_1\boldsymbol{N}\right)}{\displaystyle\sum_{r=1}^{R}\left[\boldsymbol{c}_r(0)^\mathsf{T}\boldsymbol{N}\right]\left(\boldsymbol{w}^\mathsf{T}\boldsymbol{\nu}_r\right)^2}. \tag{44}$$

We note that the reference process $Y(t)$ is not assumed to have the same complete probability distribution as $\mathcal{Z}(t)$. It provides an unstructured standard against which the structured process is calibrated: $b_{\text{eff}}$ matches its neutral demographic fluctuations, and $s_{\text{eff}}$ matches its weak-selection growth rate relative to those fluctuations.

##### 1.7 Diffusion interpretation and finite-population closure

After the stable compartmental modes have relaxed, a state with normalized effective mutant count $\mathcal{Z} = z$ has approximate compartmental composition

$$\boldsymbol{m} \simeq \frac{z_0 z}{W} \boldsymbol{N}.$$

At neutrality, the corresponding infinitesimal variance rate of $\mathcal{Z}$ is therefore

$$\frac{1}{z_0^2} \sum_{r=1}^{R} \left[ \boldsymbol{c}_r(0)^{\mathsf{T}} \left( \frac{z_0 z}{W} \boldsymbol{N} \right) \right] \left( \boldsymbol{w}^{\mathsf{T}} \boldsymbol{\nu}_r \right)^2 = \frac{G}{z_0 W} z.$$

Thus, near the rare-mutant boundary and after the faster compartmental modes have relaxed, the effective coordinate has the diffusion approximation

$$d\mathcal{Z}_t \simeq \frac{L_{\text{sel}}}{W} s \mathcal{Z}_t \, dt + \sqrt{\frac{G}{z_0 W} \mathcal{Z}_t} \, dB_t, \tag{45}$$

where the demographic-noise coefficient is evaluated at neutrality. The corresponding weak-selection approximation for the reference process is

$$dY_t \simeq b_{\text{eff}} s_{\text{eff}} Y_t \, dt + \sqrt{2 b_{\text{eff}} Y_t} \, d\widetilde{B}_t. \tag{46}$$

Matching the neutral variance coefficients and the weak-selection drift coefficients in equations (45) and (46) reproduces equations (40) and (42).

The derivation above describes the process near the rare-mutant boundary. To obtain a finite-population fixation probability, we make the additional approximation that the same $s_{\text{eff}}$ describes a one-dimensional birth–death chain between the absorbing boundaries 0 and $N_{\text{eff}}$. Then the fixation probability from one effective mutant is

$$P_{\text{fix}} = \frac{1 - (1 + s_{\text{eff}})^{-1}}{1 - (1 + s_{\text{eff}})^{-N_{\text{eff}}}}, \tag{47}$$

with the neutral value $P_{\text{fix}} = 1/N_{\text{eff}}$ obtained by continuity as $s_{\text{eff}} \to 0$. For integer $N_{\text{eff}}$, equation (47) is exact for the stated one-dimensional reference chain; for noninteger $N_{\text{eff}}$, its right-hand side provides the corresponding continuous interpolation. Its application to the original structured population is an effective approximation, since the derivation of $s_{\text{eff}}$ is local to the rare-mutant regime and does not prove that the complete finite-population process is Moran.

**Summary of formulas.** We have derived explicit expressions for the effective population size, $N_{\text{eff}}$, equation (16), and effective selection coefficient, $s_{\text{eff}}$, equation (43). The results for the evolutionary modifiers are derived in (17) and (44). We have

$$\mathcal{R}_N = \frac{N_{\text{eff}}}{N} = \frac{1}{N} \frac{\boldsymbol{w}^T \boldsymbol{N}}{\boldsymbol{w}^T \boldsymbol{P}}, \tag{48}$$

$$\mathcal{R}_s = \lim_{|s| \to 0} \frac{s_{\text{eff}}}{s} = \frac{2 \left( \boldsymbol{w}^{\mathsf{T}} \boldsymbol{P} \right) \left( \boldsymbol{w}^{\mathsf{T}} A_1 \boldsymbol{N} \right)}{\sum_{r=1}^{R} \left[ \boldsymbol{c}_r(0)^{\mathsf{T}} \boldsymbol{N} \right] \left( \boldsymbol{w}^{\mathsf{T}} \boldsymbol{\nu}_r \right)^2}. \tag{49}$$

### 2 Example: a two-compartment microbiome model with vertical and horizontal infection-type transmission

In this section we work out a specific example in some detail. We first show all the calculations for this specific system, and later investigate properties of the quantities $\mathcal{R}_N$ and $\mathcal{R}_s$ that we obtained.

#### 2.1 Biological events and their rates

The population is divided into two compartments. An individual in compartment $x$ does not carry the microbiome, whereas an individual in compartment $y$ carries it. A division in compartment $y$ produces a newborn that retains the microbiome with probability $v$ and loses it with probability $1-v$. The parent remains in compartment $y$ in both cases. The complete set of events is shown in table 2.

The resident mean-field equations generated by these events are

$$\dot{x} = r_x x\left(1-\frac{x+y}{K}\right) + (1-v)r_y y\left(1-\frac{x+y}{K}\right) - d_x x - \beta xy, \tag{50}$$

$$\dot{y} = \beta xy + v r_y y\left(1-\frac{x+y}{K}\right) - d_y y. \tag{51}$$

Let $(N_x, N_y)$ be a positive resident equilibrium and define the scalars

$$N = N_x + N_y, \qquad b_x = r_x\left(1-\frac{N}{K}\right), \qquad b_y = r_y\left(1-\frac{N}{K}\right).$$

The equilibrium conditions are

$$\big(b_x - d_x - \beta N_y\big)N_x + (1-v)b_y N_y = 0, \tag{52}$$

$$\beta N_x N_y + \big(v b_y - d_y\big)N_y = 0. \tag{53}$$

#### 2.2 Existence and stability of resident equilibria

The effective quantities derived below describe mutant dynamics around a specified resident equilibrium. They are biologically meaningful only when that equilibrium is admissible and locally stable. We therefore examine these conditions before applying the evolutionary calculation. In particular, a positive coexistence equilibrium must satisfy

$$N_x > 0, \qquad N_y > 0, \qquad N_x + N_y < K,$$

the last condition ensuring that the division rates $b_x$ and $b_y$ are nonnegative.

The resident system can have the extinction equilibrium, a microbiome-free equilibrium, and a positive coexistence equilibrium. A microbiome-associated equilibrium with $x = 0$, $y > 0$, and a positive division rate exists only in the boundary case $v = 1$. When $v < 1$, division of microbiome-associated individuals continually produces newborns in compartment $x$.

**Local stability of a positive coexistence equilibrium.** Let $(N_x, N_y)$ be a positive equilibrium, let

$$N = N_x + N_y, \qquad b_x = r_x\left(1-\frac{N}{K}\right), \qquad b_y = r_y\left(1-\frac{N}{K}\right),$$

| Event | Resident rate at state $(x,y)$ | Rare-mutant rate at resident equilibrium | Mutant jump $\boldsymbol{\nu}$ |
|---|---|---|---|
| Division in $x$, producing an $x$-newborn | $r_x x\left(1-\dfrac{x+y}{K}\right)$ | $(1+s)b_x m_x$ | $(1,0)^\top$ |
| Death in $x$ | $d_x x$ | $d_x m_x$ | $(-1,0)^\top$ |
| Infection, $x \longrightarrow y$ | $\beta xy$ | $\beta N_y m_x$ | $(-1,1)^\top$ |
| Division in $y$, with microbiome retention in the newborn | $vr_y y\left(1-\dfrac{x+y}{K}\right)$ | $v(1+s)b_y m_y$ | $(0,1)^\top$ |
| Division in $y$, with microbiome loss in the newborn | $(1-v)r_y y\left(1-\dfrac{x+y}{K}\right)$ | $(1-v)(1+s)b_y m_y$ | $(1,0)^\top$ |
| Death in $y$ | $d_y y$ | $d_y m_y$ | $(0,-1)^\top$ |

Table 2: Events and rates of the resident and rare-mutant processes.

and define

$$C := \frac{r_x N_x + (1-v) r_y N_y}{K}.$$

Using the equilibrium conditions, the Jacobian at $(N_x, N_y)$ can be written as

$$J_* = \begin{pmatrix} -\dfrac{(1-v)b_y N_y}{N_x} - C & (1-v)b_y - \beta N_x - C \\ N_y\left(\beta - \dfrac{vr_y}{K}\right) & -\dfrac{vr_y N_y}{K} \end{pmatrix}. \tag{54}$$

Its trace is

$$\operatorname{tr} J_* = -\frac{(1-v)b_y N_y}{N_x} - C - \frac{vr_y N_y}{K} < 0.$$

Consequently, a positive equilibrium is locally asymptotically stable if and only if

$$\det J_* > 0. \tag{55}$$

If $\det J_* < 0$, the equilibrium is a saddle, while $\det J_* = 0$ makes the equilibrium nonhyperbolic and identifies a possible local bifurcation boundary. Thus every equilibrium used to calculate $\mathcal{R}_N$ and $\mathcal{R}_s$ below must first be checked against condition (55).

**The case $\beta = 0$.** Suppose first that $0 < v < 1$. At a positive coexistence equilibrium, the second equilibrium equation gives

$$\ell_0 := 1 - \frac{N_x^{(0)} + N_y^{(0)}}{K} = \frac{d_y}{vr_y}.$$

Hence the total equilibrium population is

$$N^{(0)} = N_x^{(0)} + N_y^{(0)} = K\left(1 - \frac{d_y}{vr_y}\right).$$

The first equilibrium equation gives

$$q_0 := \frac{N_y^{(0)}}{N_x^{(0)}} = \frac{vd_x - (r_x/r_y)d_y}{(1-v)d_y},$$

and therefore

$$N_x^{(0)} = \frac{N^{(0)}}{1+q_0}, \qquad N_y^{(0)} = \frac{q_0 N^{(0)}}{1+q_0}.$$

This equilibrium is positive if and only if

$$d_y < vr_y, \qquad d_x > \frac{r_x d_y}{vr_y}. \tag{56}$$

The first inequality ensures $0 < N^{(0)} < K$, whereas the second ensures $q_0 > 0$.

Whenever the inequalities in (56) are strict, the coexistence equilibrium is locally asymptotically stable. Indeed,

$$\det J_* = \frac{vr_y N_y^{(0)}}{K} \left[d_x - r_x\ell_0 + (1-v)r_y\ell_0\right] > 0,$$

because positivity of $q_0$ implies $d_x - r_x\ell_0 > 0$.

The endpoint $v = 1$ is degenerate. Positive coexistence at $\beta = 0$ is possible only when

$$\frac{d_x}{r_x} = \frac{d_y}{r_y} < 1.$$

In that case every positive pair satisfying

$$N_x + N_y = K\left(1 - \frac{d_x}{r_x}\right)$$

is an equilibrium. These equilibria form a continuum, so no individual equilibrium is locally asymptotically stable; correspondingly, the Jacobian has a zero eigenvalue. When $v = 0$ and $d_y > 0$, no positive coexistence equilibrium exists at $\beta = 0$.

**The microbiome-free equilibrium and increasing $\beta$.** Suppose $0 < v < 1$ and $r_x > d_x$. The microbiome-free equilibrium is

$$E_x = (X, 0), \qquad X = K\left(1 - \frac{d_x}{r_x}\right).$$

Its two eigenvalues are

$$-(r_x - d_x) \qquad \text{and} \qquad \lambda_y(\beta) = \beta X + \frac{vr_y d_x}{r_x} - d_y.$$

Thus $E_x$ is locally asymptotically stable precisely when $\lambda_y(\beta) < 0$. The quantity $\lambda_y(\beta)$ is also the per-capita growth rate of a rare microbiome-associated population introduced at $E_x$.

If

$$\frac{vr_y d_x}{r_x} > d_y,$$

then microbiome-associated individuals can invade already at $\beta = 0$. This is the same parameter region in which the positive equilibrium described above exists at $\beta = 0$. Because that equilibrium is locally stable and nondegenerate, it continues smoothly for sufficiently small positive $\beta$. Its stability at larger $\beta$ must still be checked using (55).

If instead

$$\frac{vr_y d_x}{r_x} < d_y,$$

then $E_x$ is stable at $\beta = 0$ and loses stability at

$$\beta_c = \frac{r_x d_y - vr_y d_x}{K(r_x - d_x)}. \tag{57}$$

At $\beta = \beta_c$, the microbiome-free equilibrium and a coexistence branch meet in a transcritical bifurcation.

To determine the local direction of this branch, define

$$F_{\text{bif}} = vr_y(r_x - d_x)^2 + (d_y - vr_y)\left[r_x(r_x - d_x + d_y) - r_y d_x\right].$$

Provided $F_{\text{bif}} \neq 0$, the coexistence branch satisfies

$$N_y = \frac{K^2(r_x - d_x)^3}{r_x F_{\text{bif}}}(\beta - \beta_c) + O\big((\beta - \beta_c)^2\big). \tag{58}$$

Therefore:

- If $F_{\text{bif}} > 0$, the positive coexistence branch lies on the $\beta > \beta_c$ side of the bifurcation. Locally, stability is exchanged with $E_x$, so the coexistence branch is stable sufficiently close to $\beta_c$.
- If $F_{\text{bif}} < 0$, the positive coexistence branch lies on the $\beta < \beta_c$ side and is initially unstable. This is a backward transcritical bifurcation. It allows the possibility of bistability, but does not by itself establish it.
- If $F_{\text{bif}} = 0$, equation (58) is not valid and higher-order terms are required to determine the local bifurcation structure.

In the backward case, numerical or analytical continuation may show that the unstable coexistence branch undergoes a saddle-node bifurcation at some $\beta_{\text{SN}} < \beta_c$ and connects to a stable coexistence branch. If such a saddle-node exists, then

$$\beta_{\text{SN}} < \beta < \beta_c$$

is a bistable interval containing both the stable microbiome-free equilibrium and a stable coexistence equilibrium, separated by an unstable coexistence equilibrium. Population establishment then requires crossing a finite-abundance threshold and the system exhibits a strong Allee-type effect. These conclusions are conditional on establishing the saddle-node; they do not follow from the sign of $F_{\text{bif}}$ alone.

In all subsequent calculations, $(N_x, N_y)$ denotes a positive equilibrium satisfying

$$N_x + N_y < K, \qquad \det J_* > 0,$$

with all demographic rates nonnegative. If several stable coexistence equilibria are present, $\mathcal{R}_N$ and $\mathcal{R}_s$ must be calculated separately for each equilibrium; their values depend on the resident branch around which the mutant process is linearized.

##### 2.3 Calculating the evolutionary modifiers

We now apply the general construction of Section 1 to this two-compartment process.

**Neutral mean matrix and reproductive values.** The rare-mutant state is

$$\boldsymbol{M}(t) = \begin{pmatrix} M_x(t) \\ M_y(t) \end{pmatrix}, \qquad \overline{\boldsymbol{m}}(t) = \mathbb{E}[\boldsymbol{M}(t)].$$

The event rates and jumps are listed in Table 2. Vector $\boldsymbol{m}$ satisfies equation (5), where

$$A(s) = \begin{pmatrix} (1+s)b_x - d_x - \beta N_y & (1-v)(1+s)b_y \\ \beta N_y & v(1+s)b_y - d_y \end{pmatrix}.$$

At neutrality,

$$A_0 \equiv A(0) = \begin{pmatrix} b_x - d_x - \beta N_y & (1-v)b_y \\ \beta N_y & vb_y - d_y \end{pmatrix}, \qquad A_0 \begin{pmatrix} N_x \\ N_y \end{pmatrix} = 0.$$

The equilibrium equations also give

$$\operatorname{tr} A_0 = -\frac{(1-v)b_y N_y}{N_x} - \beta N_x < 0,$$

so the zero eigenvalue is simple and the other eigenvalue is negative, except at the degenerate boundary $\beta = 0,\ v = 1$.

For $\beta > 0$, it is convenient to normalize the reproductive values by setting $w_x = 1$. Solving $\boldsymbol{w}^{\mathsf{T}} A_0 = 0$ then gives

$$\boldsymbol{w} = \begin{pmatrix} 1 \\ \rho_y \end{pmatrix}, \qquad \rho_y = \frac{(1-v)b_y}{\beta N_x}. \tag{59}$$

Thus

$$Z(t) = M_x(t) + \rho_y M_y(t)$$

is a neutral martingale. An unnormalized left null vector that remains well defined at $\beta = 0$ is

$$\boldsymbol{w} = \begin{pmatrix} \beta N_x N_y \\ (1-v)b_y N_y \end{pmatrix}.$$

Accordingly, formulas involving $\rho_y$ assume $\beta > 0$. Nondegenerate boundary cases may be evaluated using the unnormalized vector or by taking a limit. At $\beta = 0,\ v = 1$, however, the neutral zero eigenvalue is not simple and the assumptions of Section 1 fail; the effective quantities are therefore not defined for an individual equilibrium on this continuum.

**De novo seeding and the effective population size.** Under de novo seeding, a mutation appears in the compartment occupied by the newborn. The total resident division rate is $b_x N_x + b_y N_y$. An $x$-newborn is produced either by division in $x$ or by division in $y$ followed by microbiome loss. A $y$-newborn is produced by division in $y$ followed by microbiome retention. Assuming that the mutation probability per newborn is the same for all division events, the initial compartment of a de novo mutant is distributed in proportion to the production rates of $x$- and $y$-newborns. Therefore, denoting by $P_x$ and $P_y$ the probabilities that the first mutant appears in compartments $x$ and $y$, respectively, we have

$$P_x = \frac{b_x N_x + (1-v)b_y N_y}{b_x N_x + b_y N_y}, \qquad P_y = \frac{vb_y N_y}{b_x N_x + b_y N_y}.$$

| Event | Neutral rate at $\boldsymbol{N}$ | $\delta$ | Variance contribution |
|---|---|---|---|
| Division in $x$ | $b_x N_x$ | $w_x$ | $b_x N_x w_x^2$ |
| Death in $x$ | $d_x N_x$ | $-w_x$ | $d_x N_x w_x^2$ |
| Infection, $x \longrightarrow y$ | $\beta N_x N_y$ | $w_y - w_x$ | $\beta N_x N_y (w_y - w_x)^2$ |
| Division in $y$, microbiome retained | $v b_y N_y$ | $w_y$ | $v b_y N_y w_y^2$ |
| Division in $y$, microbiome lost | $(1-v) b_y N_y$ | $w_x$ | $(1-v) b_y N_y w_x^2$ |
| Death in $y$ | $d_y N_y$ | $-w_y$ | $d_y N_y w_y^2$ |

Table 3: Calculating the instantaneous variance production. For each process, its neutral rate, the change in $Z$, and its contribution to the instantaneous variance production are listed.

We have

$$W = N_x + \rho_y N_y, \qquad z_0 = P_x + \rho_y P_y,$$

and the general result in equation (16) gives

$$N_{\text{eff}} = \frac{N_x + \rho_y N_y}{P_x + \rho_y P_y}.$$

The population-size modifier can be interpreted as a ratio of mean reproductive values. For the two-compartment model,

$$P_y^{\text{census}} = \frac{N_y}{N_x + N_y}, \qquad P_y^{\text{de novo}} = P_y.$$

With the normalization $w_x = 1$ and $w_y = \rho_y$, the mean reproductive value of a uniformly sampled census individual is

$$\langle w \rangle_{\text{census}} = P_x^{\text{census}} + \rho_y P_y^{\text{census}} = 1 + (\rho_y - 1) P_y^{\text{census}} = \frac{W}{N}. \tag{60}$$

The mean reproductive value of a newly arising mutant is

$$\langle w \rangle_{\text{de novo}} = P_x^{\text{de novo}} + \rho_y P_y^{\text{de novo}} = 1 + (\rho_y - 1) P_y^{\text{de novo}} = z_0. \tag{61}$$

Consequently,

$$\mathcal{R}_N = \frac{N_{\text{eff}}}{N} = \frac{\langle w \rangle_{\text{census}}}{\langle w \rangle_{\text{de novo}}} = \frac{1 + (\rho_y - 1) P_y^{\text{census}}}{1 + (\rho_y - 1) P_y^{\text{de novo}}}. \tag{62}$$

**Determining the effective selection coefficient.** Because selection multiplies both mutant division rates by $1 + s$,

$$A(s) = A_0 + sA_1, \qquad A_1 = \begin{pmatrix} b_x & (1-v) b_y \\ 0 & v b_y \end{pmatrix}.$$

The first-order selective growth coefficient is

$$L_{\text{sel}} = \boldsymbol{w}^{\mathsf{T}} A_1 \boldsymbol{N} = b_x N_x + (1-v) b_y N_y + \rho_y v b_y N_y,$$

and hence

$$\lambda(s) = \frac{L_{\text{sel}}}{W} s + o(s).$$

With $w_x = 1$ and $w_y = \rho_y$, summing the variance contributions in Table 3 gives

$$G = b_x N_x + d_x N_x + \beta N_x N_y (\rho_y - 1)^2 + v b_y N_y \rho_y^2 + (1 - v) b_y N_y + d_y N_y \rho_y^2.$$

Using the equilibrium equations and

$$\beta N_x \rho_y = (1 - v) b_y$$

reduces this expression to

$$G = 2D, \quad \text{where} \quad D = b_x N_x + v b_y N_y \rho_y^2 + (1 - v) b_y N_y \rho_y.$$

The general formulas (40) and (42) therefore give

$$b_{\text{eff}} = \frac{D}{z_0 W}, \qquad s_{\text{eff}} = \frac{z_0 L_{\text{sel}}}{D} s + o(s).$$

For de novo seeding,

$$z_0 = P_x + \rho_y P_y = \frac{L_{\text{sel}}}{b_x N_x + b_y N_y},$$

and consequently

$$\mathcal{R}_s = \frac{L_{\text{sel}}^2}{D(b_x N_x + b_y N_y)}. \tag{63}$$

The explicit result is

$$s_{\text{eff}} = \frac{\left\{ b_x \beta N_x^2 N_y + (1 - v) b_y N_y \left[ \beta N_x N_y + v b_y N_y \right] \right\}^2}{(b_x N_x + b_y N_y) \left\{ b_x \beta^2 N_x^3 N_y^2 + (1 - v)^2 b_y^2 N_y^2 \left[ \beta N_x N_y + v b_y N_y \right] \right\}} s + o(s).$$

#### 2.4 Dimensionless parameterization and the stable domain

We now characterize the evolutionary modifiers derived above. Recall that $\mathcal{R}_N = N_{\text{eff}}/N$ and $s_{\text{eff}} = \mathcal{R}_s s + o(s)$. We shall say that selection is *attenuated* when $0 < \mathcal{R}_s < 1$ and *amplified* when $\mathcal{R}_s > 1$. These terms refer only to the effective selection coefficient: the corresponding change in fixation probability depends jointly on $\mathcal{R}_N$ and $\mathcal{R}_s$.

To characterize the evolutionary modifiers over parameter space, it is convenient to regard the positive equilibrium $(N_x, N_y)$ as specified and use the equilibrium conditions to determine the corresponding death rates. We assume $\beta > 0$ and $0 \le v < 1$, so that $\rho_y$ is well defined; the cases $\beta = 0$ and $v = 1$ must be considered separately or obtained as appropriate limits. Solving the equilibrium equations gives

$$d_x = b_x + \frac{N_y}{N_x} \left[ (1 - v) b_y - \beta N_x \right], \qquad d_y = \beta N_x + v b_y. \tag{64}$$

Introduce the dimensionless quantities $q$, the ratio of the equilibrium compartment sizes, and $\gamma$, the ratio of their per-capita division rates:

$$q = \frac{N_y}{N_x}, \qquad \gamma = \frac{b_x}{b_y} = \frac{r_x}{r_y}, \tag{65}$$

and set

$$h = (1 - v) + v\rho_y$$

for convenience (recall that $\rho_y$ is the relative reproductive value of the $y$-compartment, equation (59)). Using

$$\beta N_x = \frac{(1 - v)b_y}{\rho_y}$$

in equation (64), and dividing by $b_y$, gives

$$\frac{d_x}{b_y} = \gamma + q(1 - v)\left(1 - \frac{1}{\rho_y}\right), \qquad \frac{d_y}{b_y} = v + \frac{1 - v}{\rho_y}. \tag{66}$$

These relations determine which combinations of the dimensionless parameters correspond to non-negative demographic rates.

The reproductive-value ratio $\rho_y$ also indicates whether the microbiome is beneficial or deleterious. For $d_x, d_y > 0$, we call the microbiome beneficial when associated individuals have the larger birth-to-death ratio, $r_y/d_y > r_x/d_x$, and deleterious when the inequality is reversed. Using equation (64),

$$\begin{aligned} b_y d_x - b_x d_y &= \big[(1 - v)b_y - \beta N_x\big](b_x + qb_y) \\ &= \beta N_x(\rho_y - 1)(b_x + qb_y). \end{aligned} \tag{67}$$

Because $b_x/r_x = b_y/r_y = 1 - N/K > 0$, it follows that

$$\operatorname{sign}\left(\frac{r_y}{d_y} - \frac{r_x}{d_x}\right) = \operatorname{sign}(\rho_y - 1). \tag{68}$$

Consequently, the microbiome is beneficial when $\rho_y > 1$, deleterious when $0 < \rho_y < 1$, and demographically neutral when $\rho_y = 1$.

In the notation of equation (65), the probabilities of de novo appearance in the two states are

$$P_x = \frac{\gamma + (1 - v)q}{\gamma + q}, \qquad P_y = \frac{vq}{\gamma + q}, \tag{69}$$

and the two evolutionary modifiers reduce to

$$\mathcal{R}_N = \frac{(\gamma + q)(1 + \rho_y q)}{(1 + q)(\gamma + qh)}, \tag{70}$$

$$\mathcal{R}_s = \frac{(\gamma + qh)^2}{(\gamma + q)(\gamma + \rho_y qh)}. \tag{71}$$

These expressions are meaningful biologically only when the equilibrium is feasible and locally stable. To state these restrictions compactly, set

$$\theta = \frac{\gamma}{q} = \frac{b_x N_x}{b_y N_y}, \qquad \theta_0 = \frac{(1 - v)(1 - \rho_y)}{\rho_y}. \tag{72}$$

Equation (66) then gives $d_x = b_y q(\theta - \theta_0)$, so that $d_x \geq 0$ requires

$$\theta \geq \theta_0. \tag{73}$$

This restriction is automatic when $\rho_y \geq 1$, because then $\theta_0 \leq 0 < \theta$. The second death rate is positive throughout the interior of the parameter domain.

Feasibility alone is not sufficient. Let

$$\chi = \frac{N_x}{K - N} > 0, \tag{74}$$

which measures the strength of density regulation at the equilibrium. On substituting the equilibrium relations into the determinant of the Jacobian derived above, one obtains

$$\frac{\det J_*}{b_y^2} = \frac{q(1-v)}{\rho_y}\left[\chi\{\gamma + qh + v(\rho_y - 1)\} - \frac{(1-v)(\rho_y - 1)}{\rho_y}\right]. \tag{75}$$

Because the trace is negative at a positive equilibrium, local stability is equivalent to

$$\chi\{\gamma + qh + v(\rho_y - 1)\} > \frac{(1-v)(\rho_y - 1)}{\rho_y}. \tag{76}$$

All regime statements below are therefore understood to be restricted to parameters satisfying equations (73) and (76).

##### 2.5 When is the effective population size larger or smaller than the census size?

Subtracting one from equation (70) gives

$$\mathcal{R}_N - 1 = \frac{q(\rho_y - 1)\,[\gamma - v + (1-v)q]}{(1+q)(\gamma + qh)}. \tag{77}$$

The denominator is positive. Hence $\mathcal{R}_N = 1$ on either of two boundaries:

$$\rho_y = 1 \qquad \text{or} \qquad \gamma + (1-v)(1+q) = 1. \tag{78}$$

The second condition has a transparent interpretation in terms of census and de novo distributions, see equations (18) and (19). From equation (69), it is equivalent to

$$P_y^{\text{de novo}} = P_y^{\text{census}}. \tag{79}$$

Thus $N_{\text{eff}} = N$ when the two states have equal reproductive value, or when the probability that a new mutation first appears in state $y$ equals the census fraction of individuals in that state.

For a beneficial microbiome ($\rho_y > 1$), $\mathcal{R}_N > 1$ when $P_y^{\text{de novo}} < P_y^{\text{census}}$, and $\mathcal{R}_N < 1$ when $P_y^{\text{de novo}} > P_y^{\text{census}}$. For a deleterious microbiome ($0 < \rho_y < 1$), these inequalities are reversed. Therefore the sign of the microbiome's demographic effect does not, by itself, determine whether the effective population size is larger or smaller than the census size. Over the stable parameter domain, $\mathcal{R}_N$ has no finite universal upper bound and can approach zero in limiting parameter regimes; the equality surfaces in equation (78), rather than the beneficial–deleterious boundary alone, separate the two regimes.

##### 2.6 When is effective selection amplified or attenuated?

From equation (71),

$$\mathcal{R}_s - 1 = \frac{q(1-\rho_y)\,\{\gamma[1 - v(2-\rho_y)] + (1-v)qh\}}{(\gamma + q)(\gamma + \rho_y qh)}. \tag{80}$$

The denominator is again positive. If $\rho_y > 1$, we have

$$1 - v(2 - \rho_y) = (1 - v) + v(\rho_y - 1) > 0,$$

the expression in curly brackets in (80) is positive, and therefore

$$\rho_y > 1 \quad \Longrightarrow \quad 0 < \mathcal{R}_s < 1. \tag{81}$$

Thus a beneficial microbiome always attenuates the effective selection coefficient in this model. At $\rho_y = 1$, one has $\mathcal{R}_s = 1$.

For a deleterious microbiome, $0 < \rho_y < 1$, two cases arise. If

$$v \leq \frac{1}{2 - \rho_y}, \tag{82}$$

the expression in curly brackets in equation (80) is positive for every $\gamma, q > 0$, and hence $\mathcal{R}_s > 1$. Selection is therefore amplified throughout the feasible and stable part of this parameter region.

If instead $v > 1/(2 - \rho_y)$, define

$$\theta_s = \frac{(1 - v)h}{v(2 - \rho_y) - 1}. \tag{83}$$

Then

$$\mathcal{R}_s > 1 \quad \Longleftrightarrow \quad \theta < \theta_s, \qquad \mathcal{R}_s < 1 \quad \Longleftrightarrow \quad \theta > \theta_s, \tag{84}$$

with $\mathcal{R}_s = 1$ at $\theta = \theta_s$. Because feasibility also requires $\theta \geq \theta_0$, a feasible amplification interval exists precisely when

$$\theta_s > \theta_0 \quad \Longleftrightarrow \quad v(1 - \rho_y) < \frac{1}{2}. \tag{85}$$

The actual interval must, in addition, satisfy the stability condition (76).

The same boundary can be expressed in terms of where new mutations arise. Since $P_y^{\text{de novo}} = v/(1 + \theta)$, equation (84) is equivalent to

$$\mathcal{R}_s > 1 \quad \Longleftrightarrow \quad P_y^{\text{de novo}} > P_y^*, \qquad P_y^* = 1 - \frac{1 - v}{v(1 - \rho_y)}, \tag{86}$$

in the regime $v > 1/(2 - \rho_y)$. Thus, for a deleterious microbiome with sufficiently faithful vertical transmission, whether selection is amplified or attenuated depends on the relative contribution of the two states to the production of new mutations.

#### 2.7 Maximum amplification of effective selection

The preceding inequalities identify the regimes in which $\mathcal{R}_s$ is above or below one. They also permit a global bound on its magnitude. Writing equation (71) in terms of $\theta = \gamma/q$ gives

$$\mathcal{R}_s(\theta) = \frac{(\theta + h)^2}{(\theta + 1)(\theta + \rho_y h)}, \qquad \theta \geq \theta_0. \tag{87}$$

For $0 < \rho_y < 1$,

$$\frac{d}{d\theta} \log \mathcal{R}_s = -\frac{(1 - \rho_y)\,\{[1 - v(2 - \rho_y)]\theta + h(1 - v\rho_y)\}}{(\theta + h)(\theta + 1)(\theta + \rho_y h)}. \tag{88}$$

Table 4: Parameter sets illustrating large increases and decreases in effective population size. The microbiome is classified as beneficial when $r_y/d_y > r_x/d_x$ and as deleterious when $r_y/d_y < r_x/d_x$.

| $v$ | $r_x$ | $r_y$ | $d_x$ | $d_y$ | $\beta$ | $K$ | Microbiome | $N_x$ | $N_y$ | $\mathcal{R}_N$ |
|---|---|---|---|---|---|---|---|---|---|---|
| 0 | 1 | $5.0 \times 10^{-2}$ | $5.249874937 \times 10^{-1}$ | $1.250625 \times 10^{-5}$ | $2.501250625 \times 10^{-8}$ | $2.0 \times 10^{3}$ | Beneficial | $5.00 \times 10^{2}$ | $5.00 \times 10^{2}$ | $1.00 \times 10^{3}$ |
| 0.9 | 1 | $2.50437922403 \times 10^{3}$ | $1.12528154 \times 10^{-4}$ | $2.25394142697 \times 10^{-1}$ | $6.270347312 \times 10^{-16}$ | $2.000200020 \times 10^{7}$ | Beneficial | $1.999 \times 10^{7}$ | $1.000 \times 10^{4}$ | $1.00 \times 10^{-3}$ |
| $v \to 1^-$ | 1 | $1.999 \times 10^{3}$ | $5.0 \times 10^{-4}$ | $1.9989995 \times 10^{3}$ | $2.4975 \times 10^{-7}$ | $1.6 \times 10^{10}$ | Deleterious | $4.00 \times 10^{6}$ | $4.00 \times 10^{6}$ | $1.00 \times 10^{3}$ |
| $10^{-6}$ | 1 | $2.501250625 \times 10^{-7}$ | $1.0 \times 10^{-8}$ | $5.0 \times 10^{-9}$ | $5.0 \times 10^{-12}$ | $2.000020000 \times 10^{6}$ | Deleterious | $1.00 \times 10^{3}$ | $1.999 \times 10^{6}$ | $1.00 \times 10^{-3}$ |

If this derivative vanishes in the feasible interval, the stationary point is a minimum. Because $\mathcal{R}_s(\theta) \to 1$ as $\theta \to \infty$, the maximum over the feasible interval is therefore attained either at $\theta = \theta_0$ or in the limit $\theta \to \infty$. At the feasibility boundary, $\theta_0 = h/\rho_y - 1$, and

$$\mathcal{R}_s(\theta_0) = \frac{[h(1+\rho_y) - \rho_y]^2}{h\,[h(1+\rho_y^2) - \rho_y]}, \qquad \rho_y \le h \le 1. \tag{89}$$

As a function of $h$, this expression has no interior maximum; its endpoint values are 1 at $h = \rho_y$ and $1/(1 - \rho_y + \rho_y^2)$ at $h = 1$. Maximizing the latter expression over $0 < \rho_y < 1$ gives

$$\mathcal{R}_s \le \frac{4}{3}. \tag{90}$$

Equality is realized at

$$v = 0, \qquad \rho_y = \frac{1}{2}, \qquad \theta = \theta_0 = 1, \tag{91}$$

or, equivalently, when $d_x = 0$, $b_x N_x = b_y N_y$, and $\beta N_x = 2b_y$. This point satisfies the local-stability condition (76) for every $\chi > 0$. If strictly positive $d_x$ or $v$ is required, 4/3 is a supremum approached as the corresponding parameter tends to zero rather than an attained maximum.

#### 2.8 Summary of the regime boundaries

The behavior of the two evolutionary modifiers is organized by five boundaries. The surface $\rho_y = 1$ separates beneficial from deleterious microbiomes. The additional surface $\gamma + (1-v)(1+q) = 1$ determines whether $\mathcal{R}_N$ is above or below one. For a deleterious microbiome with $v > 1/(2-\rho_y)$, the surface $\theta = \theta_s$ separates amplification from attenuation of the effective selection coefficient. Finally, $\theta = \theta_0$ is the feasibility boundary $d_x = 0$, and equality in equation (76) is the local-stability boundary. The algebraic regimes for $\mathcal{R}_N$ and $\mathcal{R}_s$ must always be intersected with the feasible and stable side of the last two boundaries.

The principal conclusions are therefore distinct. A beneficial microbiome always gives $\mathcal{R}_s < 1$, but it may give either $\mathcal{R}_N < 1$ or $\mathcal{R}_N > 1$. A deleterious microbiome may also place $\mathcal{R}_N$ on either side of one, whereas $\mathcal{R}_s$ may be either above or below one depending on vertical transmission and the relative production of new mutations in the two states. Across the entire stable domain, amplification of the effective selection coefficient is bounded by $\mathcal{R}_s \le 4/3$, while no analogous finite universal upper bound exists for $\mathcal{R}_N$.

#### 2.9 Numerical examples

**Potential magnitude of changes in effective population size.** The absence of a finite universal upper bound on $\mathcal{R}_N$, and the fact that its infimum is zero, allow the effective population size to differ greatly from the census population size. Table 4 provides illustrative parameter sets for which $N_{\text{eff}}$ is either $10^3$ times larger or $10^3$ times smaller than $N$. Both outcomes can occur

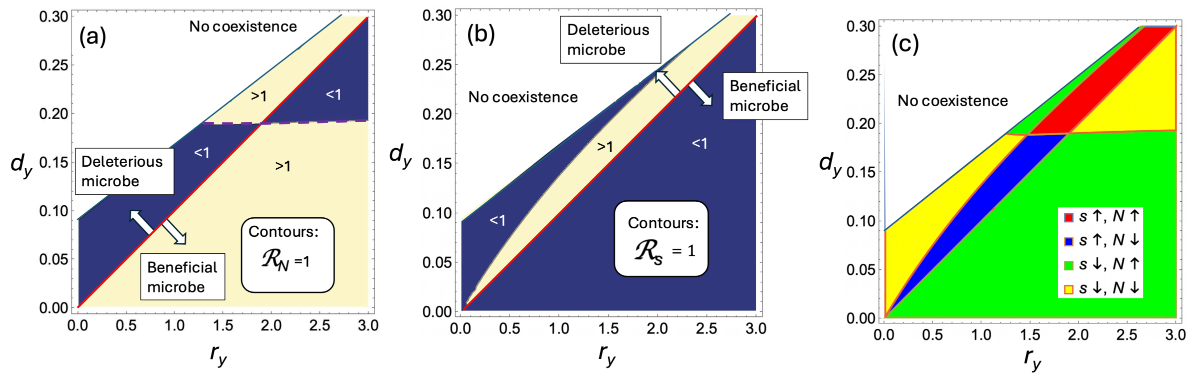

Figure 1: The effects of newborn allocation and microbiome type. (a) Effective population size: the regions where quantity $\mathcal{R}_N > 1$ (yellow) and $\mathcal{R}_N < 1$ (blue) are plotted with the axes that are division and death rate of microbiome-associated individuals, $r_y$ and $d_y$. White area: no stable coexistence solution. (b) Effective selection coefficient: the same plot for $\mathcal{R}_s$. (c) The combination of the two quantities reveals the existence of four different regions (see the legend). The rest of the parameters are $r_x = 1, d_x = 0.1, v = 0.8, \beta = 10^{-4}, K = 10^3$. Two domain boundaries are of note: (1) in panels (a) and (b), above the red line, the microbiome is deleterious and below it, it is beneficial; (2) in panel (a), above the purple dashed line, a microbiome-associated individual is created with the probability greater than its census fraction, and below it, less than its census fraction.

with either a beneficial or a deleterious microbiome. These parameter sets are constructed examples, rather than estimates for particular biological systems, but they demonstrate that microbiome structure can in principle alter the strength of genetic drift by several orders of magnitude. They also reinforce that the direction of this change cannot be inferred from whether the microbiome is beneficial or deleterious.

**The effects of newborn allocation and microbiome type.** Figure 1 illustrates how the demographic effect of the microbiome interacts with the allocation of newborns between the two compartments. The red line separates beneficial from deleterious microbiomes, whereas the dashed purple line marks the condition

$$P_y^{\text{de novo}} = P_y^{\text{census}}.$$

The latter boundary is important because mutations arise in newborns and therefore need not be distributed among compartments in proportion to their census abundances. For a beneficial microbiome, the associated compartment has the larger reproductive value, and $\mathcal{R}_N > 1$ when this compartment is underrepresented among new mutant origins, $P_y^{\text{de novo}} < P_y^{\text{census}}$; the inequality is reversed for a deleterious microbiome. Figure 1(b) also shows that a beneficial microbiome always gives $\mathcal{R}_s < 1$, whereas a deleterious microbiome can either amplify or attenuate the effective selection coefficient. Combining these effects produces the four regimes shown in Figure 1(c), demonstrating that the changes in effective population size and effective selection strength are not determined by one another.

**The effect of the mode of transmission.** Figure 2 shows that vertical and infection-type horizontal transmission can have nonmonotonic and strongly context-dependent effects on both evolutionary modifiers. Changing either $\beta$ or $v$ alters not only the rate of movement between compartments, but also the equilibrium composition, the distribution of new mutant origins, and the reproductive values of the two states. Consequently, increasing transmission does not produce a universal increase or decrease in either $\mathcal{R}_N$ or $\mathcal{R}_s$. The two beneficial examples in Figures 2(a,b)

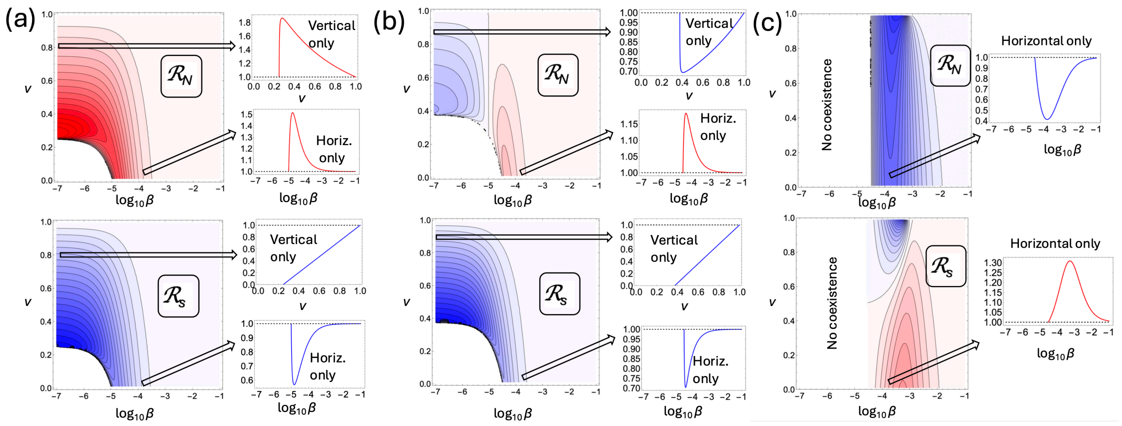

Figure 2: Evolutionary modifies. $\mathcal{R}_N$ (top row) and $\mathcal{R}_s$ (bottom row) for different values of $\beta$ and $v$. (a) Beneficial microbiome with $d_y < d_x$ ($r_y = 2, d_y = 0.01$); (b) Beneficial microbiome with $d_y > d_x$ ($r_y = 4, d_y = 0.03$); (c) Deleterious microbiome ($r_y = 0.075, d_y = 0.03$). For each case, a heatplot of $\mathcal{R}_N$ (top) and $\mathcal{R}_s$ (bottom) is presented, where values greater than one are shown in shades of red and values smaller than one in shades of blue. For purely horizontal transmission ($v = 0$), the corresponding quantity is shown as a function of $\log_{10}\beta$. For purely vertical transmission ($\beta \to 0$), the corresponding quantity is shown as a function of $v$. The rest of the parameters are $r_x = 1, d_x = 0.02, K = 10^3$.

already produce qualitatively different patterns for $\mathcal{R}_N$, although both satisfy $\mathcal{R}_s < 1$, as predicted above. In the deleterious example in Figure 2(c), the effective population size is reduced while the effective selection coefficient is amplified over much of the parameter range. The one-dimensional sections show the purely infection-type limit, $v = 0$, and the purely vertical limit, $\beta \to 0$. Their different behavior emphasizes that vertical and horizontal transmission are not interchangeable: their evolutionary effects depend on how transmission is combined with the demographic rates of the two compartments.

### 3 Examples of specific biological systems

In this section we apply our model to several biological examples, which are summarized in table 5. In this section the examples are all two-compartment systems, which consist of a microbiome-unassociated and microbiome-associated hosts. Application of the model to more than two compartments is considered in section 4.

**Fold change in fixation probability.** To quantify the combined effects of the microbiome on drift and selection, we compare the predicted fixation probability with the standard unstructured prediction based on the census population size $N$ and selection coefficient $s$. We define the fold change

$$\mathcal{F}(s) = \frac{P_{\text{fix}}(N\mathcal{R}_N, \mathcal{R}_s s)}{P_{\text{fix}}(N, s)}. \tag{92}$$

For a deleterious mutant, $s < 0$, the condition $\mathcal{F} > 1$ means that the mutant fixes more frequently than predicted by the unstructured model and therefore indicates suppression of selection against deleterious mutations. For an advantageous mutant, $s > 0$, the condition $\mathcal{F} < 1$ means that the mutant fixes less frequently than predicted by the unstructured model and therefore indicates suppression of positive selection. At neutrality,

$$\mathcal{F}(0) = \frac{1/N_{\text{eff}}}{1/N} = \frac{1}{\mathcal{R}_N}, \tag{93}$$

which need not equal one because the neutral fixation probability is determined by $N_{\text{eff}}$, rather than by the census population size $N$.

| Host | Microbe | Beneficial or deleterious | Transmission type | Notes |
|---|---|---|---|---|
| *Paramecium caudatum* | *Holospora undulata* | Deleterious | V & I | Two evolutionary treatments: Front/UP and Core/DOWN |
| *Drosophila melanogaster* | Lp & Lb | Deleterious | V & E | |
| *Drosophila melanogaster* | Ap, At, Ao | Beneficial | V & E | |
| *Tetrahymena utriculariae* | *Micractinium tetrahymenae* | Deleterious | V & E | Density-dependent birth and/or death |
| *Aphis fabae* | *Hamiltonella defensa* | Deleterious or beneficial | V & I | Different rates of attack define microbe fitness |
| *Agrostis hyemalis* | *Epichloë amarillans* | Beneficial | V & I | Ambient and irrigated conditions |

Table 5: Here "V" stands for vertical transmission, "I" for infection-type transmission, and "E" for environmental transmission.

##### 3.1 *Paramecium caudatum–Holospora undulata*

**Biological system and experiments.** *Paramecium caudatum* is a freshwater ciliate, and *Holospora undulata* is an obligately intracellular bacterial parasite of its germline micronucleus. The parasite is transmitted vertically when an infected host divides and horizontally when susceptible hosts ingest infectious forms released from infected cells. Production of infectious forms can reduce host division and survival, generating a trade-off between horizontal transmission and virulence. Nørgaard *et al.* [6] experimentally evolved replicate parasite populations either at the expanding front of a two-patch range expansion or in the range core. The two conditions corresponding to the front and the core of the colony are referred to as Front/UP and Core/DOWN, respectively [6].

**Ecological model.** The authors of study [6] fitted a continuous-time epidemiological model to the densities of uninfected and infected hosts. We will use their results to investigate the evolutionary consequences of this parasite. Before we proceed, we summarize the model, the parameterization procedure, and its results. We will use notations that are consistent with those of the present study. Let $x$ and $y$ denote uninfected and infected hosts and write the equations for the mean quantities as

$$\dot{x} = \frac{r_x}{1+\eta(x+y)}x + (1-v)\frac{r_y}{1+\eta(x+y)}y - d\,x - \beta xy, \tag{94}$$

$$\dot{y} = \beta xy + v\frac{r_y}{1+\eta(x+y)}y - d\,y. \tag{95}$$

Here $r_x$ is the maximum division rate of an uninfected host, $r_y$ is the corresponding rate for an infected host, $d$ is the common death rate, $\eta$ controls density dependence, $\beta$ is the horizontal-transmission coefficient, and $v$ is vertical-transmission fidelity. The source paper [6] also fitted a latency parameter, $\tau$, but it only determined when horizontal transmission began after experimental inoculation, and is not used in the post-latency equilibrium calculation.

For each posterior parameter set, we solve equations (94-95) for its positive equilibrium $(N_x, N_y)$. The rare-mutant calculation then uses the equilibrium division rates

$$b_x = \frac{r_x}{1+\eta(N_x+N_y)}, \qquad b_y = \frac{r_y}{1+\eta(N_x+N_y)}, \tag{96}$$

together with $d_x = d_y = d$, $\beta$, and $v$. Thus the fitted model can be used directly to explore the evolutionary consequences of the parasite.

**Parameter values, uncertainty and limitations.** Nørgaard *et al.* first fitted $r_x$, $d$, and $\eta$ to growth curves from uninfected populations. These fitted distributions were then used as priors when the full model was fitted to infected and uninfected density time series from days 6–13 after inoculation. The full model was fitted separately for each host genotype and parasite-selection treatment using Bayesian inference. We use the fit for host genotype 63D, the genotype on which the experimental-evolution lines were maintained, and propagate complete posterior draws rather than combining independently sampled marginal estimates.

The reported uncertainty comes from the joint Bayesian fit of the demographic and transmission parameters and is propagated through the nonlinear equilibrium and fixation calculations. Posterior draws are reported in [6]. All 30,000 Front/UP draws and 29,914 of 30,000 Core/DOWN draws produced a positive equilibrium. The marginal values are reported in Table 6.

The fitted model assumes a common death rate for infected and uninfected hosts and represents virulence through reduced division, although separate assays indicated that infection can also reduce survival. Including an additional infection-specific death effect can be performed e.g. using the method similar to that we describe in Section 3.3. The vertical-transmission prior was restricted to $0.5 \le v \le 1$, which contributes to uncertainty in the balance between vertical and horizontal transmission.

| Parameter | Front/UP | Core/DOWN |
|---|---|---|
| $r_x$ (d$^{-1}$) | 4.56 | 4.33 |
| $r_y$ (d$^{-1}$) | 3.50 | 0.478 |
| $d$ (d$^{-1}$) | 0.994 $[0.622, 1.566]$ | 1.002 $[0.720, 1.488]$ |
| $\eta$ (ml cell$^{-1}$) | 0.0126 | 0.0135 |
| $\beta$ (ml cell$^{-1}$ d$^{-1}$) | 0.0040 $[0.0022, 0.0070]$ | 0.0094 $[0.0043, 0.0148]$ |
| $v$ | 0.844 $[0.541, 0.990]$ | 0.694 $[0.508, 0.981]$ |

Table 6: Parameter values for the *Paramecium caudatum–Holospora undulata* examples used here. Values in brackets are 95% posterior intervals where available; rows without intervals give the central values used for illustration. The ecological parameters are expressed in the original Beverton–Holt parameterization.

**Effect on evolution.** We used the methodology developed in this paper to calculate the modifiers $\mathcal{R}_N$ and $\mathcal{R}_s$ for this parameterized system. In panel (a) of figures 3 and 4, we present an approximation of the probability distributions of $\mathcal{R}_N$ and $\mathcal{R}_s$ for the Front/UP and Core/DOWN

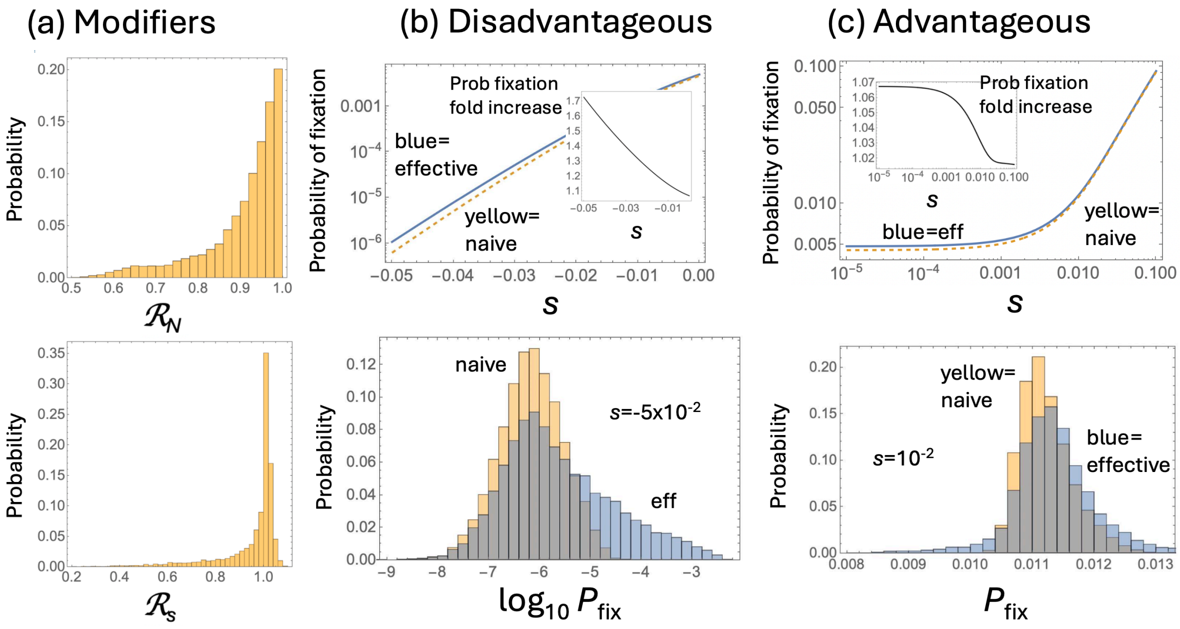

Figure 3: Modifications to selection in Paramecium caudatum infected with the bacterial parasite Holospora undulata: Front/UP. (a) The modifiers $\mathcal{R}_s$ and $\mathcal{R}_N$ are calculated using the posteriors for all the parameters in [6]. (b) Disadvantageous mutants: The probability of mutant fixation for disadvantageous mutants is plotted as a function of $s$, with $P_{\text{fix}}(N\mathcal{R}_N, s\mathcal{R}_s)$ in blue and $P_{\text{fix}}(N, s)$ in dashed yellow (top panel). The inset shows the probability ratio, $\mathcal{F}$. The bottom panel shows the estimated probability distributions of $P_{\text{fix}}(N\mathcal{R}_N, s\mathcal{R}_s)$ (blue) and $P_{\text{fix}}(N, s)$ (yellow) with a fixed value $s = -5 \times 10^{-2}$. (c) Same as (b) but for advantageous mutants; the fixed value in the bottom panel is $s = 10^{-2}$. The rest of the parameters are given in table 6, unless otherwise noted.

conditions, respectively. In order to obtain these panels we used the posteriors of all the parameters provided in [6] and used our method with the density dependent reproduction rates given by equation (96). We can see that the two treatments have different evolutionary consequences. Core/DOWN parasites are more virulent and transmit horizontally at a higher rate, producing a substantially smaller effective population size, $\mathcal{R}_N = 0.687$, while Front/UP parasites produce a smaller reduction in effective population size, $\mathcal{R}_N = 0.939$. The effect on the selective coefficient in the Core/DOWN condition is more modest, and we observe essentially no change in selection at the posterior median, for the Front/UP condition.

Panels (b) and (c) of figures 3 and 4 study the effect of the bacterial parasite on the probability of mutant host fixation, as predicted by our theory. The top panels plot the fixation probability calculated using the effective quantities, $P_{\text{fix}}(N\mathcal{R}_N, s\mathcal{R}_s)$ (blue) and compare it with the naive estimate, $P_{\text{fix}}(N, s)$ (yellow dashed line). The top panels of figure 3(b,c) assume the equilibrium population $N_x = 53.0, N_y = 167.4, N = 220.4$ (calculated using the central values of all the parameters, table 6). We observe a very modest increase of the fixation probability for disadvantageous mutants and an even smaller difference for advantageous mutants. The bottom panels fix a specific value of $s$ (with $s < 0$ in bottom panel 3(b) and $s > 0$ in bottom panel 3(c)) and show how the probability of fixation is distributed, again using the posteriors of all the parameters provided in [6] and comparing $P_{\text{fix}}(N\mathcal{R}_N, s\mathcal{R}_s)$ (blue) with $P_{\text{fix}}(N, s)$ (yellow). We conclude that for the Front/UP condition (figure 3) there is only a very small difference between the probability of fixation compared to that expected from the usual formula for the unstructured population.

In contrast to that, for the Core/DOWN treatment (figure 4, panels (b) and (c)) we observe

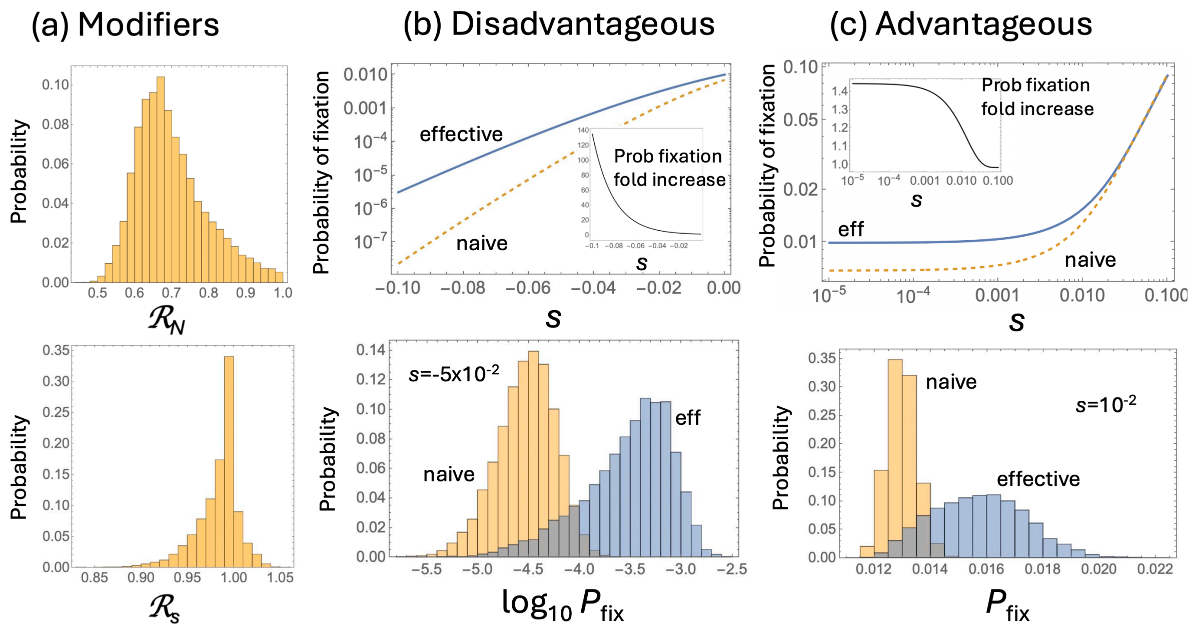

Figure 4: Modifications to selection in Paramecium caudatum infected with the bacterial parasite Holospora undulata: Core/DOWN. (a) The modifiers $\mathcal{R}_s$ and $\mathcal{R}_N$ are calculated using the posteriors for all the parameters in [6]. (b) Disadvantageous mutants: The probability of mutant fixation for disadvantageous mutants is plotted as a function of $s$, with $P_{\text{fix}}(N\mathcal{R}_N, s\mathcal{R}_s)$ in blue and $P_{\text{fix}}(N, s)$ in dashed yellow (top panel). The inset shows the probability ratio, $\mathcal{F}$. The bottom panel shows the estimated probability distributions of $P_{\text{fix}}(N\mathcal{R}_N, s\mathcal{R}_s)$ (blue) and $P_{\text{fix}}(N, s)$ (yellow) with a fixed value $s = -5 \times 10^{-2}$. (c) Same as (b) but for advantageous mutants; the fixed value in the bottom panel is $s = 10^{-2}$. The rest of the parameters are given in table 6, unless otherwise noted.

a very significant effect. The top panels 4(b,c) use $N_x = 94.7, N_y = 51.3$, and $N = 146.0$ to plot the probability of fixation as a function of $s$. For disadvantageous mutants, as $s$ increases, the probability of fixation estimated with the effective modifiers becomes orders of magnitude larger than that obtained using the census population size. For advantageous mutants, the probability of fixation is also increased (by up to about 40%). The bottom panels in figure 4(b,c) also show a significant difference in the probability distributions of $P_{\text{fix}}(N\mathcal{R}_N, s\mathcal{R}_s)$ (blue) with $P_{\text{fix}}(N, s)$ (yellow) with fixed $s$ values.

Given that it is the value of $\mathcal{R}_N$ that is significantly different from one (with $\mathcal{R}_s$ only slightly deviating from unity), see figure 4(a), we conclude that the fixation effects observed in panels 4(b,c) are driven mainly by increased drift.

It is interesting that for disadvantageous mutants we observe $\mathcal{F} >> 1$, that is, strong suppression of selection, and for advantageous mutants, again $\mathcal{F} > 1$. For both disadvantageous and advantageous mutants the presence of bacterial parasite results in an increase of the fixation probability compared to its non-modified value in the absence of the microbe. So in a sense, the effect is the opposite: selection is suppressed for disadvantageous mutants (i.e., the probability of fixation increases) and enhanced for advantageous mutants (again, the probability of fixation increases).

#### 3.2 *Drosophila melanogaster*

**Biological system and experiments.** The fruit fly *Drosophila melanogaster* carries a gut microbiome in which lactic-acid bacteria and acetic-acid bacteria can alter host development, reproduction, and adult survival. Gould *et al.* isolated five bacterial species found in their laboratory flies, generated germ-free flies by surface-sterilizing embryos, and inoculated adult flies through their food with every possible combination of these species [7]. The five species were *Lactobacillus plantarum* (Lp), *Lactobacillus brevis* (Lb), *Acetobacter pasteurianus* (Ap), *Acetobacter tropicalis* (At), and *Acetobacter orientalis* (Ao). For each microbial treatment, the authors measured lifetime fecundity, adult survival, offspring development time, and bacterial abundance, thereby allowing the demographic effects of individual species and multispecies communities to be compared with those of germ-free flies.

**Model and parameterization.** We use the germ-free and five single-species treatments to parameterize five separate two-state models. In each model, $x$ denotes germ-free females and $y$ denotes females associated with one bacterial species; the multispecies treatments are not used in this parameterization. Because the flies acquired bacteria from inoculated food, we replace the host-to-host acquisition term $\beta xy$ by environmental acquisition at rate $\lambda x$. The mean dynamics in the absence of mutations are

$$\dot{x} = r_x x \left(1 - \frac{x+y}{K}\right) + (1-v) r_y y \left(1 - \frac{x+y}{K}\right) - d_x x - \lambda x, \tag{97}$$

$$\dot{y} = \lambda x + v r_y y \left(1 - \frac{x+y}{K}\right) - d_y y, \tag{98}$$

where $\lambda$ is the environmental-acquisition rate and $v$ is the probability that the offspring of an associated female retains the microbiome. As in the general model, a host mutation multiplies both reproduction rates by $1+s$, while leaving death and transmission rates unchanged.

For each microbial state $i$, we obtained a coarse-grained death rate and reproduction rate from the published mean time to death, $L_i$, and total lifetime fecundity, $F_i$:

$$d_i = \frac{1}{L_i}, \qquad r_i = \frac{F_i}{2L_i}. \tag{99}$$

The factor 1/2 converts total offspring to daughters under an assumed 1:1 sex ratio, because the model tracks females. We set $(r_x, d_x)$ to the germ-free values and $(r_y, d_y)$ to the values for the focal single-species association. The resulting rates, in $d^{-1}$, are listed in Table 7.

| State | $r_i$ ($d^{-1}$) | $d_i$ ($d^{-1}$) |
|---|---|---|
| Germ-free | 1.225 [1.112, 1.344] | 0.01878 [0.01780, 0.01984] |
| Lp | 1.034 [0.828, 1.248] | 0.02144 [0.02017, 0.02291] |
| Lb | 0.821 [0.530, 1.119] | 0.01916 [0.01800, 0.02050] |
| Ap | 1.590 [1.437, 1.754] | 0.01922 [0.01818, 0.02039] |
| At | 1.878 [1.783, 1.979] | 0.02070 [0.01989, 0.02162] |
| Ao | 2.220 [2.098, 2.353] | 0.02317 [0.02199, 0.02444] |

Table 7: Demographic parameters for the *Drosophila* examples. Brackets give approximate 95% propagated uncertainty intervals. The reproduction and death rates are effective age-independent rates constructed from lifetime fecundity and mean adult lifespan, respectively.

**Uncertainty and limitations.** Neither $\lambda$ nor $v$ is identifiable from the experiment. Adults were experimentally inoculated and repeatedly exposed to bacteria in their food, but the study did not measure time to colonization, an acquisition dose response, or natural vertical transmission. Accordingly, parameters $\lambda$ and $v$ will be varied to explore sensitivity.

Uncertainty in the demographic rates was propagated from the published standard errors of $F_i$ and $L_i$. For each treatment, we generated $5 \times 10^5$ independent normal draws of these two quantities, recalculated equation (99), and used the 2.5th and 97.5th percentiles as approximate 95% intervals. These intervals are not formal confidence intervals: the covariance between lifetime fecundity and lifespan, as well as the complete replicate and batch structure, was unavailable. The strongest reproductive effects (the reduction for Lb and increases for At and Ao) are nevertheless well separated from the germ-free value; the Lp reproduction interval and the Ap and Lb death-rate intervals overlap the corresponding germ-free intervals. There is also structural uncertainty in reducing age-dependent *Drosophila* demography to constant rates. Equation (99) preserves mean adult lifetime and expected lifetime production of daughters, but it does not reproduce the observed survival-time distribution, the approximately ten-day developmental delay, or the absolute population growth rate.

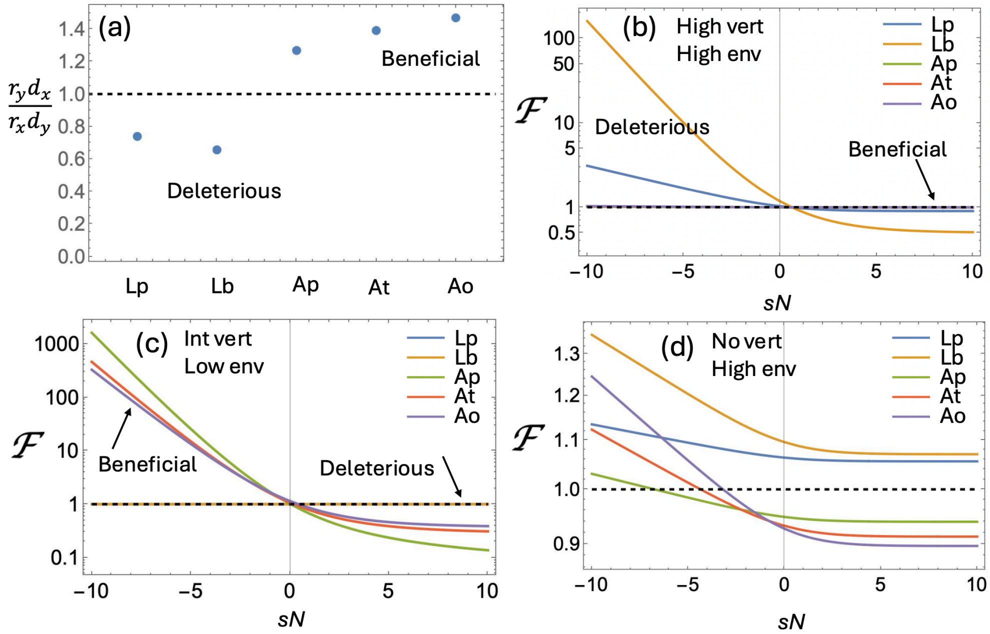

Figure 5: Drosophila. (a) The quantity $\frac{r_y d_x}{d_y r_x}$ shown for each of the five bacterial species, using the data in table 7. In (b-d), the fold-change in fixation probability, $\mathcal{F}$, is plotted for all five bacteria as a function of $sN$, for specific values of vertical and environmental transmission rates: (b) high vertical and high environmental transmission rates, $v = 0.99, \lambda = 10^{-2}$; (c) intermediate vertical and low environmental transmission rates, $v = 0.8, \lambda = 10^{-6}$; (d) Essentially no vertical and high environmental transmission rates, $v = 10^{-2}, \lambda = 10^{-2}$. Other parameters are given in table 7 (the central values are used).

**Effect on evolution.** Out of the five bacteria studied here, two (Lp and Lb) are deleterious to the host, and three (Ap, At, and Ao) are beneficial, see figure 5(a). The two groups are expected to show different patterns of behavior in the context of evolution.

The model parameters that are available for this system (see table 7) do not include the rates of vertical ($v$) or environmental ($\lambda$) transmission. Therefore, to investigate possible effects of the microbes on the evolutionary modifiers, we varied $v$ and $\lambda$ over a large range, see figure 6. In this figure, we used shades of red for the regions where the quantity of interest is greater than one, and shades of blue where it is smaller than one. For example, in the bottom left panel that shows $\mathcal{R}_N$

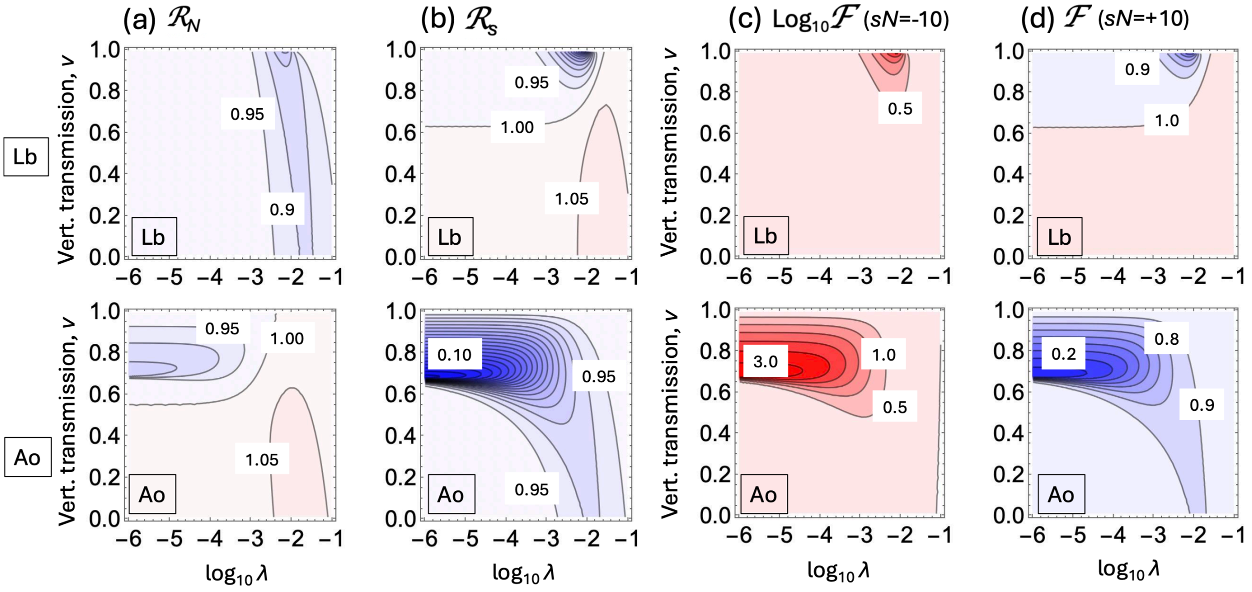

Figure 6: Drosophila. Evolutionary modifiers and the factor $\mathcal{F}$ are plotted as heat-plots as function of $\lambda$ (please note the logarithmic scale) and $v$. Regions $> 1$ ($< 1$) are colored in shades of red (blue). Top row corresponds to bacterium Lb (deleterious) and bottom row to bacterium Ao (beneficial). (a) $\mathcal{R}_N$ is shown with a step-size in contours $\Delta = 0.05$; (b) $\mathcal{R}_s$, $\Delta = 0.05$, (c) for $Ns = -10$, we plot $\log_{10} \mathcal{F}$ with $\Delta = 0.5$, (d) for $Ns = +10$, we plot $\mathcal{F}$ with $\Delta = 0.1$. Other parameters are given in table 7 (the central values are used).

for Ao, in the blue-colored regions we have $N_{\text{eff}} < N$ and in red-colored regions we have $N_{\text{eff}} > N$. For this figure, we chose one deleterious (Lb, top row) and one beneficial (Ao, bottom row) species of bacteria. The other species show qualitatively similar behavior.

In panel 6(a) we show a heat-plot of the modifier $\mathcal{R}_N$. For the deleterious bacteria (Lb) we observe a modest decrease of the effective population around $\lambda \sim 10^{-2}$. For Advantageous bacteria (Ao) both increase and decrease of the effective population is observed. In panel (b), $\mathcal{R}_s$ is shown, and the behavior is again different for deleterious and beneficial microbes: the effective selection coefficient might be slightly increased or decreased for deleterious microbes (the top panel) and we observe a strong decrease in a large parameter region for beneficial microbes (the bottom panel).

To show how these changes translate into the changes in fixation probability, we explored the behavior of the ratio $\mathcal{F}$ for disadvantageous mutants (panel 6(c), $sN = -10$) and advantageous mutants (panel 6(d), $sN = 10$). As expected, the behavior is different between deleterious and beneficial microbes. For deleterious microbes, the effect is modest for most of the parameter space, but for advantageous mutants (top panel (d)) we observe both suppression of selection (the blue region) and amplification of selection (the red region). For beneficial microbes, we only have suppression of selection, but the effect can be very strong. For example, $\mathcal{F}$ could be as high as $10^3$ for disadvantageous mutants (that is, they are a thousand times more likely to fixate compared to the homogeneous population prediction), and $\mathcal{F}$ could be as low as 0.2 for advantageous mutants (that is, the probability of fixation is reduced by 80% compared to the homogeneous population prediction).

In figure 5(b-d) we used three specific $(\lambda, v)$ pairs and plotted the ratio $\mathcal{F}$ as a function of $Ns$ for all five bacteria. Panel 5(b) shows the high vertical transmission and high environmental transmission case, where beneficial microbes do not change the probability of fixation, while deleterious microbes cause strong selection suppression. In panel 5(c) we have intermediate vertical and low environmental transmission, where deleterious microbes make no change on evolution but beneficial microbes cause very strong suppression of selection. Finally, in panel 5(d) (no vertical transmission, high environmental transmission) the effect is modest for both types of bacteria, but we observe an approximately 10% increase in advantageous mutant fixation probability for deleterious microbes.

##### 3.3 *Tetrahymena utriculariae–Micractinium tetrahymenae*

**Biological system and experiments.** *Tetrahymena utriculariae* is a freshwater ciliate that naturally carries intracellular green algae. The symbiont was initially reported as *Micractinium* sp. [8] and was subsequently described as *Micractinium tetrahymenae* [9]. The association is facultative and environmentally dependent: algae are transmitted to daughter cells during host division, symbiotic cultures can lose their algae under oxic, bacteria-rich conditions, and aposymbiotic ciliates can reacquire algae from the environment. Kamal *et al.* found that symbiosis benefits the host under illuminated hypoxia but substantially reduces host growth under the oxic, bacteria-fed conditions used for the present parameterization [10].

**Model and parameterization.** Let $x(t)$ and $y(t)$ denote the densities of aposymbiotic (i.e. without the algae) and symbiotic hosts. To shed light on mechanisms of transmission, the authors of [8] added isolated Micractinium cells to the aposymbiotic ciliates. After 44 days, approximately 80% of the ciliates contained algae when bacterial food was also supplied, and approximately 50% contained algae when only the algae were supplied. Therefore, aposymbiotic ciliates acquired algae placed in the external medium, without requiring contact with an already symbiotic ciliate. This experiment demonstrates the presence of environmental acquisition (term $\lambda x$), which we used in the model below. In addition, we assume vertical transmission at rate $v$, where an offspring of a symbiotic parent retains the alga with probability $v$.

To model the processes of births and deaths we note that (i) these could not be measured separately in the experiment and (ii) evidence from a related species, *Tetrahymena pyriformis*, shows the possibility of density-dependent births and/or deaths [11]. As population density increased, they observed reduced cell-division rates and inferred increased death rates, indicating that density regulation can operate through both demographic processes. To incorporate this possibility in the model, define the per-capita birth and death rates of a host in state $i \in \{x, y\}$ as

$$B_i(x,y) = r_i\left[1-(1-\phi)\frac{x+y}{K}\right], \qquad D_i(x,y) = d_i + \phi r_i\frac{x+y}{K}, \qquad 0 \le \phi \le 1. \tag{100}$$

The resident equations are

$$\dot{x} = \big[B_x(x,y) - D_x(x,y)\big]x + (1-v)B_y(x,y)y - \lambda x, \tag{101}$$

$$\dot{y} = \lambda x + vB_y(x,y)y - D_y(x,y)y. \tag{102}$$

For rare mutants, selection multiplies the birth rate $B_i$ by $1+s$, while the death rate $D_i$ is evaluated using the resident demographic parameters and is not multiplied by $1+s$. Parameter value $\phi = 0$ places all density regulation in reduced birth, whereas $\phi = 1$ makes birth density independent and places all density regulation in increased death. Intermediate values allow both processes, as observed in the related ciliate *T. pyriformis* [11]; that study motivates the model structure but is not used to estimate $\phi$ for *T. utriculariae*.

To understand the role of $\phi$ we note that the net per-capita growth rate,

$$B_i(x,y) - D_i(x,y) = r_i\left(1-\frac{x+y}{K}\right) - d_i, \tag{103}$$

is independent of $\phi$. The experiments followed aposymbiotic and symbiotic cultures separately. When only state $i$ is present, all values of $\phi$ consequently produce the same deterministic density trajectory and the same positive equilibrium density, $K\left(1-\frac{d_i}{r_i}\right)$, for population $i \in \{x, y\}$. The observed density trajectories therefore cannot determine whether density regulation acts through reduced reproduction, increased mortality, or a combination of the two.

This distinction nevertheless matters for the evolutionary calculation (see below) because births and deaths are separate stochastic events, even when their difference is unchanged. Moreover, in the two-state model, loss of the symbiont occurs during reproduction: a fraction $1-v$ of offspring produced by symbiotic hosts enters the aposymbiotic state. Changing $\phi$ can therefore change both demographic turnover and the equilibrium composition of the population. We consequently treat $\phi$ as an unknown mechanism parameter and examine representative values between 0 and 1.

*Estimating birth rates and density-regulation scale.* The separate density trajectories constrain the low-density net growth rates $r_i - d_i$ much more strongly than they constrain reproduction and mortality separately. To inform the model we performed the following steps. (1) We assumed a common baseline mortality, $d_x = d_y = d$, and used $d = 0.03$ d$^{-1}$ as a reference value to obtain the joint fit, which yielded

$$r_x = 5.9600 \text{ d}^{-1}, \qquad r_y = 1.3629 \text{ d}^{-1}, \qquad K = 9.812 \times 10^4 \text{ cells ml}^{-1}.$$

These are the values given in table 8. Conditional intervals associated with this choice are described below. (2) We also explored how the model behavior changes as we vary $d$. To do this, we calculated the low-density net growth rates, $g_x = r_x - d = 5.9300$ d$^{-1}$ and $g_y = r_y - d = 1.3329$ d$^{-1}$. We then varied $d$ while preserving these fitted low-density net growth rates. Thus, for each value of $d$, we set

$$r_x(d) = g_x + d, \qquad r_y(d) = g_y + d. \tag{104}$$

These are the dependencies used below.

*Environmental acquisition.* To estimate the parameter $\lambda$, we used the fractions of ciliates that were symbiotic after 44 days—not per-day acquisition probabilities. Šimek *et al.* reported that approximately 80% of bacteria-fed ciliates and 50% of ciliates supplied with algae alone had reacquired symbionts after 44 days [8]. Applying the endpoint relation

$$\lambda = -\frac{\log(1 - f_{44})}{44} \tag{105}$$

gives $\lambda = 0.0366$ d$^{-1}$ and $0.0158$ d$^{-1}$ for the two treatments. We use the intermediate value $\lambda = 0.025$ d$^{-1}$ as the reference environmental-acquisition rate.

| Quantity | Reference value or range | How obtained |
|---|---|---|
| $r_x$ | 5.9600 $[5.661, 6.385]$ d$^{-1}$ | Joint conditional fit |
| $r_y$ | 1.3629 $[1.297, 1.439]$ d$^{-1}$ | Joint conditional fit |
| $d_x, d_y$ | $d_x = d_y = 0.03$ d$^{-1}$; also varied in $\log_{10} d \in [-3, -1]$ | Assumed/sensitivity tested |
| $K$ | $9.812 \times 10^4$ $[8.60 \times 10^4, 1.10 \times 10^5]$ cells ml$^{-1}$ | Joint conditional fit |
| $\lambda$ | 0.025 d$^{-1}$; endpoint range 0.0158–0.0366 | Calculated from reinfection |
| $v$ | No point estimate; vary $0 \le 1 - v \le 1$ | Not measured |
| $\phi$ | $0 \le \phi \le 1$; representative values $0, 0.5, 1$ | Not measured |

Table 8: Parameters for the *Tetrahymena utriculariae–Micractinium tetrahymenae* analysis. Brackets are 95% bootstrap intervals conditional on $d_x = d_y = 0.03$ d$^{-1}$ and on the fitting procedure described in the text.

**Uncertainty and limitations.** The conditional intervals for $r_x$, $r_y$, and $K$ were obtained by drawing 3,000 nonparametric bootstrap samples from the data provided in [10]. Complete replicate trajectories were resampled within each of the four state–light groups, and every resampled data set was refitted using the procedure above. These intervals are therefore conditional on $d = 0.03$ d$^{-1}$, the days 0–2 fitting window, the common-$K$ assumption, and the log-scale error model.

The remaining uncertainties are structural. The *T. pyriformis* study supports allowing both density-dependent birth suppression and density-dependent mortality, but provides no transferable estimate of $\phi$. We therefore examine the range $0 \le \phi \le 1$, using several representative cases. Vertical-transmission fidelity was not measured. The value of $\lambda$ is an effective endpoint-derived scale that only provides a rough order-of magnitude estimate. Finally, $K$ is a laboratory density-regulation parameter rather than an estimate of effective population size in nature. It however does not enter the calculation of $\mathcal{R}_N$ or $\mathcal{R}_s$ below.

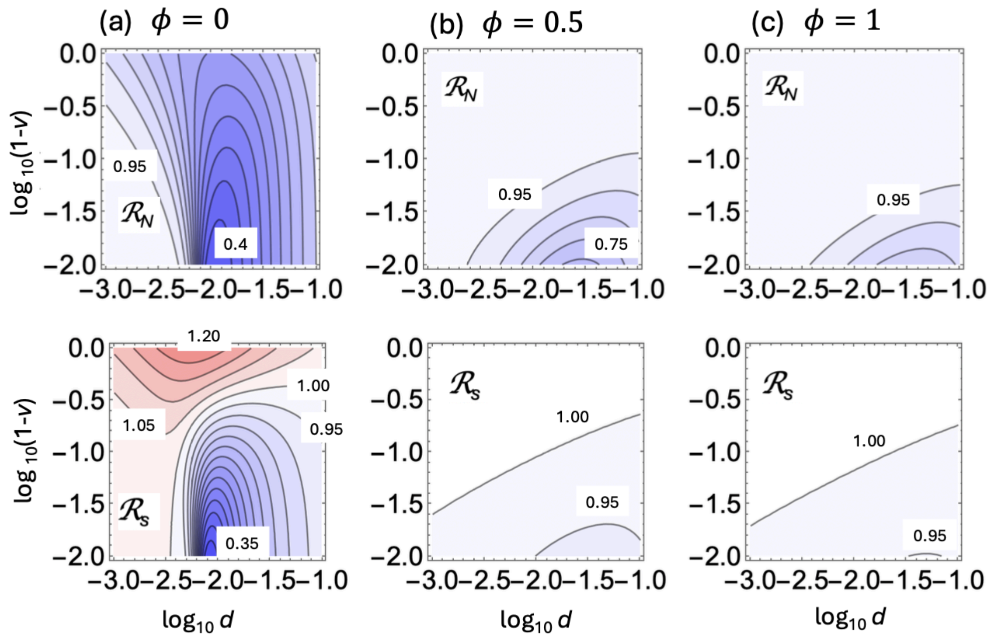

Figure 7: Tetrahymena: selection modifiers. Quantities $\mathcal{R}_N$ (top row) and $\mathcal{R}_s$ (bottom row) are plotted as functions of $\log_{10}(1-v)$ and $\log_{10} d$ (for reference, the bottom side of the graphs corresponds to high vertical transmission ($v \to 1$)). Red (blue) shading corresponds to values $> 1$ ($< 1$). (a) Only births are density dependent ($\phi = 0$), (b) both births and deaths are density-dependent ($\phi = 0.5$), (c) only deaths are density-dependent ($\phi = 1$). The step-size in the contour values is $\Delta = 0.05$ in all panels. The rest of the parameters are given in table 8 (central values).

**Effect on evolution.** To study the evolutionary consequences of this symbiont, in figure 7 we plotted the modifiers $\mathcal{R}_N$ (top row) and $\mathcal{R}_s$ (bottom row), varying the (unknown) strength of vertical transmission $v$, and the death rate, $d_x = d_y = d$ (using equation (104)). To explore the effect of density dependence, we assumed $\phi = 0$ (only reproduction is density-dependent) in panel (a), $\phi = 0.5$ (both reproduction and death are density-dependent) in panel (b), and $\phi = 1$ (only death is density-dependent) in panel (c). We observe very different dependence of the evolutionary modifiers on parameters. While for $\phi = 1$ (panel (c)) the effect of the symbiont on the effective population size and selection coefficient is modest, under $\phi = 0$ (panel (a)) we observe a significant reduction in population size: $\mathcal{R}_N \approx 0.4$ for $1 - v \sim 10^{-2}$ and small death rates. For $\mathcal{R}_s$ we see both

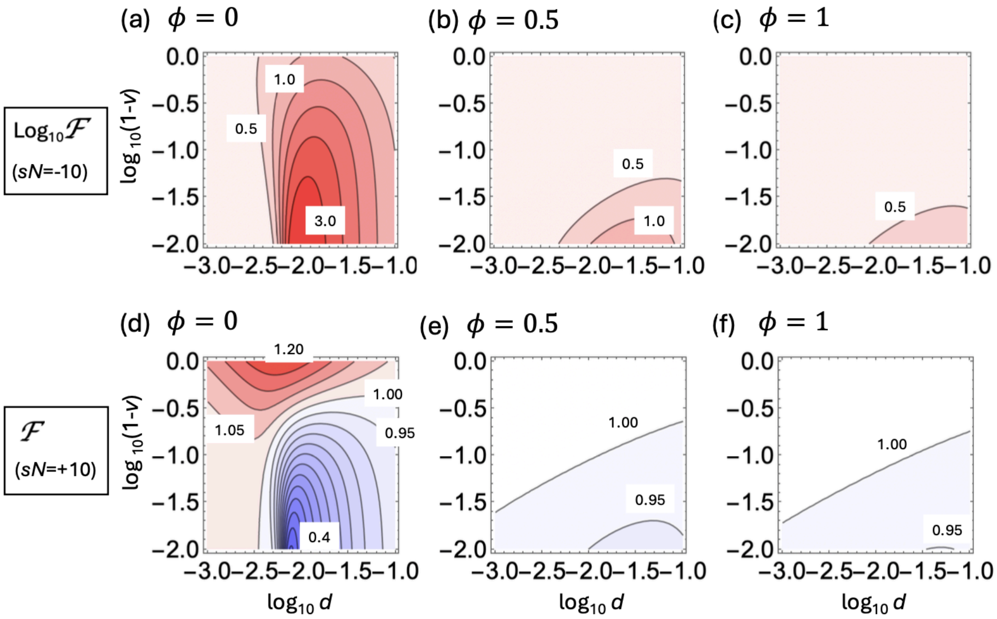

Figure 8: Tetrahymena: probability of fixation. Quantities $\log_{10}\mathcal{F}$ with $sN = -10$ (top row) and $\mathcal{F}$ with $sN = 10$ (bottom row) are plotted as functions of $\log_{10}(1-v)$ and $\log_{10} d$ (for reference, the bottom side of the graphs corresponds to high vertical transmission ($v \to 1$)). Red (blue) shading corresponds to values $> 1$ ($< 1$). (a,d) only births are density dependent ($\phi = 0$); (b,e) both births and deaths are density dependent ($\phi = 0.5$); (c,f) only deaths are density dependent ($\phi = 1$). The step-size in the contour values is $\Delta = 0.5$ in the top panels and $\Delta = 0.05$ in the bottom panels. The rest of the parameters are given in table 8 (central values).

a decrease ($\mathcal{R}_s \approx 0.35$ in the same parameter region) and an increase ($\mathcal{R}_s \approx 1.2$) for larger death rates.

Figure 8 shows how these values of the modifiers translate into changes in mutant fixation probability. The top row of panels plots $\log_{10}\mathcal{F}$ for disadvantageous mutants (fixing $sN = -10$). The bottom row plots $\mathcal{F}$ for advantageous mutants ($sN = 10$). Again, the difference between density-regulated births (panels (a) and (d)) and density-dependent deaths (panels (c) and (f)) is striking. In the latter case ($\phi = 1$), the effect of the symbiont on fixation probability is a modest suppression of selection. In the case of density-dependent births ($\phi = 0$), we observe a many-fold increase in disadvantageous mutant fixation probability, especially around $1 - v \sim 10^{-2}$ and small death rates, where it can exceed a factor of $10^3$ (panel (a)). For advantageous mutants we observe a 60% decrease in fixation probability in the same parameter region, and a 20% increase in fixation probability for larger death rates (panel (d)).

Figure 9 focuses on these two parameter regions and plots the fold-increase in fixation probability, $\mathcal{F}$, as a function of $sN$ for five values of $\phi$ from 0 to 1. Panel 9(a) represents the blue region in figure 8(d) (high vertical transmission, low death). We see a decrease in selection suppression for disadvantageous mutants as $\phi$ increases from 0 to 1.

Panel 9(b) represents the red region in figure 8(d) (low vertical transmission, high death). While the fold-increase $\mathcal{F}$ is lower for disadvantageous mutants compared to panel (a) (that is, a lower degree of selection suppression), we observe a slight amplification of selection for advantageous mutants. Again, the effects decrease as soon as density-dependence of births decreases.

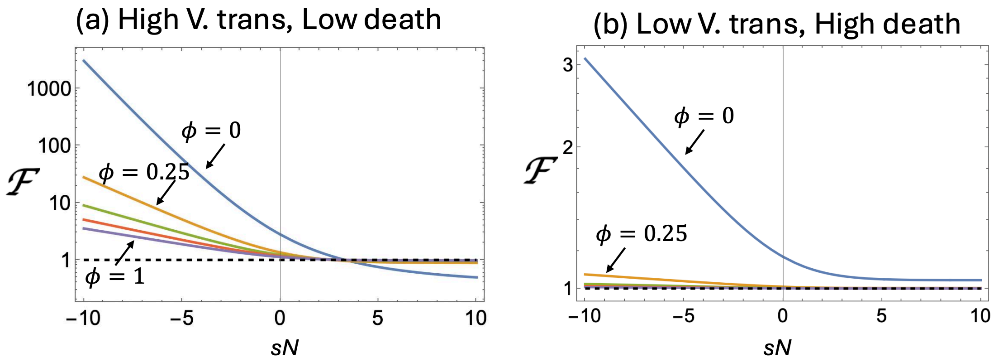

Figure 9: Tetrahymena: the ratio $\mathcal{F}$ is plotted as a function of $sN$ under fixed values of $v$ and $d$: (a) $v = 0.99, d = 0.01$; (b) $v = 0.01, d = 0.1$. The different curves correspond to five different values of $\phi$: $0, 0.25, 0.5, 0.75$ and 1. The rest of the parameters are given in table 8 (central values).

##### 3.4 *Aphis fabae–Hamiltonella defensa*

**Biological system and experiments.** The black bean aphid *Aphis fabae* can carry the facultative bacterial endosymbiont *Hamiltonella defensa*, which is transmitted maternally and can protect aphids against parasitoid wasps. This protection is strongly context dependent: in the absence of parasitoids, *H. defensa* commonly reduces aphid lifespan and lifetime reproduction, whereas some isolates provide substantial protection against the parasitoid *Lysiphlebus fabarum* [12–15]. The protection experiment most directly relevant here compared two otherwise genetically identical aphid lines: clone 405, which lacked *H. defensa*, and clone 405H76, which carried the protective H76 isolate. After a single observed parasitoid attack, 38% of uninfected aphids but only 4% of infected aphids were successfully parasitized and mummified [13]. Other experiments measured the demographic cost of infection in naturally infected, cured and reinfected aphid lines, the fidelity of maternal transmission, and the possibility of parasitoid-mediated horizontal transfer [15–18].

**Model and parameterization.** Let $x$ and $y$ denote the densities of aphids without and with a functional *H. defensa* association, respectively. We use

$$\dot{x} = r_x x \left(1 - \frac{x+y}{K}\right) + (1-v) r_y y \left(1 - \frac{x+y}{K}\right) - D_x(a)x - \beta xy, \tag{106}$$

$$\dot{y} = \beta xy + v r_y y \left(1 - \frac{x+y}{K}\right) - D_y(a)y. \tag{107}$$

Here $r_x$ and $r_y$ are the reproductive rates of uninfected and infected aphids, $v$ is the probability of maternal transmission, and $\beta$ is the effective horizontal-transmission coefficient. In this model, horizontal acquisition is represented by the term $\beta xy$ because the experimentally demonstrated route involves transfer from infected to uninfected aphids: parasitoids can acquire *H. defensa* while probing an infected aphid and introduce it into an uninfected aphid during a subsequent attack, thereby establishing a heritable infection [16, 18]. Thus, the acquisition rate depends on the availability of infected donors. Finally, parasitoid exposure is represented by an effective attack rate $a$; $q_x$ and $q_y$ are the probabilities that a single attack successfully parasitizes an uninfected or infected aphid, so parasitoids add the mortality hazards $q_x a$ and $q_y a$. In system (106-107), we denoted the total death rates as $D_x(a) = d_x + q_x a$ and $D_y(a) = d_y + q_y a$, where $d_x$ and $d_y$ are the death rates of the two populations in the absence of parasitoids.

Equations (106-107) were not fitted jointly to a single data set. Instead, their coefficients were calibrated from separate experiments. Mean lifespans of 39.6 d for cured aphids (that is, aphids from which *H. defensa* was experimentally eliminated using antibiotics) and 22.7 d for naturally infected aphids were converted to constant mortality hazards, giving $d_x = 1/39.6$ and $d_y = 1/22.7$ [15]. The uninfected reproductive rate was obtained from the reported intrinsic rate of increase, $r_m = 0.359$ d$^{-1}$, using the age-independent approximation $r_m = r_x - d_x$ [19].

The experiments indicate that the principal cost of carrying *H. defensa* in the absence of parasitoids is shortened lifespan, whereas its direct effect on reproduction rate is more modest. We therefore represent most of this cost through $d_y > d_x$ and use $r_y = 0.95r_x$ as an explicit modeling assumption for a modest reduction in reproductive rate. Maternal-transmission fidelity was set to 597/601 using the large-sample estimate available for the related aphid *A. craccivora* [17], and $q_x$ and $q_y$ were set to the reported mummification proportions following one controlled attack [13]. Host density was scaled so that $K = 1$; the two coefficients not identified by these experiments, $a$ and $\beta$, were varied over the ranges given in Table 9.

| Parameter | Value used | Source or calculation |
|---|---|---|
| $r_x,\ r_y$ | 0.384, 0.365 d$^{-1}$ | $r_x = r_m + d_x$; $r_y = 0.95r_x$ |
| $d_x,\ d_y$ | 0.0253, 0.0441 d$^{-1}$ | reciprocals of mean lifespans |
| $v$ | 0.993 | 597/601 maternally infected offspring |
| $q_x,\ q_y$ | 0.38, 0.04 | mummification after one attack |
| $a$ | 0–1 d$^{-1}$ | varied; not estimated from field data |
| $\beta$ | 0–0.10 d$^{-1}$ | varied after setting $K = 1$ |
| $K$ | 1 | normalization of host density |

Table 9: Parameters used for the *Aphis fabae*–*H. defensa* model.

**Uncertainties.** This parameterization is necessarily approximate. Aphid development, reproduction and mortality are age dependent, whereas equation (107) replaces them by constant rates; consequently, converting mean lifespan to $1/L$ and using $r_x = r_m + d_x$ are coarse-graining assumptions rather than formal maximum-likelihood fits. The value of $r_y/r_x$ is a modeling choice based on the notion that young-adult fecundity is only weakly affected (while longevity is identified as a major constitutive cost [12]). The numerical estimate of $v$ comes from *A. craccivora*, not *A. fabae*, and $q_x$ and $q_y$ apply specifically to aphid clone 405, the H76 symbiont isolate and the tested *L. fabarum* line. Moreover, a controlled single-attack outcome is only an approximation to a continuous mortality hazard, and neither the effective field attack rate $a$ nor the population-level coefficient $\beta$ is identified by the available experiments. The parameter values are therefore interpreted as an empirically anchored reference set, with $a$ and $\beta$ treated explicitly as sensitivity parameters.

**Effect on evolution.** To explore the effect of the bacterial endosymbiont on aphid evolution, we varied the attack rate, $a$, and the rate of horizontal transmission, $\beta$, neither of which could be determined from the experiments. As the attack rate, $a$, increases, there are two threshold values that are crossed. The first threshold value, which we call $a_{c,1}$, corresponds to the aphids with symbionts becoming fitter than those without. This value satisfies

$$\frac{r_y D_x(a_{c,1})}{r_x D_y(a_{c,1})} = 1,$$

and for our system it is $a_{c,1} = 0.0626$ d$^{-1}$, see figure 10(a). If $a < a_{c,1}$, the symbiont is deleterious and if $a > a_{c,1}$, it is beneficial. This transition occurs because parasitoid attack increases mortality

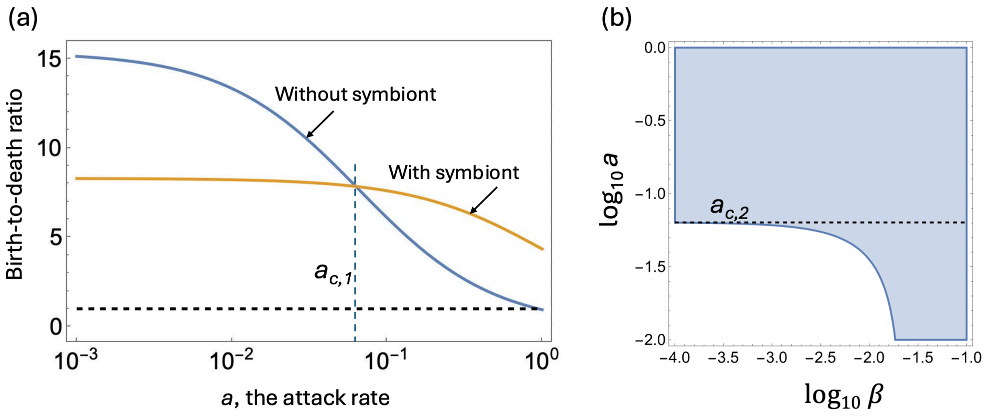

Figure 10: Aphids. (a) The birth-to-death ratios, $r_x/D_x(a)$ (blue) and $r_y/D_y(a)$ (yellow), are plotted as a function of the attack rate, $a$. The microbe becomes beneficial for $a > a_{c,1}$. (b) The region of existence of a positive solution, $(N_x, N_y)$, is plotted in the $(\log_{10}\beta, \log_{10} a)$ space. If $a > a_{c,2}$, the solution exists for any $\beta > 0$. If $a < a_{c,2}$, there is a positive threshold value, $\beta_c$. The rest of the parameters are as in table 9.

much more rapidly in aphids without the symbiont than in those carrying it. Above $a_{c,1}$, the protection against parasitoid-induced mortality outweighs the constitutive demographic cost of carrying the symbiont.

The second threshold, $a_{c,2}$, determines whether the symbiont-associated compartment can persist without horizontal transmission. At the equilibrium containing only unassociated aphids, $N_x^{(0)} = K[1 - D_x(a)/r_x]$, the per-capita growth rate of a rare symbiont-associated compartment is

$$\beta N_x^{(0)} + v r_y \frac{D_x(a)}{r_x} - D_y(a).$$

Consequently, if $a < a_{c,2}$, where $a_{c,2}$ satisfies

$$\frac{v r_y D_x(a_{c,2})}{r_x D_y(a_{c,2})} = 1,$$

vertical transmission alone is insufficient to maintain the symbiont, and coexistence requires

$$\beta > \beta_c(a) = \frac{D_y(a) - v r_y D_x(a)/r_x}{N_x^{(0)}} > 0.$$

As $a$ increases, parasitoid-induced mortality rises much faster in unassociated than in symbiont-associated aphids, reducing the amount of horizontal transmission required for persistence. At $a = a_{c,2}$, $\beta_c = 0$; for $a > a_{c,2}$, the protection supplied by the symbiont is sufficient for the associated compartment to persist through vertical transmission alone, so a positive coexistence equilibrium exists for every $\beta \geq 0$. For the parameters in table 9, we have $a_{c,2} = 0.06359$ d$^{-1}$, which is only slightly larger than $a_{c,1}$. Figure 10(b) shows the region of existence for the positive $(N_x, N_y)$ solution.

For each fixed attack rate $a > a_{c,2}$, let $(N_x(\beta; a), N_y(\beta; a))$ be the positive coexistence equilibrium and denote $q(\beta; a) = \frac{N_y(\beta;a)}{N_x(\beta;a)}$. Let us then take the limit $\beta \to 0$ along this equilibrium branch. In this limit,

$$q(\beta; a) \longrightarrow q_0(a) = \frac{v r_y D_x(a) - r_x D_y(a)}{(1 - v) r_y D_y(a)} > 0,$$

whereas

$$\rho_y(\beta;a) = \frac{(1-v)b_y}{\beta N_x(\beta;a)} \longrightarrow \infty.$$

Substitution into the general expressions for $\mathcal{R}_N$ and $\mathcal{R}_s$ gives

$$\lim_{\beta\to 0} \mathcal{R}_N(a,\beta) = \frac{r_x + r_y q_0(a)}{v r_y[1+q_0(a)]}, \qquad \lim_{\beta\to 0} \mathcal{R}_s(a,\beta) = \frac{v r_y q_0(a)}{r_x + r_y q_0(a)}.$$

If, after taking this fixed-$a$ limit, we subsequently let $a \to a_{c,2}$ from above, then $q_0(a) \to 0$, and hence

$$\lim_{a\downarrow a_{c,2}} \lim_{\beta\downarrow 0} \mathcal{R}_N(a,\beta) = \frac{r_x}{r_y v} \simeq 1.0597, \qquad \lim_{a\downarrow a_{c,2}} \lim_{\beta\downarrow 0} \mathcal{R}_s(a,\beta) = 0.$$

Note that these are iterated limits, not a joint limit as $(a,\beta) \to (a_{c,2}, 0)$; the joint limit is singular and depends on the path of approach. Thus $\mathcal{R}_s$ can be made arbitrarily small by choosing $a > a_{c,2}$ sufficiently close to $a_{c,2}$ and then taking $\beta$ sufficiently small, but it does not approach zero as $\beta \downarrow 0$ for an arbitrary fixed $a > a_{c,2}$. The arbitrarily strong attenuation of the effective selection coefficient therefore results from the non-uniformity of the limit $\beta \downarrow 0$ as $a \downarrow a_{c,2}$: the limiting value depends on the path by which this corner $(a,\beta) = (a_{c,2}, 0)$ is approached.

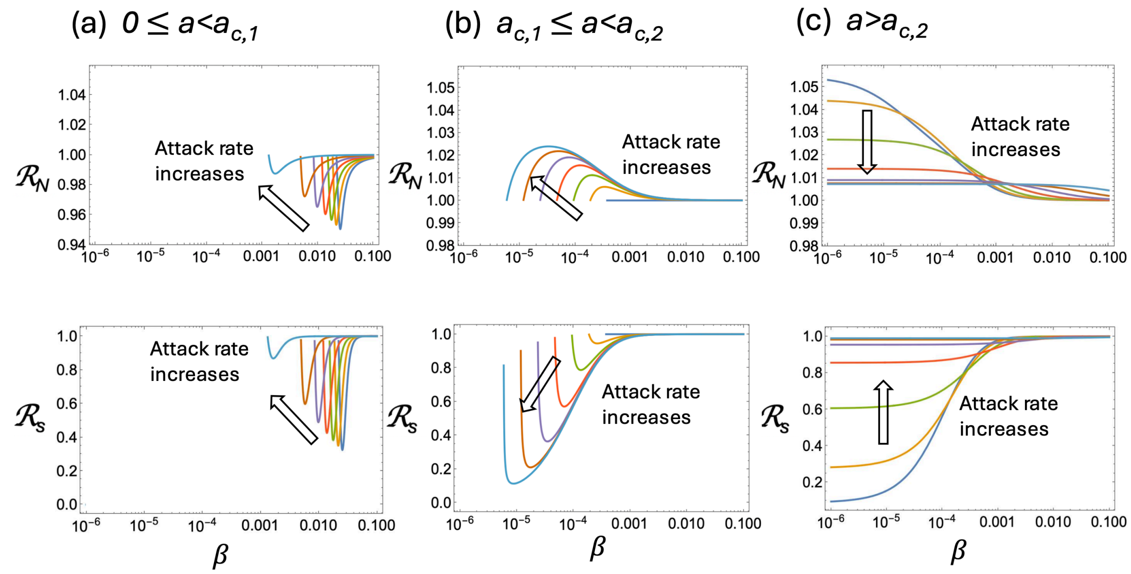

Figure 11: Aphids: the effect of the endosymbiont on the modifiers $\mathcal{R}_N$ (the top row) and $\mathcal{R}_s$ (the bottom row). The modifiers are plotted as functions of $\beta$, and each curve represents a different value of the attack rate, $a$. (a) $0 \le a < a_{c,1}$. The values of $a \in \{0.01k\}_{k=0}^{6}$. (b) $a_{c,1} \le a < a_{c,2}$ with $a \in \{a_{c,1} + \frac{(2^k-1)a_{c,2}}{2^k}\}_{k=0}^{6}$. (c) $a > a_{c,2}$ with $a \in \{a_{c,2} + 10^{-4} \times 4^k\}_{k=0}^{6}$. The direction of $a$ increase is indicated in each panel. Please note the same horizontal and vertical range for panels (a-c). The rest of the parameters are as in table 9.

Figure 11 studies the behavior of the modifiers $\mathcal{R}_N$ (top row) and $\mathcal{R}_s$ (bottom row) for different values of the attack rate, $a$, and for a range of the horizontal transmission rate, $\beta$. Because there are two thresholds for the attack rate, the three regions of $a$ are shown. In panel (a), we explore the range of $a$ where the symbiont makes the host disadvantageous: $0 \le a \le a_{c,1}$. Seven different values of $a$ are presented. As $a$ increases, the lower bound of $\beta$ for which a nonzero $(N_x, N_y)$

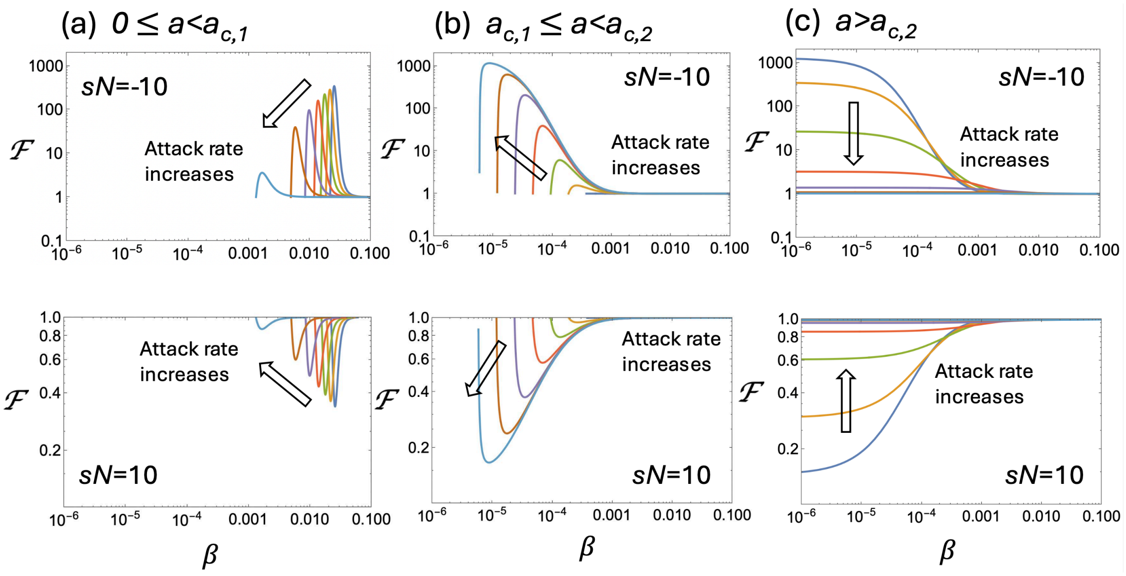

Figure 12: Aphids: the effect of the endosymbiont on the fixation probability for disadvantageous ($sN = -10$, the top row) and advantageous ($sN = 10$ the bottom row) mutants. function $\mathcal{F}$ is plotted as functions of $\beta$, and each curve represents a different value of the attack rate, $a$. (a) $0 \le a < a_{c,1}$. The values of $a \in \{0.01k\}_{k=0}^{6}$. (b) $a_{c,1} \le a < a_{c,2}$ with $a \in \{a_{c,1} + \frac{(2^k-1)a_{c,2}}{2^k}\}_{k=0}^{6}$. (c) $a > a_{c,2}$ with $a \in \{a_{c,2} + 10^{-4} \times 4^k\}_{k=0}^{6}$. The direction of $a$ increase is indicated in each panel. Please note the same horizontal and vertical range for panels (a-c). The rest of the parameters are as in table 9.

solution exists, becomes lower (see figure 10(b)). The values of $\mathcal{R}_N$ and $\mathcal{R}_s$ are both less than one, nonmonotonic with $\beta$, and experience a minimum for $a = 0$.

Panel (b) shows the interval $a_{c,1} \le a < a_{c,2}$, where the symbiont becomes beneficial, but there is still a nonzero lower bound of $\beta$ to guarantee existence of solutions. This interval is very narrow under the parameter values used here, so these panels mostly demonstrate extreme sensitivity of the modifiers to small changes of the attack in this region. In this region, the effect on $\mathcal{R}_N$ changes sign (but remains very small), while $\mathcal{R}_s$ continues to be smaller than one, but the effect increases significantly.

Finally, we study $a > a_{c,2}$ in panel 11(c). While $\mathcal{R}_N$ continues to be $> 1$ in this region, the magnitude of the effect on the effective population size decreases further. $\mathcal{R}_N$ has a limiting value at $a = a_{c,2}$ and $\beta \to 0$, given by $r_x/(r_y v) \approx 1.0597$. The effect on $\mathcal{R}_s$ is much more significant. As shown by the iterated limit above, $\mathcal{R}_s$ can be made arbitrarily small by first choosing $a > a_{c,2}$ sufficiently close to $a_{c,2}$ and then taking $\beta$ sufficiently small. It is the decrease in the effective selection coefficient that leads to very strong selection suppression in this system, as we show next.

Figure 12 explores the fold change in fixation probability, $\mathcal{F}$, for disadvantageous and advantageous mutants, using $sN = -10$ in the top row and $sN = 10$ in the bottom row. The three panels correspond to the three panels in figure 11. We observe that selection is always suppressed in these cases (that is, $\mathcal{F} > 1$ for disadvantageous and $\mathcal{F} < 1$ for advantageous mutants). The effect can be significant for the case of zero and low attack rate, where the fold-increase in the probability of fixation for disadvantageous mutants can reach about $10^2$, and the probability of advantageous mutant fixation can be decreased by about 60%. The effect is the strongest for the case where the symbiont is advantageous (and $a$ is around $a_{c,2}$).

##### 3.5 *Agrostis hyemalis–Epichloë amarillans*

**Biological system and experiments.** *Agrostis hyemalis* (winter bentgrass) is a short-lived perennial grass that can carry the facultative fungal endophyte *Epichloë amarillans.* The fungus is transmitted principally from an associated plant to its seeds, although transmission is imperfect and horizontal acquisition is biologically possible. Donald *et al.* established replicated field populations with initial endophyte prevalences of 5, 35, 65, or 95% and exposed them to either ambient or experimentally elevated precipitation [20]. They assayed 5,485 plants for endophyte status and followed 954 individual plants to estimate survival, flowering, seed production, recruitment, and vertical transmission.

The use of several initial prevalences allowed the authors to estimate how prevalence in one annual census predicts prevalence in the next. The relationship differed between the two annual transitions studied. During the 2015–2016 transition, the fitted prevalence map crossed the line of no change at an intermediate prevalence under both precipitation treatments: prevalence tended to increase below this point and decrease above it. These stable interior fixed points provide the equilibrium associated-host fractions used below. We use this transition, rather than the qualitatively different 2014–2015 transition, because it supports positive coexistence of endophyte-free and endophyte-associated plants in both environments.

**Model and parameterization.** Let $x(t)$ and $y(t)$ denote the densities of endophyte-free and endophyte-associated plants. The model is

$$\dot{x} = r_x x\left(1 - \frac{x+y}{K}\right) + (1-v)r_y y\left(1 - \frac{x+y}{K}\right) - dx - \beta xy, \tag{108}$$

$$\dot{y} = \beta xy + v r_y y\left(1 - \frac{x+y}{K}\right) - dy, \tag{109}$$

where $r_x$ and $r_y$ are the reproduction rates of endophyte-free and endophyte-associated plants, $d$ is their common mortality rate, $v$ is vertical-transmission fidelity, and $\beta$ is the horizontal-transmission coefficient. If $N_x$ and $N_y$ are the positive equilibrium densities, then $N = N_x + N_y$ is the total equilibrium population and

$$f^* = \frac{N_y}{N} \tag{110}$$

is the equilibrium fraction of endophyte-associated plants.

Donald *et al.* [20] did not fit the continuous-time rates $r_x$ and $r_y$. Instead, their hierarchical Bayesian model estimated the annual life-history components that determine recruitment. Let $F_i$ be the probability that a plant in state $i \in \{x, y\}$ flowers, $S_i$ its seed production conditional on flowering, and $G_x$ and $G_y$ the recruitment probabilities of endophyte-free and endophyte-associated seeds. Collapsing these components into the expected number of successful recruits per plant gives

$$\widetilde{r}_x = F_x S_x G_x, \qquad \widetilde{r}_y = F_y S_y\big[vG_y + (1-v)G_x\big]. \tag{111}$$

Here $S_i$ incorporates the fitted inflorescence production, seed mass per inflorescence, and the empirical conversion from seed mass to seed number. The quantities $\widetilde{r}_x$ and $\widetilde{r}_y$ are annual, life-cycle-integrated recruitment outputs rather than continuous-time reproduction rates. To translate these estimates into the continuous-time model, we assume that their ratio is preserved: $\frac{r_y}{r_x} = \frac{\widetilde{r}_y}{\widetilde{r}_x}$. This assumption determines the relative reproduction rates but not their absolute continuous-time scale.

Using the central estimates for 2015–2016 gives

$$\widetilde{r}_x \simeq 1.99, \qquad \widetilde{r}_y \simeq 2.52, \qquad \frac{r_y}{r_x} = \frac{\widetilde{r}_y}{\widetilde{r}_x} = \frac{2.52}{1.99} \simeq 1.27 \qquad \text{(ambient)}, \tag{112}$$

$$\widetilde{r}_x \simeq 1.59, \qquad \widetilde{r}_y \simeq 2.21, \qquad \frac{r_y}{r_x} = \frac{\widetilde{r}_y}{\widetilde{r}_x} = \frac{2.21}{1.59} \simeq 1.39 \qquad \text{(elevated precipitation)}. \tag{113}$$

The quantities $\widetilde{r}_x$ and $\widetilde{r}_y$ are annual, life-cycle-integrated recruitment outputs, and not continuous-time rates. Only their dimensionless ratio is used in the model, so the arbitrary time scale of $r_x$ and $r_y$ is immaterial.

The equilibrium fraction $f^*$ was obtained as the stable fixed point of the fitted 2015–2016 prevalence map, and $v$ was estimated from the fraction of seeds that retained the endophyte. The fitted survival effect of the endophyte was centred near zero, motivating the equal-mortality assumption $d_x = d_y = d$. The empirical inputs required for the calculation are summarized in Table 10.

| Precipitation treatment | $r_y/r_x$ | $v$ | $f^*$ |
|---|---|---|---|
| Ambient | $2.52/1.99 = 1.27$ | $0.883$ $[0.806, 0.938]$ | $0.720$ $[0.635, 0.795]$ |
| Elevated | $2.21/1.59 = 1.39$ | $0.909$ $[0.842, 0.956]$ | $0.840$ $[0.755, 0.937]$ |

Table 10: Empirical inputs for the *Agrostis hyemalis*–*Epichloë amarillans* model. Brackets are 95% posterior intervals reported for vertical-transmission fidelity $v$ and equilibrium endophyte prevalence $f^*$.

**Uncertainties.** Donald *et al.* estimated the prevalence and vital-rate components jointly in a hierarchical Bayesian model. Unobserved endophyte states were treated as latent variables, population-level random effects represented variation among field populations, and posterior predictive checks were used to assess model adequacy. The intervals for $v$ and $f^*$ therefore describe uncertainty in the corresponding transmission and prevalence submodels.

No posterior interval was reported for the composite ratio $r_y/r_x$. The values 1.27 and 1.39 combine central estimates of several life-history components; a valid uncertainty interval would require their joint posterior draws. Accordingly, the reported $\mathcal{R}_N$ and $\mathcal{R}_s$ values are point calculations rather than posterior estimates. Moreover, $f^*$ is the fixed point of a prevalence relationship inferred from one annual transition, not a population observed at equilibrium for many generations. Its interpretation assumes that the ecological conditions and prevalence map of 2015–2016 persist.

**Effect on evolution.** At a first glance, there is a lot of uncertainty in the system, as table 10 does not contain separate measurements for the reproduction rates, does not list the death rates or the horizontal-transmission coefficient, $\beta$. It turns out however that the knowledge of $f^*$ removes a lot of uncertainty, at least if our purpose is to calculate $\mathcal{R}_N$ and $\mathcal{R}_s$. Below we show that if we assume that $d_x = d_y = d$, then the knowledge of specific values of $d$ and $\beta$ is not required to calculate $\mathcal{R}_N$ and $\mathcal{R}_s$.

Define as before,

$$q = \frac{N_y}{N_x} = \frac{f^*}{1 - f^*}, \qquad \gamma = \frac{r_x}{r_y}, \qquad \ell = 1 - \frac{N_x + N_y}{K}. \tag{114}$$

At a positive coexistence equilibrium, dividing the equilibrium equations for $x$ and $y$ by $N_x$ and

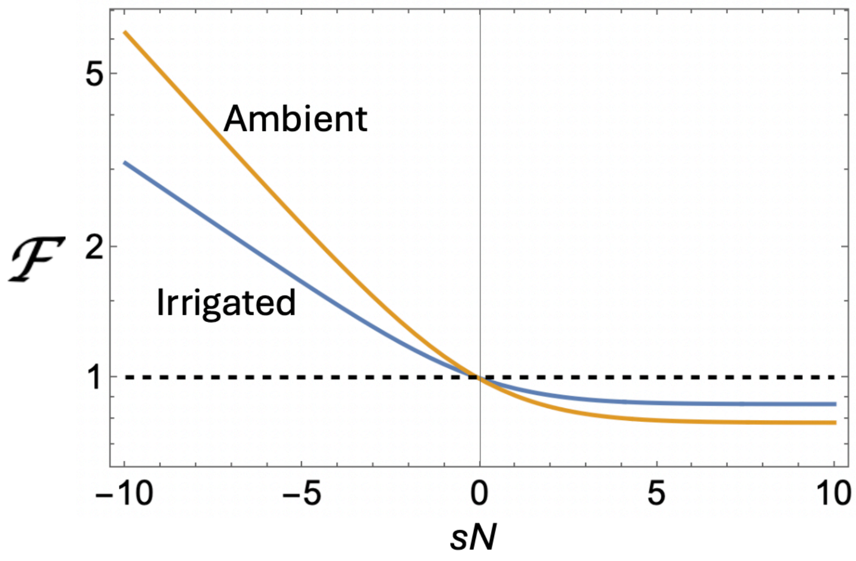

Figure 13: Agrostis: the effect of the fungal endophyte on the fixation probability. The fold-change $\mathcal{F}$ is plotted as a function of $sN$ for the ambient and irrigated conditions. Parameter values are given in table 10 (central values).

$N_y$, respectively, gives

$$d = r_x \ell + (1-v) r_y q \ell - \beta q N_x, \tag{115}$$
$$d = \beta N_x + v r_y \ell. \tag{116}$$

Eliminating $d$ between these equations yields

$$\frac{\beta N_x}{r_y \ell} = \frac{\gamma - v + (1-v)q}{1+q}. \tag{117}$$

The right-hand side is fixed by the empirically specified values of $r_y/r_x = 1/\gamma$, $v$, and $f^*$. It follows that the reproductive value of a mutant in state $y$, relative to that of a mutant in state $x$, is

$$\rho_y = \frac{(1-v) r_y \ell}{\beta N_x} = \frac{(1-v)(1+q)}{\gamma - v + (1-v)q}. \tag{118}$$

The probabilities that a de novo mutation arises in the two states are

$$P_x^{\text{de novo}} = \frac{\gamma + (1-v)q}{\gamma + q}, \qquad P_y^{\text{de novo}} = \frac{vq}{\gamma + q}. \tag{119}$$

Therefore, defining

$$h = (1-v) + v\rho_y, \tag{120}$$

the general expressions for the two evolutionary modifiers become

$$\mathcal{R}_N = \frac{1 + \rho_y q}{(1+q)\left(P_x^{\text{de novo}} + \rho_y P_y^{\text{de novo}}\right)}, \tag{121}$$
$$\mathcal{R}_s = \frac{(\gamma + qh)^2}{(\gamma + q)(\gamma + \rho_y q h)}. \tag{122}$$

These expressions contain neither $d$ nor $\beta$. Consequently, for fixed $r_y/r_x$, $v$, and $f^*$, every admissible pair $(d, \beta)$ that produces the specified positive, locally stable coexistence equilibrium gives the same values of $\mathcal{R}_N$ and $\mathcal{R}_s$. The central parameter values in Table 10 give

$$(\mathcal{R}_N, \mathcal{R}_s) = (1.027, 0.900) \tag{123}$$

under ambient precipitation and

$$(\mathcal{R}_N, \mathcal{R}_s) = (1.022, 0.932) \tag{124}$$

under elevated precipitation.

This invariance requires fixed $r_y/r_x$, $v$, and $f^*$, as well as equal mortality rates in the two states. Although $d$ and $\beta$ may affect the total equilibrium population $N$, and therefore the absolute effective population size $N_{\text{eff}} = N\mathcal{R}_N$, they do not affect either dimensionless modifier under these assumptions.

Figure 13 shows the fixation-probability ratio $\mathcal{F}$ as a function of $Ns$ under ambient and elevated precipitation. Both treatments produce modest suppression of selection over the range shown: relative to the unstructured model, $\mathcal{F} > 1$ for deleterious mutations and $\mathcal{F} < 1$ for advantageous mutations.

### 4 More complex microbiome states

Here we develop a 4-compartment model that contains two types of microbes and three microbial states. Let us denote by $y_0$ the number of organisms that are not associated with any microbiome; by $y_i$ with $i \in \{1, 2\}$ the number of organisms that are associated with microbe $i$ only, and by $y_{12}$ the number of organisms that are associated with both microbes 1 and 2. The ODEs that describe the expected dynamics are as follows:

$$\begin{aligned}
\dot{y}_0 &= [r_0 y_0 + (1 - v_1) r_1 y_1 + (1 - v_2) r_2 y_2 + (1 - v_1)(1 - v_2) r_{12} y_{12}]\,\ell \\
&\quad - d_0 y_0 - y_0(\lambda_1 + \lambda_2), && (125)\\
\dot{y}_1 &= \lambda_1 y_0 + [v_1 r_1 y_1 + v_1(1 - v_2) r_{12} y_{12}]\,\ell - d_1 y_1 - y_1 \lambda_2, && (126)\\
\dot{y}_2 &= \lambda_2 y_0 + [v_2 r_2 y_2 + (1 - v_1) v_2 r_{12} y_{12}]\,\ell - d_2 y_2 - y_2 \lambda_1, && (127)\\
\dot{y}_{12} &= \lambda_2 y_1 + \lambda_1 y_2 + v_1 v_2 r_{12} y_{12} \ell - d_{12} y_{12}, && (128)
\end{aligned}$$

where we denoted the density regulation term,

$$\ell = 1 - \frac{y_0 + y_1 + y_2 + y_{12}}{K}.$$

The dynamics of rare mutant hosts is described by the events in table 11.

Hosts can now occupy four states: carrying neither microbe, carrying either microbe alone, or carrying both. Each state can have its own reproduction and mortality rates, and each microbe can be acquired environmentally and transmitted vertically with its own fidelity. Consequently, reproduction of a doubly colonized host can produce offspring retaining neither, either, or both microbes.

We illustrate this extension using the *Drosophila melanogaster* system considered above. We selected two bacteria with contrasting demographic effects on the host: *Lactobacillus brevis* (Lb), which is deleterious, and *Acetobacter orientalis* (Ao), which is beneficial. The demographic rates of germ-free, Lb-associated, and Ao-associated flies are anchored by the single-species measurements

| Description | Rates, $\boldsymbol{c}_r(s)^\mathsf{T}$ | Change, $\boldsymbol{\nu}^\mathsf{T}$ |
|---|---|---|
| Increase in $y_0$ by reproduction | $(1+s)\ell \times (r_0, (1-v_1)r_1, (1-v_2)r_2, (1-v_1)(1-v_2)r_{12})$ | $(1,0,0,0)$ |
| Increase in $y_1$ by reproduction | $(1+s)\ell \times (0, v_1 r_1, 0, v_1(1-v_2)r_{12})$ | $(0,1,0,0)$ |
| Increase in $y_2$ by reproduction | $(1+s)\ell \times (0, 0, v_2 r_2, (1-v_1)v_2 r_{12})$ | $(0,0,1,0)$ |
| Increase in $y_{12}$ by reproduction | $(1+s)\ell \times (0,0,0, r_{12}v_1 v_2)$ | $(0,0,0,1)$ |
| Death in $y_0$ | $(d_0,0,0,0)$ | $(-1,0,0,0)$ |
| Death in $y_1$ | $(0,d_1,0,0)$ | $(0,-1,0,0)$ |
| Death in $y_2$ | $(0,0,d_2,0)$ | $(0,0,-1,0)$ |
| Death in $y_{12}$ | $(0,0,0,d_{12})$ | $(0,0,0,-1)$ |
| Acquisition of 1st microbe by $y_0$ | $(\lambda_1,0,0,0)$ | $(-1,1,0,0)$ |
| Acquisition of 2nd microbe by $y_0$ | $(\lambda_2,0,0,0)$ | $(-1,0,1,0)$ |
| Acquisition of 1st microbe by $y_2$ | $(0,0,\lambda_1,0)$ | $(0,0,-1,1)$ |
| Acquisition of 2nd microbe by $y_1$ | $(0,\lambda_2,0,0)$ | $(0,-1,0,1)$ |

Table 11: Ecological processes associated with the dynamics of hosts in the presence of two microbes. Events and rates for rare-mutant populations are shown.

of Gould *et al.* [7]. The principal additional uncertainty concerns flies carrying both bacteria simultaneously. Rather than assigning a single, poorly supported demographic phenotype to this state, we therefore examined a range of possibilities. Specifically, we varied the birth and death rates of doubly colonized hosts between the values estimated for Lb-associated and Ao-associated flies (Fig. 14a). This analysis is not intended as a quantitative prediction for the Lb–Ao combination. Instead, it provides a biologically anchored demonstration of the framework when multiple microbial states must be represented explicitly.

The resulting evolutionary quantities remain strongly dependent on microbiome-associated population structure across this entire range (Fig. 14b–d). For all six assumptions about the doubly colonized state shown here, $\mathcal{R}_N < 1$ and $\mathcal{R}_s < 1$: the effective population size is smaller than census size and the magnitude of selection on host mutations is reduced relative to its intrinsic value. The extent of both effects nevertheless depends substantially on the demographic properties assigned to the doubly colonized hosts. When their demography is closer to that of the deleterious Lb association, both modifiers are strongly reduced; as it approaches the beneficial Ao-associated state, $\mathcal{R}_N$ moves closer to one and the suppression of selection becomes less pronounced (Fig. 14b,c).

These changes translate into substantial differences in mutant fixation (Fig. 14d). In particular, weakened selection and enhanced drift can greatly increase the relative fixation probability of deleterious host mutations, whereas advantageous mutations can become less likely to fix than predicted from census population size and intrinsic selection alone. The quantitative magnitude depends on the uncertain properties of the doubly colonized state, but the qualitative conclusion does not: introducing additional microbial states does not remove the microbiome-mediated modification of host population genetics.

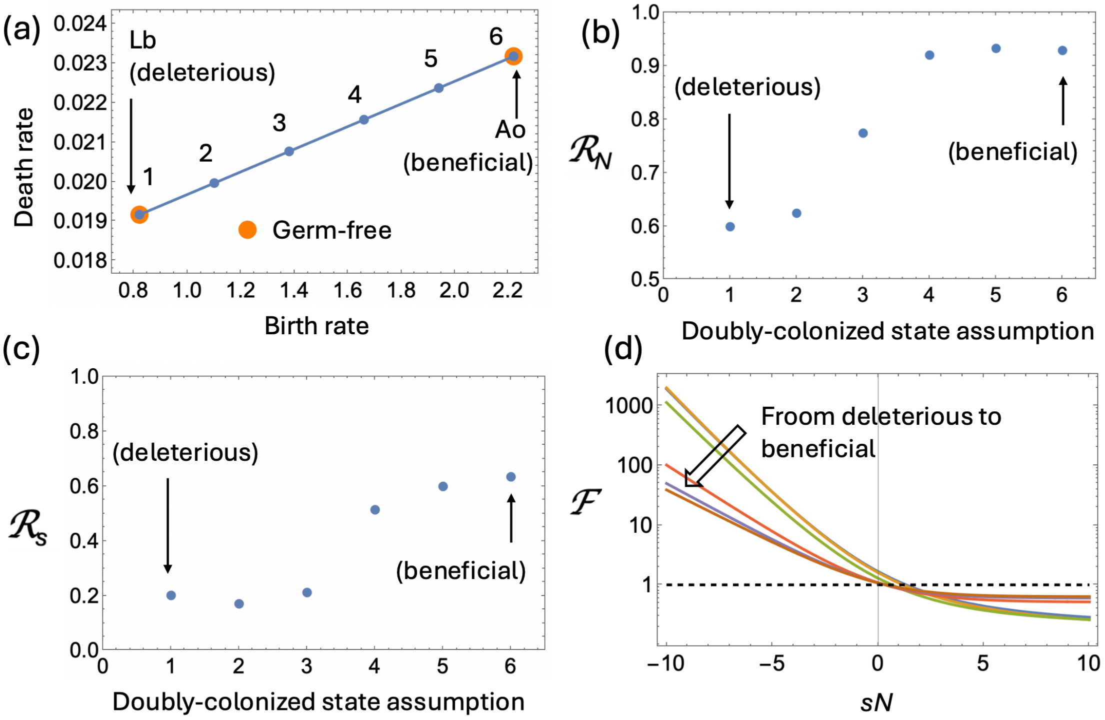

Figure 14: Evolutionary consequences of a microbiome containing two bacterial species in *Drosophila melanogaster*. The model contains four host states: germ-free, associated with *Lactobacillus brevis* (Lb) alone, associated with *Acetobacter orientalis* (Ao) alone, and associated with both bacteria. (a) Estimated birth and death rates of germ-free, Lb-associated, and Ao-associated flies are shown by orange circles. Lb is deleterious and Ao beneficial according to the demographic criterion used here. Because the demographic rates of the doubly colonized state are uncertain, six illustrative assumptions (1–6) span the range between the Lb- and Ao-associated states. (b,c) Effective population-size and selection modifiers, $\mathcal{R}_N$ and $\mathcal{R}_s$, respectively, for each assumption about the doubly colonized state. (d) The resulting fold change in fixation probability, $\mathcal{F} = P_{\text{fix}}(N\mathcal{R}_N, s\mathcal{R}_s)/P_{\text{fix}}(N, s)$, as a function of the scaled intrinsic selection coefficient $sN$. The rest of the demographic parameters are given in table 7 where species 1 is Lb and species 2 is Ao. The transmission parameters are $v_1 = 0.99, v_2 = 0.8, \lambda_1 = 10^{-2}, \lambda_2 = 10^{-6}$.

[8] Karel Šimek, Gianna Pitsch, Michaela M Salcher, Dagmara Sirová, Tanja Shabarova, Lubomír Adamec, and Thomas Posch. Ecological traits of the algae-bearing tetrahymena utriculariae

(ciliophora) from traps of the aquatic carnivorous plant utricularia reflexa. *Journal of Eukaryotic Microbiology*, 64(3):336–348, 2017.
